# Phox2a defines a developmental origin of the anterolateral system in mice and humans

**DOI:** 10.1101/2020.06.10.144659

**Authors:** R. Brian Roome, Farin B. Bourojeni, Bishakha Mona, Shima Rastegar-Pouyani, Raphael Blain, Annie Dumouchel, Charleen Salesse, W. Scott Thompson, Megan Brookbank, Yorick Gitton, Lino Tessarollo, Martyn Goulding, Jane E. Johnson, Marie Kmita, Alain Chédotal, Artur Kania

**Affiliations:** Institut de Recherches Cliniques de Montréal (IRCM), Montréal, QC, H2W 1R7, Canada; Integrated Program in Neuroscience, McGill University, Montréal, QC, H3A 2B4, Canada; Department of Neuroscience, UT Southwestern Medical Center, Dallas, TX, 75390, United States; Sorbonne Université, INSERM, CNRS, Institut de la Vision, 17 Rue Moreau, Paris, 75012, France; Neural Development Section, Mouse Cancer Genetics Program, National Cancer Institute, Frederick, MD, 21702, United States; Molecular Neurobiology Laboratory, The Salk Institute for Biological Studies, La Jolla, CA, 92037, United States; Department of Pharmacology, UT Southwestern Medical Center, Dallas, TX, 75390, United States; Department of Anatomy and Cell Biology, McGill University, Montréal, QC, H3A 0C7, Canada; Division of Experimental Medicine, McGill University, Montréal, QC, H3A 2B2, Canada

## Abstract

Anterolateral system neurons relay pain, itch and temperature information from the spinal cord to pain-related brain regions, but the differentiation of these neurons and their specific contribution to pain perception remain poorly defined. Here, we show that virtually all mouse spinal neurons that embryonically express the autonomic system-associated Paired-like homeobox 2A (Phox2a) transcription factor innervate nociceptive brain targets, including the parabrachial nucleus and the thalamus. We define Phox2a anterolateral system neuron birth order, migration and differentiation, and uncover an essential role for Phox2a in the development of relay of nociceptive signals from the spinal cord to the brain. Finally, we also demonstrate that the molecular identity of Phox2a neurons is conserved in the human foetal spinal cord. The developmental expression of Phox2a as a uniting feature of anterolateral system neurons suggests a link between nociception and autonomic nervous system function.

## Introduction

In vertebrates, somatosensory information about noxious stimuli is carried from peripheral nociceptors to the brain, via spinal projection neurons collectively known as the anterolateral system (AS). Together the brain regions innervated by these interpret the transmitted signals as pain, a sensation endowed with discriminative and affective components that, respectively, convey the identity, location, and intensity of the stimulus, as well as elicit behavioural responses driven by arousal and aversion (Melzack and Casey, 1968). Anatomical, clinical and physiological studies suggest that dedicated AS channels convey these different facets of pain (Price and Dubner, 1977), but since the molecular identity of AS neurons remains unknown, insights into the functional logic of nociceptive information relay from the periphery to the brain remain limited.

The AS innervates brain regions that have distinct functions in nociception. Prominent targets include the ventroposterolateral thalamus (VPL) (Gauriau and Bernard, 2004; Willis et al., 1979), which relays somatotopically organised nociceptive information (Guilbaud et al., 1980) to the primary somatosensory cortices, and the parabrachial nucleus (pB) (Bernard et al., 1995), which is considered to mediate affective components of pain by relaying noxious information to the amygdala (Han et al., 2015), and via the medial thalamus, to the prefrontal cortex (Bourgeais et al., 2001). Clinical evidence supports the division between discriminative and affective dimensions of pain, as prefrontal lobotomy (Freeman and Watts, 1948) and insular cortex-related pain asymbolia (Berthier et al., 1988; Rubins and Friedman, 1948) result in the discriminatory nature of noxious stimuli being appreciated in the absence of the negative affect normally associated with them. The critical role of AS in relaying both discriminative and affective components of nociception to its brain targets is revealed by effects of lesions to the spinal anterolateral tract, which abolish all somatic pain without affecting light touch sensibility (Spiller and Martin, 1912).

The anatomy of AS neurons is well known in rodents, where they are found principally in laminae I and V and the lateral spinal nucleus (LSN) of the spinal dorsal horn (Davidson et al., 2010; Kitamura et al., 1993). Lamina I AS neurons have small receptive fields (Willis et al., 1974) and respond to specific classes of stimuli and their modalities (e.g.: temperature, itch, mechanical vs. thermal pain (Andrew and Craig, 2001; Craig and Serrano, 1994), which are relayed to targets thought to mediate discriminatory responses such as the VPL thalamus. Lamina V/LSN AS neurons, in contrast, have broad receptive fields, wide dynamic ranges of receptivity (Craig, 2003b), and their physiology corresponds poorly with the qualitative descriptions of pain (Craig, 2004). Based on their prominent projections to the dorsal pB (Feil and Herbert, 1995) and medial thalamus (Gauriau and Bernard, 2004), lamina V/LSN neurons likely transmit the affective and motivational dimensions of pain. These AS neuron functions are in line with substance-P receptor (NK1R)-directed AS neuron ablation resulting in analgesia; however, a precise interpretation of this experiment is obscured by NK1R expression in non-AS neurons (Cameron et al., 2015; Mantyh et al., 1997). Recently developed genetic tools have advanced our understanding of afferent pathways to AS neurons by uncovering the identity of interneurons that gate transmission of innocuous sensations to AS neurons (Duan et al., 2014; Petitjean et al., 2019), but genetically targeting AS neurons has been difficult. A recent study using the gene *Tachykinin1* (*Tac1*) demonstrated ablation of a subset of spinal interneurons and pB-innervating AS neurons (Huang et al., 2019), producing behavioural deficits consistent with the loss of supraspinal transmission of nociceptive information without affecting the function of spinal nocifensive reflexes, demonstrating that Tac1-positive neurons contribute to the AS. Despite these advances, the genes expressed selectively in AS neurons remain unknown, precluding insights into how AS neuron function contributes to the experience of pain.

Much of the diversity of spinal neurons arises from a molecular logic of developmental gene expression that is no longer apparent in the adult nervous system. Developmental gene expression has been instrumental in studying locomotor circuits of the ventral spinal cord (Arber, 2012; Goulding, 2009), and may be also useful in accessing dorsal spinal cord somatosensory circuits. Like the ventral spinal cord, the developing dorsal horn is divided into discrete neural precursor domains via the expression of specific transcription factors that control their identities, but whose link to adult neuronal classes remains obscure (Lai et al., 2016). Whereas some spino-thalamic neurons express the transcription factor LIM homeobox transcription factor 1b (Lmx1b) (Szabo et al., 2015), a marker of the dI5 spinal progenitor domain, so do many dorsal horn interneurons. In contrast, the Paired-like homeobox 2a (Phox2a) transcription factor is a more selective marker of developing dI5 neurons, although its transient expression prevents investigation of their adult function (Ding et al., 2004). Interestingly, Phox2a and its close relative Phox2b are required for the development of the autonomic nervous system (Pattyn et al., 1997), with Phox2a being required for the formation of the locus coeruleus (Brunet and Pattyn, 2002; Morin et al., 1997). Since AS neurons have been proposed to subserve autonomic nuclei of the CNS (Craig, 1996), we considered whether developmental Phox2a expression may be their uniting feature.

Here, using genetic fate mapping, we report that transient embryonic expression of Phox2a in spinal neurons defines the identity of several AS projection neuron classes. Using this insight, we reveal a developmental diversity of AS neurons and show that a loss of Phox2a impairs AS neuron innervation of their brain targets, resulting in attenuated supraspinal responses to noxious stimuli. Furthermore, we show that the molecular identity of Phox2a AS neurons is conserved in the developing human spinal cord, suggesting an evolutionarily conserved molecular logic of AS function.

## Results

### Spinal Phox2a^Cre^ neurons reside in lamina I, V and LSN

Mouse *Phox2a* and its proxy, BAC transgene *Phox2a*^*GFP*^ are expressed embryonically and perinatally in the superficial and deep dorsal horn, where many AS neurons reside (Allen Institute for Brain Science, 2008; GENSAT, 2008). In order to label the adult descendants of these neurons, we created the transgenic *Phox2a*^*Cre*^ mouse line by inserting a Cre-polyA minigene into the BAC RP23-333J21 (GENSAT, 2008), at the *Phox2a* ATG codon (Fig. 1A), and assessed Cre expression via the Cre-dependent tdTomato reporter *R26*^*LSL-tdT*^ (Ai14). Adult *Phox2a*^*Cre*^; *R26*^*LSL-tdT*^ mice showed tdTomato (tdT) expression throughout the rostrocaudal length of the spinal cord in dorsal horn neurons, principally in lamina I (Fig. 1B) and lamina V/Lateral Spinal Nucleus (LSN; Fig. 1B, S1A), as well as in spinal accessory nerve (mXI) motor neurons (Fig. S1A). Although rare, large “antenna”-like neurons were also found in laminae III/IV (Fig. S3) (Marshall et al., 1996; Schoenen, 1982), which have been shown to receive a wide range of primary afferent inputs (Fernandes et al., 2018). Phox2a is expressed in embryonic day (e) 11.5 spinal cords, within the dI5 cardinal spinal neuron domain. At this age in *Phox2a*^*Cre*^; *R26*^*LSL-tdT*^ spinal cords, 91% of tdT+ cells co-expressed Phox2a, while at e16.5, as Phox2a expression begins to wane, this proportion decreased to 76% and then to 0% in adults (Fig. 1C, 1D). Conversely, at e11.5, 74% of all Phox2a cells co-expressed tdT, and this proportion decreased to 45% by e16.5. This low co-expression is primarily accounted for by lamina V/LSN Phox2a+ (Phox2a^Deep^) cells, 33% of which expressed tdT, in contrast to lamina I neurons (Phox2a^LamI^) for which this fraction was 82% (Fig. 1C,E). Similar proportions were observed at e18.5 (Fig. S1B), arguing against a delayed onset of Cre expression in Phox2a^Deep^ neurons. Together, these data constitute evidence that *Phox2a*^*Cre*^ can be used to trace the fate of Phox2a-expressing spinal neurons and is a potential genetic tool for AS neuron manipulation.

**Figure 1:**
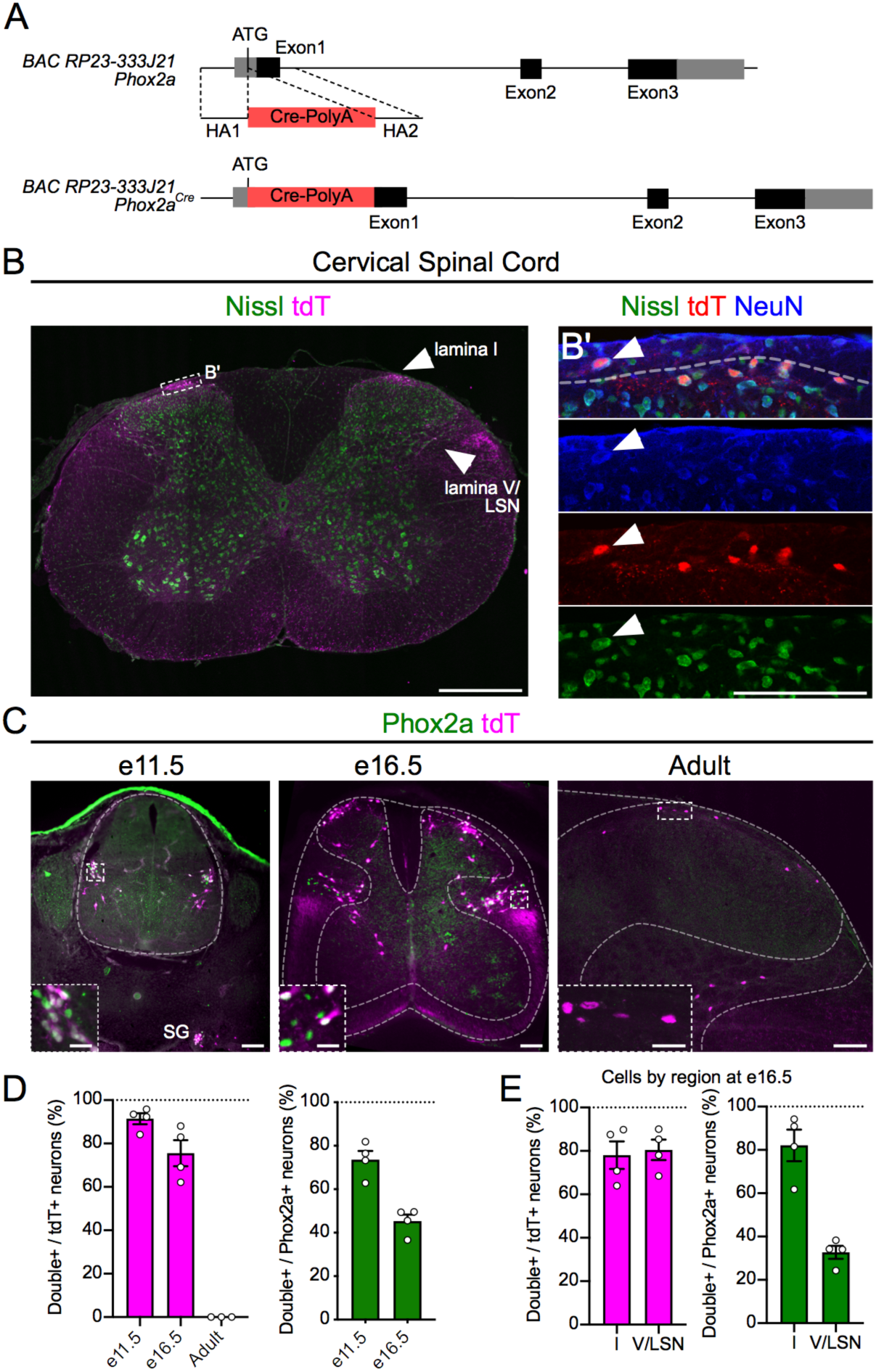
Spinal Phox2a^Cre^ neurons reside in lamina I, V and LSN. (A) BAC recombination strategy: *Cre-PolyA* insertion 3’ to the *Phox2a* ATG codon in the BAC RP23-333J21. (B) tdTomato (tdT)+ neurons in laminae I, V and LSN of the cervical spinal cord of adult *Phox2a*^*Cre*^; *R26*^*LSL-tdT/*+^ mice. (B’) Magnified box in (B) showing lamina I Neurotrace, tdT and NeuN co-labeling. (C) Expression of tdT and Phox2a in e11.5, e16.5 and adult *Phox2a*^*Cre*^; *R26*^*LSL-tdT/*+^ mouse spinal cord. (D) Percent of tdT+ neurons that express Phox2a, as well as percent of Phox2a+ neurons that express tdT at e11.5, e16.5 and adult *Phox2a*^*Cre*^; *R26*^*LSL-tdT/*+^ mice. (E) Percent of tdT+ neurons that express Phox2a, as well as percent of Phox2a+ neurons that express tdT in the superficial and deep dorsal horn of e16.5 *Phox2a*^*Cre*^; *R26*^*LSL-tdT/*+^ mouse spinal cords. Data are represented as mean ± SEM. Numbers: n=4 e11.5, n=4 e16.5 and n=3 adult *Phox2a*^*Cre*^; *R26*^*LSL-tdT/*+^ mice. Scale bars: (B) 500 μm, (B’) 100 μm, (C) 100 μm and insets 25 μm. Abbreviations: SG (sympathetic ganglia).

### Spinal Phox2a^Cre^ neurons innervate AS targets

To reveal the connectivity of spinal *Phox2a*^*Cre*^ neurons, we restricted *Phox2a*^*Cre*^-driven reporter expression to the spinal cord using the Cre-Flp recombinase-dependent reporter *R26*^*FSF-LSL-tdT*^ (Ai65) combined with the caudal neural tube-specific Flp recombinase driver *Cdx2*^*FlpO*^ to generate *Phox2a*^*Cre*^; *Cdx2*^*FlpO*^; *R26*^*FSF-LSL-tdT*^ mice ((Britz et al., 2015) Fig. 2A). To validate this genetic intersection, we compared cellular tdT reporter expression between adult *Phox2a*^*Cre*^; *R26*^*LSL-tdT*^ (Fig. 2B–F, Fig. S2E–H) and *Phox2a*^*Cre*^; *Cdx2*^*FlpO*^; *R26*^*FSF-LSL-tdT*^ mice (Fig. 2B’–F’, Fig. S2E’–H’). In the brain, *Phox2a*^*Cre*^ drove cellular tdT expression in motor and autonomic nuclei (Fig. 2B–E, S2A–H), which was not observed in *Phox2a*^*Cre*^; *Cdx2*^*FlpO*^; *R26*^*FSF-LSL-tdT*^ mice (Fig. 2B’–E’, S2E’–H’). In the caudal spinal cord of *Phox2a*^*Cre*^; *Cdx2*^*FlpO*^; *R26*^*FSF-LSL-tdT*^ mice however, the cellular expression of tdT+ expression was preserved (Fig. 2F, F’), allowing us to map Phox2a-expressing neuron axonal trajectories and brain targets.

**Figure 2:**
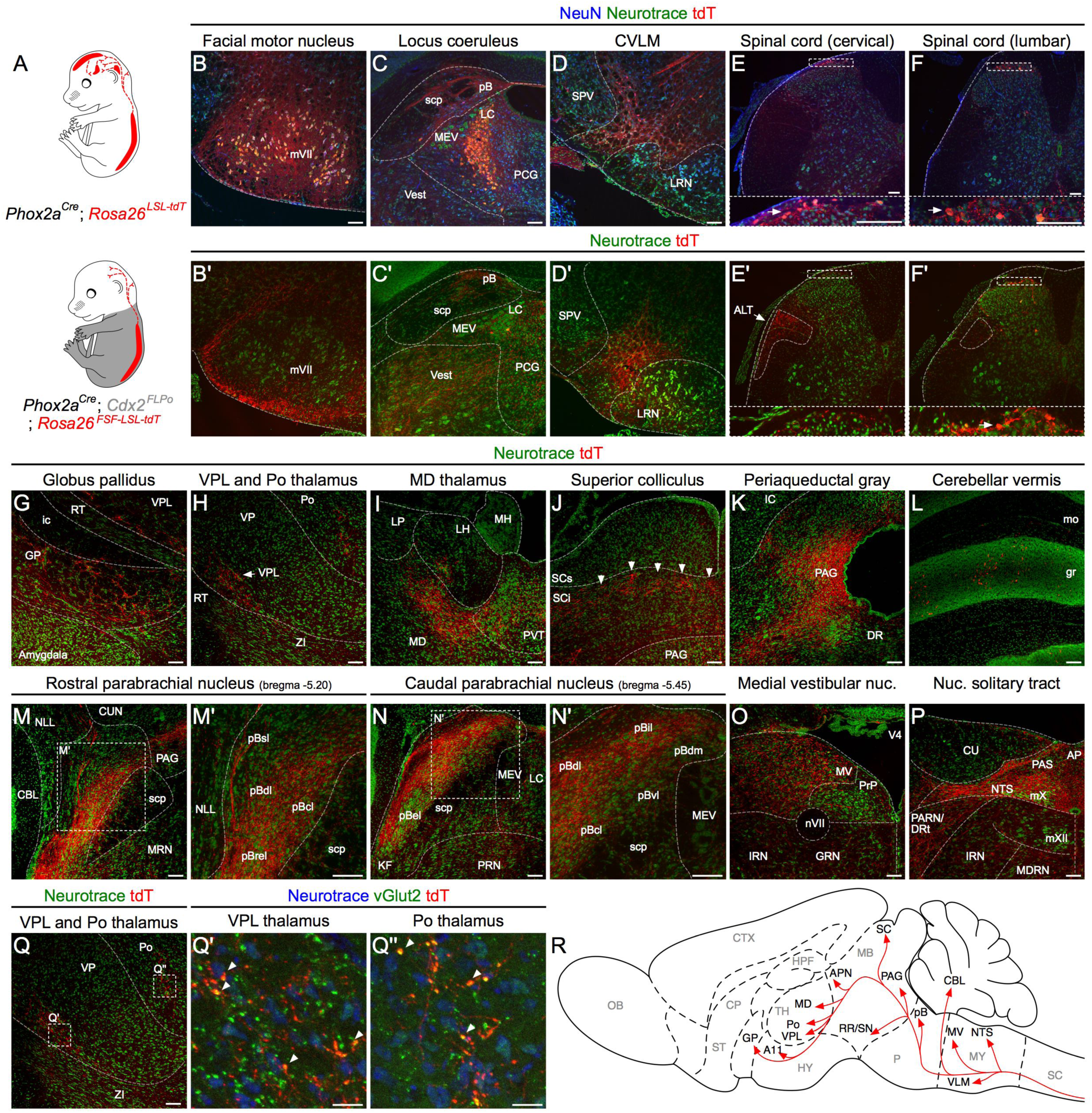
Spinal Phox2a^Cre^ neurons innervate AS targets. (A) Intersectional genetic strategy to visualise spinofugal axons with tdT. *Phox2a*^*Cre*^; *R26*^*LSL-tdT/*+^ mice have tdT cellular expression in the brain and spinal cord while *Phox2a*^*Cre*^; *Cdx2*^*FlpO*^; *R26*^*FSF-LSL-tdT/*+^ mice have cellular tdT expression only in spinal Phox2a neurons. (B–F) *Phox2a*^*Cre*^; *R26*^*LSL-tdT/*+^ mouse tdT expression, and NeuN and Neurotrace staining in the facial motor nucleus (B), locus coeruleus (C) and the caudal ventrolateral medulla (CVLM) (D), as well as the spinal dorsal horn (E, F). (B’–F’) Same regions as in (B–F) have no cellular tdT expression in *Phox2a*^*Cre*^; *Cdx2*^*FlpO*^; *R26*^*FSF-LSL- tdT/*+^ mice except below the cervical spinal cord. (E’) Arrow: presumptive anterolateral (ALT) tract axons in white matter, not detectable in (E) presumably due to weaker axonal tdT expression from *R26*^*LSL-tdT*^. Insets in (E, F, E’ and F’) correspond to stippled boxes and show tdT+ cell bodies (arrows). (G–P) Prominent targets of tdT+ spinofugal axons. Higher magnification insets in M’ and N’. (Q, Q’, Q’’) Neurotrace, tdT and vGluT2 staining in the thalamus. Arrowheads in J indicate axon termini in putative orientation barrels of the superior colliculus. Arrowheads in higher magnification insets in Q’, Q’’ point to putative synaptic termini where tdT co-localises with vGluT2 signal. (R) Diagram summarising the termination sites of tdT+ spinofugal axons. Numbers: n=3 *Phox2a*^*Cre*^; *R26*^*LSL-tdT/*+^ adult mice, n=3 *Phox2a*^*Cre*^; *Cdx2*^*FlpO*^; *R26*^*FSF-LSL-tdT/*+^ adult mice. Scale bars: 100 μm, except (Q’, Q’’) 25 μm. Abbreviations: ALT (anterolateral tract), AP (area postrema), CBL (cerebellum), CU (cuneate nucleus), CUN (cuneiform nucleus), DR (dorsal raphe), DRt (dorsal reticular nucleus), GP (globus pallidus), gr (granular layer of the cerebellum), GRN (gigantocellular reticular nucleus), ic (internal capsule), IRN (intermediate reticular nucleus), KF (Kölliker-Fuse nucleus), LC (locus coeruleus), LH (lateral habenula), LP (lateral posterior thalamus), LRN (lateral reticular nucleus), MD (mediodorsal thalamus), MDRN (medullary reticular nucleus), MEV (midbrain trigeminal nucleus), MH (medial habenula), mo (molecular layer of the cerebellum), MV (medial vestibular nucleus), mVII (facial motor nucleus), mX (vagal motor nucleus), mXII (hypoglossal motor nucleus), NLL (nucleus of the lateral lemniscus), NTS (nucleus of the solitary tract), nVII (facial motor nerve), PAG (periaqueductal gray), PARN (parvocellular reticular nucleus), PAS (parasolitary nucleus), pB (parabrachial nucleus), pBcl (central-lateral parabrachial nucleus), pBdl (dorsal-lateral parabrachial nucleus), pBdm (dorsal-medial parabrachial nucleus), pBel (external-lateral parabrachial nucleus), pBil (internal-lateral parabrachial nucleus), pBrel (rostral external-lateral parabrachial nucleus), pBsl (superior-lateral parabrachial nucleus), pBvl (ventral-lateral parabrachial nucleus), PCG (pontine central gray), Po (posterior thalamus), PRN (pontine reticular nucleus), PRP (nucleus prepositus), PVT (paraventricular thalamus), RT (reticular thalamic nucleus), SCi (superior colliculus, intermediate laminae), scp (superior cerebellar peduncle), SCs (superior colliculus, superficial laminae), SPV (spinal trigeminal nucleus), Vest (vestibular nuclei), VP (ventral posterior thalamus), VPL (ventral posterolateral thalamus), ZI (zona incerta).

In *Phox2a*^*Cre*^; *Cdx2*^*FlpO*^; *R26*^*FSF-LSL-tdT*^ mice, tdT+ axons were observed in the lateral funiculus in a similar distribution to previous reports of lamina I spinofugal axon locations (Apkarian et al., 1985; McMahon and Wall, 1983) (Fig. 2E’). We observed tdT+ axons in known AS targets such as the globus pallidus (Fig. 2G), VPL and posterior (Po) thalamus (Fig. 2H, S2I), mediodorsal thalamus (MD, Fig. 2I), the posterior triangular thalamus (PoT) and anterior pretectal nucleus (Fig. S2K), the deep layers of the superior colliculus, possibly within the orientation barrels (Masullo et al., 2019) (Fig. 2J), periaqueductal gray (PAG) (Fig. 2K), the pB, (Fig. 2M, N), the nucleus of the solitary tract (Fig. 2P), the locus coeruleus (Fig. 2C’) and the caudal ventrolateral medulla (CVLM) (Fig. 2D’). These termini contained the presynaptic marker vGluT2 suggesting that they contained glutamatergic synapses (Fig. 2Q, Q’). Within the pB, the dorsal-lateral (pBdl), central-lateral (pBcl), internal-lateral (pBil) subnuclei and regions surrounding the external-lateral (pBel) contained many tdT+ axons, while the superior-lateral (pBsl) and medial (pBm) subnuclei contained fewer axons (Fig. 2M, N, Fig. S2M, N). Consistent with previous reports, the pBel received very limited spinal innervation (Fig. 2N, S2E, S2N; (Bernard et al., 1995)). Additionally, spinal Phox2a^Cre^ axons were also seen in brain regions not previously thought to receive direct AS innervation, such as the granular layers of the cerebellum (Fig. 2L), the vestibular nuclei, (Fig. 2O), the posterior hypothalamus near the A11 dopaminergic cell group (Fig. S2J), and a region of the retrorubral area / dorsomedial substantia nigra (Fig. S2L). Thus, spinal Phox2a neurons innervate brain regions predominantly involved in autonomic regulation and homeostasis such as the pBdl, the nucleus of the solitary tract (NTS) and CVLM, as well as nociceptive areas (VPL, PAG, pBil). The identities of these targets suggest that spinal Phox2a neurons may orchestrate a wide range of pain-related and autonomic responses.

### Spinal Phox2a neurons are predominantly AS neurons

We next considered whether Phox2a expression could be the feature uniting the morphologically, anatomically and physiologically diverse classes of AS neurons. Thus, we determined the fraction of AS neurons retrograde labelled from their principal targets that also expressed tdT (referred to as *Phox2a*^*Cre*^ neurons). We focused our analysis on the VPL thalamus and the pB, whose tracer injections results in efficient labelling of their afferent AS neurons. Adult *Phox2a*^*Cre*^; *R26*^*LSL-tdT*^ mice of both sexes were injected unilaterally with fluorogold (FG) in the VPL thalamus (Fig. 3A), and with CTb-488 in the pB (Fig. 3B). After 7 days, we examined the proportion of spinal neurons labelled with either or both tracers (Tracer+) that were also tdT+, sampled at all spinal cord levels (1023 FG+, 6620 CTb-488+ and 3345 tdT+ cells from 7 mice), although we focussed our analysis on the cervical spinal cord, as spino-thalamic neurons are relatively sparse in the mouse caudal spinal cord (Davidson et al., 2010). Overall, *Phox2a*^*Cre*^ labelled similar ratios of AS neurons traced from the VPL and the pB (26.9±5.0 % and 19.7±4.3 %, respectively, n=7), and, in a separate experiment, the MD thalamus (22.8±2.5 %; Fig. S3, n=3). These results corroborated the genetic anterograde tracing experiments and formally demonstrate that developmental Phox2a expression is a feature of many AS neurons.

**Figure 3:**
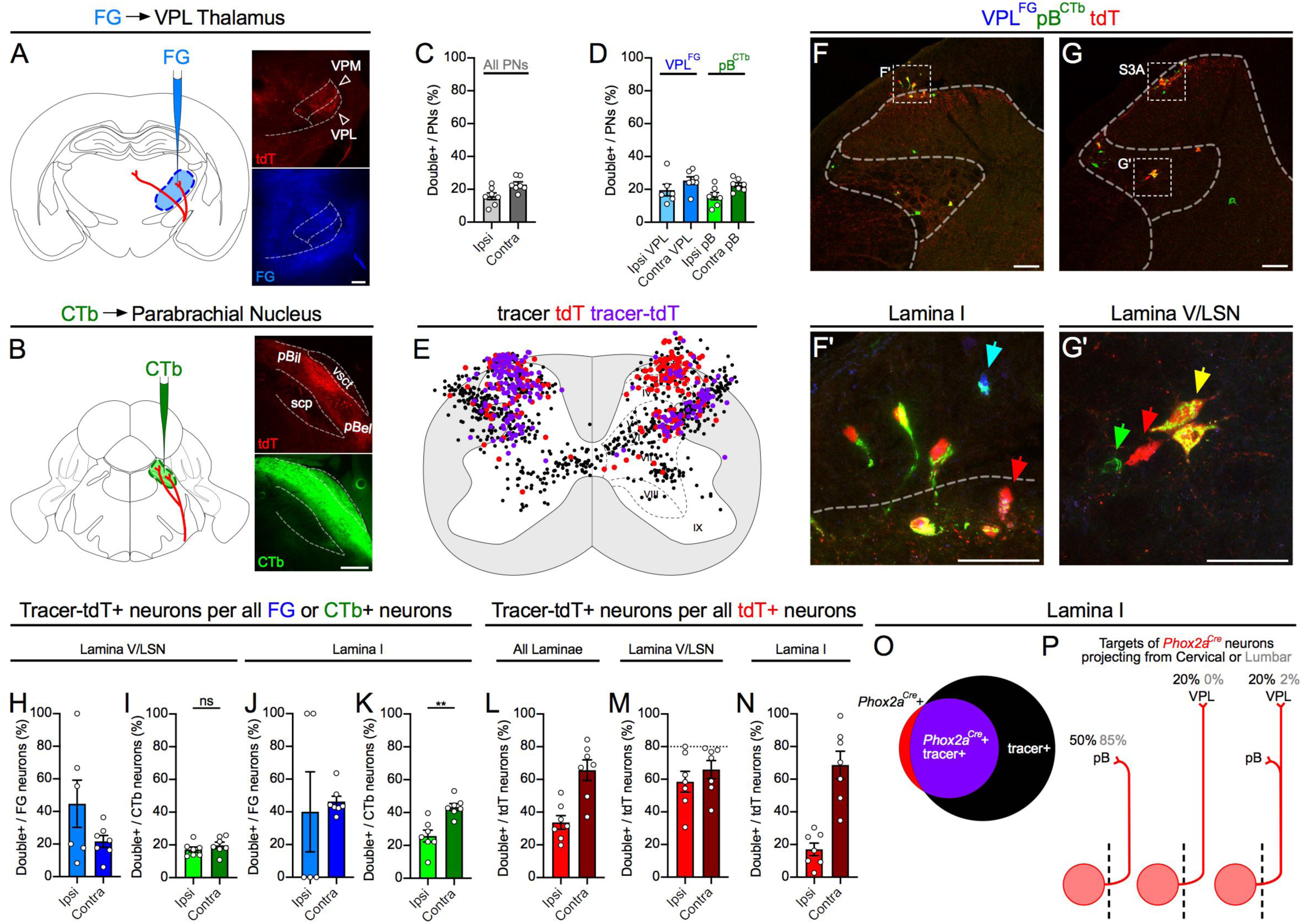
Spinal Phox2a^Cre^ neurons are predominantly AS neurons. Adult *Phox2a*^*Cre*^; *R26*^*LSL-tdT/*+^ mice injected with FG in the VPL thalamus and CTb-488 in the parabrachial nucleus. (A, B) Representative image of FG (A) and CTb-488 (B) injection sites. (C, D) Percent of cervical spinal cord dorsal horn projection neurons expressing tdT, classified as those labelled with either tracer (All PNs) in (C) or those labelled selectively with FG or CTb in (D). In this quantification scheme, some neurons positive for one tracer may also be positive for the other. (E) Diagram of location of tdT+ only (red), retrograde label only (FG or CTb, black) or tdT+ and tracer-labelled (purple) neurons, in 5 non-sequential 25 µm sections of the cervical spinal cord of one representative animal. (F, G’) Representative images of the cervical spinal cord demonstrating tdT+ neuron labelling by FG or CTb tracer injections in, respectively, the VPL or the pB. (F’) High magnification of boxed area in lamina I in (F). (G’) High magnification of boxed area in lamina V/LSN in (G). Also see Fig. S3A for lamina I box in (G). (F’, G’) Red arrow: tdT-only cell; cyan arrow: FG and CTb double-labelled cell; green arrow: CTb-only cell; yellow arrow: tdT+ cell labelled with CTb. (H, I) Percent of FG (H) or CTb (I)-labelled lamina V/LSN neurons also expressing tdT, in the cervical spinal cord ipsilateral or contralateral to tracer injection. (J, K) Percent of FG (H) or CTb (I)-labelled lamina I neurons also expressing tdT, in the cervical spinal cord ipsilateral or contralateral to tracer injection. (L–N) Percent of tdT-labelled neurons labelled with either or both tracers in all laminae (L), in lamina V/LSN (M), or in lamina I (N) in the cervical spinal cord ipsilateral or contralateral to tracer injection. (O) Diagram depicting overlap between tdT and retrograde tracer in lamina I of a representative mouse. (P) Diagrams illustrating the estimated percentages of cervical and lumbar lamina I Phox2a^Cre^ neurons projecting to mouse VPL and/or pB. Stippled line represents spinal midline. Note the high degree of VPL/pB collateralised innervation by cervical Phox2a^Cre^ neurons. Data are represented as mean ± SEM. Numbers: n=7 *Phox2a*^*Cre*^; *R26*^*LSL-tdT/*+^ adult mice (4 male, 3 female). Statistics: (I, K) Mann-Whitney test, ns: non-significant, **: p<0.01. Scale bars: (A, B) 250 μm, (F, G) 100 μm, (F’, G’) 50 μm.

Since Phox2a expression did not correlate strongly with AS neuron target identity, we next examined whether it may be linked to AS neuron laterality or laminar position. Overall, *Phox2a*^*Cre*^ labelled 16% of ipsilateral and 23% of contralateral Tracer+ neurons (Fig. 3C), with similar fractions labeled from the VPL and pB (Fig. 3D). A two-dimensional distribution of Phox2a^Cre^ labelled AS neurons demonstrated a concentration in the contralateral lamina I (Fig. 3E, F) where high rates of co-localization occurred, in contrast to lamina V/LSN (Fig. 3E, G) where neurons were frequently seen labeled only with retrograde tracer. Indeed, tracer labelled neurons expressing tdTomato were far less frequent in lamina V/LSN (Fig. 3H, I) than lamina I (Fig. 3J, K), likely due to Phox2a^Cre^ underreporting Phox2a expression in deep laminae. Together, these data demonstrate that *Phox2a*^*Cre*^ expression defines approximately 20% of all spino-thalamic and spino-parabrachial AS neurons, and approximately half of AS neurons in the superficial dorsal horn. Additionally a strong tdT expression bias in contralaterally projecting lamina I AS neurons, many of which are organised somatotopically, suggests that many *Phox2a*^*Cre*^ neurons are involved in the localisation of noxious stimuli (Fig. 3K).

We next quantified the fraction of tdT+ neurons that contribute to the AS. We speculated that if all *Phox2a*^*Cre*^ neurons were AS neurons, then a highly efficient tracer injection would result in tracer accumulation in all tdT+ neurons. In our most comprehensive injections of tracer into the pB and VPL, we reached a labelling ceiling of ∼80% of lamina V/LSN tdT+ neurons bilaterally and as much as 100% of lamina I tdT+ neurons, strongly suggesting that all spinal *Phox2a*^*Cre*^ neurons give rise to the AS (Fig. 3L–N). Given the heterogeneity of LSN neuron targets (Leah, 1988), it is likely that the tdT+ neurons in the LSN unlabelled by the tracers project to AS targets other than the VPL or the pB. Among lamina I and lamina V/LSN neuron types, smaller fractions were labelled by tracer injection into to pB or VPL suggesting *Phox2a*^*Cre*^ neurons represent a variety of projection types (Fig. 3L vs Fig. S3C, D). The efficiency of tracer labelling of the rare antenna and lamina X tdT+ neurons was too low to quantify with confidence, although they predominantly to project contralaterally (Fig. S3G–K). We also examined spinal projections to the MD thalamus via retrograde tracer injection, which labelled much fewer neurons than pB/VPL injections, but also included tdT+ neurons (Fig. S3L– R). In the hindbrain, pB, VPL and MD retrograde tracer injections also labelled tdT+ neurons in the CVLM, parvocellular reticular nucleus and spinal trigeminal lamina I / paratrigeminal region, suggesting these Phox2a^Cre^ neurons share axonal targets and perhaps functions with spinal Phox2a^Cre^ neurons. Together, our data demonstrate that spinal tdT (*Phox2a*^*Cre*^) expression is a nearly exclusive label of AS neurons.

### Heterogeneity of spinal Phox2a neuron migration, sensory afferent interaction and birth time

To gain insights into the functional diversification of Phox2a AS neurons implied by their dorsal horn laminar location and connectivity, we turned to the cellular and molecular events underlying their development. We first asked whether the laminar distribution of Phox2a neurons is a consequence of radial migration, typical of laminated CNS structures. We examined this possibility by following Phox2a and tdT expression in *Phox2a*^*Cre*^; *R26*^*LSL-tdT*^ spinal cords throughout embryonic development. The first Phox2a neurons appear at e10.5 in the cervical region and begin expressing tdT one day later (Fig. 4A). At e12.5, three Phox2a populations are evident: Phox2a+ tdT+ (Phox2a^LamI^) neurons ventrolateral to the nascent dorsal horn, and two medial populations consisting of Phox2a+ tdT+ and those expressing tdT alone. At e13.5, Phox2a^LamI^ neurons disperse on the surface of the nascent superficial dorsal horn in a tangential orientation, while deeper Phox2a neurons (Phox2a^Deep^) acquire distinct positions that correlate with tdT expression: Phox2a^Deep^ tdT+ neurons remained ventrolateral to the dorsal horn, while Phox2a^Deep^ tdT-neurons accumulated above the central canal. At e14.5, Phox2a^Deep^ tdT-neurons translocate laterally and eventually become intermingled with Phox2a^Deep^ tdT+ neurons at e15.5, achieving their final configuration (Fig. S4A). Collectively we are able to identify three distinct migratory paths of Phox2a neurons based on their Phox2a^Cre^ expression.

**Figure 4:**
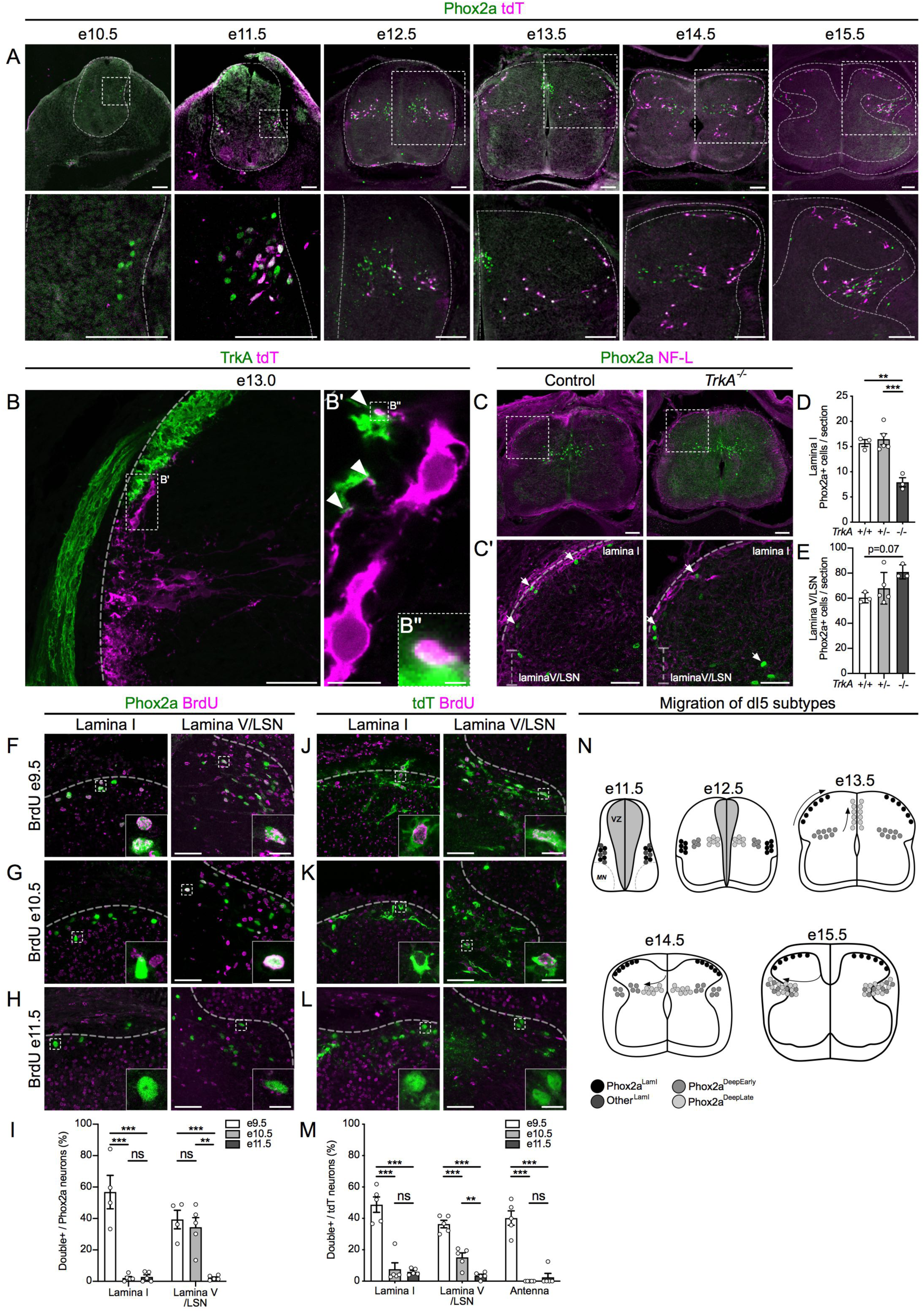
Heterogeneity of spinal Phox2a neuron migration, sensory afferent interaction and birth time. (A) Migration of Phox2a+ (green), tdT+ (magenta) and Phox2a+ tdT+ (white) neurons in embryonic spinal cords of *Phox2a*^*Cre*^; *R26*^*LSL-tdT/*+^ mice aged between e10.5 and e15.5. Boxed regions in upper panels are magnified below. Spinal cord and spinal white matter are bounded by stippled lines (B, B’) Location of tdT+ neurons in the dorsal horn of *Phox2a*^*Cre*^; *R26*^*LSL-tdT/*+^ spinal cords at e13.0, highlighting contacts (B’, B’’) between lamina I neurons (magenta) and TrkA+ sensory afferents arriving in the dorsal horn (green). Micrograph in B is a flattened multi-layer z-stack, while images in B’, B’’ are single confocal micrographs. (C–E) Spinal Phox2a (green) neuron and Neurofilament light chain (NF-L, magenta) localisation in e14.5 *TrkA*^+*/*+^, *TrkA*^+*/-*^ and *TrkA*^*-/-*^ mouse embryos. Boxed regions in (C) magnified in (C’) with arrows pointing to Phox2a cells in laminae I, V and the LSN. Counts of Phox2a neurons in lamina I (D) and lamina V/LSN (E). (F–I) Birthdating of spinal Phox2a neurons in e16.5 *Phox2a*^*Cre*^; *R26*^*LSL-tdT/*+^ mouse embryos, exposed to BrdU at e9.5 (F), e10.5 (G) or e11.5 (H). Phox2a+/BrdU+ neurons as a percent of all Phox2a+ neurons in either lamina I or lamina V/LSN and compared between groups (I). (J–M) Birthdating of spinal tdT+ neurons in e16.5 *Phox2a*^*Cre*^; *R26*^*LSL- tdT/*+^ mouse embryos, exposed to BrdU at e9.5 (J), e10.5 (K) or e11.5 (L). tdT+/BrdU+ neurons numbers as a percent of all tdT+ neurons in either lamina I, lamina V/LSN or laminae II/III (“Antenna”-like cells) and compared between groups (M). tdT was detected with an anti-red fluorescent protein (RFP) antiserum. (N) Diagram of migration and birth patterns of spinal Phox2a neuron subpopulations. Data are represented as mean ± SEM. Numbers: *Phox2a*^*Cre*^; *R26*^*LSL-tdT/*+^ embryos: (A) n=3 e10.5, n=3 e11.5, n=3 e12.5, n=3 e13.5, n=3 e14.5, n=3 e15.5, (B, B’, B’’) n=3 e13.0, (F–M) n=4-5 e16.5 per condition. (C–E) n=3 *TrkA*^+*/*+^, n=5 *TrkA*^+*/-*^, n=3 *TrkA*^*-/-*^. Statistics: (D, E) One-way ANOVA, with Tukey’s Multiple Comparisons test, (I,M) individual one-way ANOVAs for each cell type (lamina I, lamina V/LSN and Antenna) with Tukey’s multiple comparisons test; **: p<0.01, ***: p<0.001. Scale bars: (A, C) 100 μm, (B, C’, F–H, J–L) 50 μm, (B’) and insets in (F–H, J–L) 10 μm, and (B’’) 1 μm.

As the tangential dispersal of Phox2a^LamI^ neurons within the dorsal horn occurs at the time of primary afferent innervation of this domain, we asked how these two events are related. In e12.5 *Phox2a*^*Cre*^; *R26*^*LSL-tdT*^ spinal cords, prior to their entry into lamina I, *Phox2a*^*Cre*^ neurons project tdT+ processes towards the dorsal root entry zone, the arrival site of sensory afferent axons that eventually synapse with AS neurons (S4C,D). At e13.0, at the onset of Phox2a^LamI^ neuron migration into lamina I (Fig. 4B), such tdT+ processes were found in close apposition to TrkA+ nociceptive primary afferent axons (Fig. 4B’, S4B’, S4B’’), which enter the superficial dorsal horn, and eventually surround Phox2a^LamI^ neurons (Fig. S4E, S4F). To determine whether TrkA+ axons contribute to Phox2a^LamI^ neuron positioning, we examined the location of Phox2a neurons in *TrkA*-null (*TrkA*^-/-^) mouse embryos, in which most TrkA+ primary afferents are absent (Smeyne et al., 1994). Compared to controls, the number of Phox2a^LamI^ neurons in *TrkA*^-/-^ embryos was approximately halved in the superficial dorsal horn (Fig. 4C–E), suggesting that the interplay between nociceptive afferents and their eventual synaptic targets is functionally important. In contrast, although we found no significant effects of TrkA+ axon loss on Phox2a^Deep^ neuron count, it tended to increase, consistent with Phox2a^LamI^ neurons’ failure to migrate (Fig. 4C–E; data not shown).

To determine whether spinal Phox2a neuron diversity and migration patterns correlate with the time of their birth, we injected pregnant *Phox2a*^*Cre*^; *R26*^*LSL-tdT*^ mice with Bromodeoxyuridine (BrdU) at e9.5, 10.5 and e11.5, and examined strong BrdU co-staining with Phox2a or tdT in e16.5 *Phox2a*^*Cre*^; *R26*^*LSL-tdT*^ embryos (Fig. 4F–M). Nearly all Phox2a^LamI^ neurons were born at e9.5, while Phox2a^Deep^ neurons were born between e9.5 and e10.5. Very few neurons of either type were born at e11.5 suggesting that by that age, all Phox2a AS neurons have been born. Furthermore, examination of e11.5 embryos labelled with BrdU at e9.5 or e10.5 revealed that e11.5 Phox2a neurons are predominantly born at e9.5; this, together with our migration analysis and e16.5 birthdating, supports the notion that the earliest Phox2a neurons to appear become Phox2a^LamI^ neurons (Fig. S4G–M). Taking advantage of the differential expression of Phox2a and tdT in Phox2a^Deep^ cells, our data argue that Phox2a^LamI^ and antenna neurons are born first, followed by Phox2a^Deep^ tdT+ (Phox2a^DeepEarly^), while Phox2a^Deep^ tdT-neurons are born last (Phox2a^DeepLate^) but not beyond e11.5 (Fig. S4N–Q). Thus, spinal Phox2a neuron birthdate correlates with their migration trajectory and defines a distinct set of AS neuron types (Fig. 4N). More generally, our data argue against previous conclusions that AS neurons are born concurrent with dorsal horn neurons (Nornes and Carry, 1978), and show that AS neurons constitute one of the earliest-born spinal neuron populations (Fig. S4R–T).

### The molecular identity and specification of spinal Phox2a neurons

To uncover the molecular pathways controlling Phox2a AS neuron specification, we studied their expression of neuronal identity determinant genes, identified transcription factor programs that specify them, and sought molecular markers that subdivide them. Spinal Phox2a expression begins at e9.5 in accessory motor neurons (Fig. S5A–D), likely the precursors of e10.5 cervical spinal cord Phox2a+ neurons that co-express Phox2b and Isl1. Non-motor neuron Phox2a expression is first visible at e10.5 in post-mitotic neurons expressing the cardinal dI5 transcription factor Lmx1b, adjacent to dorsal interneuron (dI) progenitors expressing the Ascl1 or Pax7 transcription factors, (Fig. 5A–D, M). Phox2a/Lmx1b neurons also express the dI5 transcription factors Lbx1, Tlx3, and Brn3b/Pou4F2 but not the dI1, 3, 4/6 transcription factors Lhx2, Isl1, or Pax2, respectively (Fig. 5E–J, M). Phox2a/Lmx1b neurons also express the commissural neuron guidance receptors Robo3 and DCC (Fig. 5K–M). These findings were largely similar at e11.5, when first tdT+ cells appear in Phox2a+ dI5 neurons of *Phox2a*^*Cre*^; *R26*^*LSL-tdT*^ spinal cords (Fig. S5E–O) demonstrating that non-motor neuron spinal Phox2a cells are predominantly commissural dI5 neurons.

**Figure 5:**
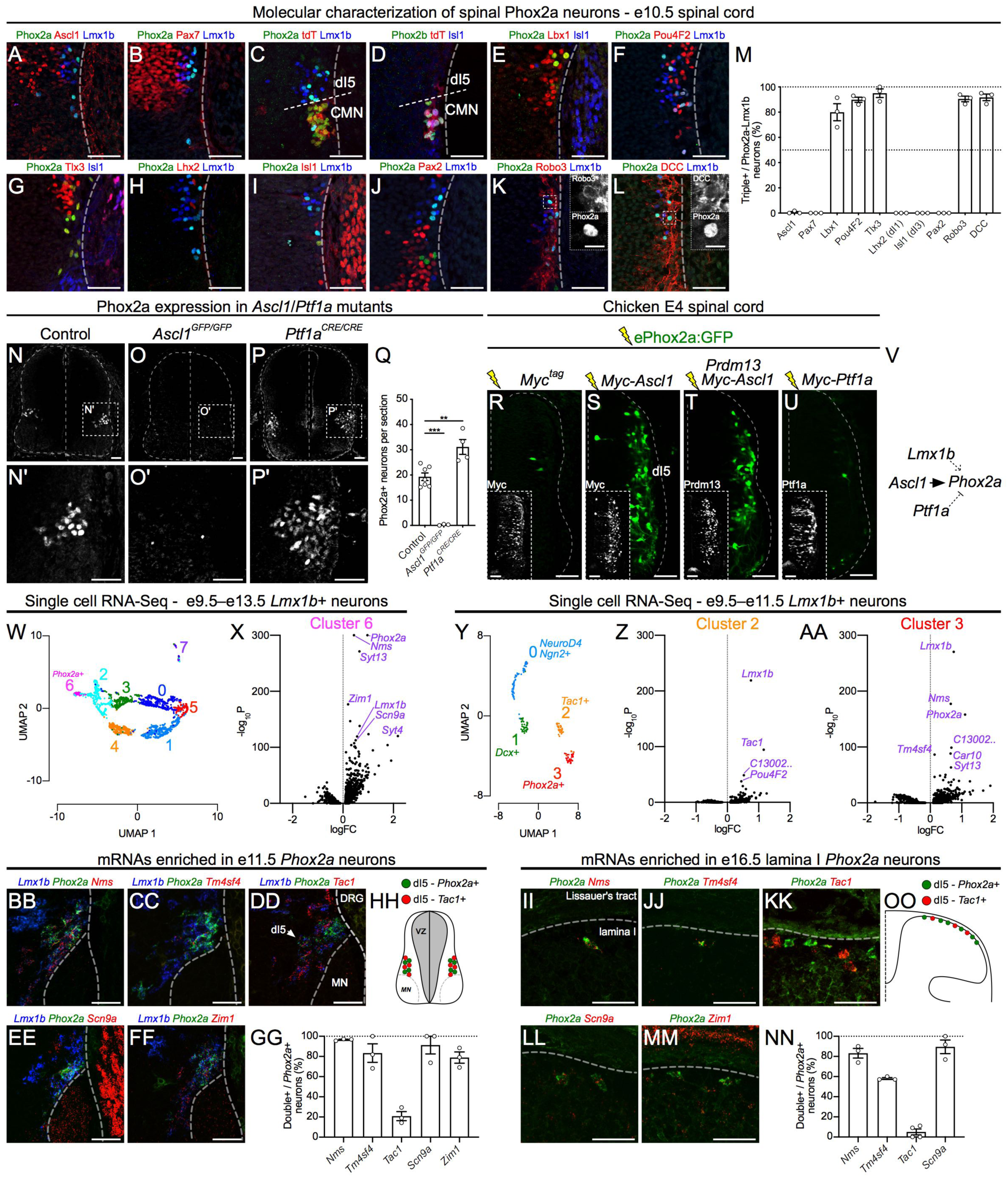
The molecular identity and specification of spinal Phox2a neurons. (A–M) Molecular characterisation of spinal Phox2a neurons in the e10.5 *Phox2a*^*Cre*^; *R26*^*LSL-tdT/*+^ spinal cord. (A, B) Lack of co-expression of Phox2a and progenitor markers Ascl1 (A) and Pax7 (B). (C,D) Spinal dI5 tdT+ neurons co-express Phox2a, Lmx1b but not the spinal accessory motor neuron (SMN) markers Phox2b or Isl1. (E–J) Spinal Phox2a neurons express dorsal interneuron markers Lbx1 (E), Pou4F2 (F), Tlx3 (G), but not Lhx2 (H), Isl1 (I), or Pax2 (J). (K, L) Spinal Phox2a neurons express commissural neuron markers Robo3 (K) and DCC (L). (M) Quantification of marker expression as a percentage of Phox2a and Lmx1b co-expressing cells. (N–P) Phox2a expression in control (N, N’), *Ascl1* null (O, O’), and *Ptf1a* null (P, P’) E11.5 spinal cords. (N’, O’ and P’) High magnification of boxed regions, in respectively, (N, O and P). (Q) Average numbers of Phox2a+ cells in spinal cord sections in control, *Ascl1* and *Ptf1a* mutant e11.5 spinal cords. (R–U) Representative images from transverse sections of embryonic day 4 chick neural tubes co-electroporated with the *ePhox2a-GFP* reporter and expression plasmids: control (Myc-tag only) (R), ^*Myc*^*Ascl1* (S), ^*Myc*^*Ascl1* and *Prdm13* (T), or ^*Myc*^*Ptf1a* (U). Insets show Myc-tag expression. (V) Diagram of *Phox2a* expression regulation showing direct activation by Ascl1, potential activation by Lmx1b (Fig. S5), and inhibition by Ptf1a. (W–A1) UMAP Analysis using single-cell RNA sequencing data (Delile et al., 2019). (W–X) *Lmx1b*+ neurons from e9.5-e13.5 compared between each other; cluster 6 is enriched in *Phox2a*+ neurons (W). Other potential cluster 6-enriched mRNAs are revealed in a volcano plot (X). (Y–AA) *Lmx1b*+ neurons from e9.5-e11.5 compared between each other; clusters 2 and 3 are enriched in *Tac1* and *Phox2* mRNAs, respectively (Y), and Cluster 2 and 3-enriched mRNAs are revealed in respective volcano plots (Z, AA). (BB–OO) *In-situ* hybridisation of select mRNAs enriched in lamina I neurons, from UMAP analyses. (BB–FF) *Nms, Tm4sf4, Scn9, Tac1* and *Zim1* mRNAs predicted as enriched in dI5 cells (X, AA) are co-expressed with *Phox2a*, with the exception of *Tac1* which is present in dI5 *Lmx1b*+ neurons that do not express *Phox2a* (DD). (GG) Percent of e11.5 *Phox2a*+ cells co-expressing predicted dI5-enriched mRNAs. (HH) A diagram of dI5 neurons highlighting the non-overlapping populations of dI5-Phox2a+ and dI5-Tac1+ neurons. (II–MM) dI5-enriched mRNAs identified in (X, AA) are co-expressed in *Phox2a*+ lamina I neurons at e16.5, except for *Tac1* (KK). (NN) Quantification of novel dI5-enriched mRNAs co-expression with *Phox2a* in e16.5 lamina I neurons. (OO) Summary diagram demonstrating subdivision of lamina I neurons by Phox2a and Tac1 expression. Data are represented as mean ± SEM. Numbers: *Phox2a*^*Cre*^; *R26*^*LSL-tdT/*+^ embryos (A–M) n=3 e10.5, (BB–GG) n=3 e11.5, (II– NN) n=3-4 e16.5. (N–Q) n=6 control, n=3 Ascl1^GFP/GFP^, n=4 Ptf1a^Cre/Cre^ embryos, (R–U) n=6 E4 chicken embryos for each condition. Statistics: (Q) Student’s t-test, **: p<0.01, ***: p<0.001. (W–AA) Data derived from Delile et al., (2019); data processing and statistics described in STAR Methods. Scale bars: All 50 μm, except (K, L) insets 10 μm. Abbreviations: DRG (dorsal root ganglion), MN (motor neurons).

Since spinal Phox2a neurons are develop from dI5 embryonic neurons, and dI5 neuron identity is specified by the bHLH (basic helix-loop-helix) transcription factor Ascl1 while Ptf1a suppresses dI5 identity and induces the neighbouring dI4 identity (Glasgow et al., 2005; Helms et al., 2005), we assessed whether Phox2a expression was altered in *Ascl1* null (*Ascl1*^*GFP/GFP*^) and *Ptf1a* null (*Ptf1a*^*CRE/CRE*^) e11.5 and e14.5 spinal cords. Compared to littermate controls, virtually no Phox2a neurons were found in e11.5 *Ascl1*^*GFP/GFP*^ spinal cords while additional Phox2a neurons were found in e11.5 and e14.5 *Ptf1a*^*CRE/CRE*^ embryos (Fig. 5N–Q, S5P, S5Q). To determine whether Ascl1 and Ptf1a transcription factors control Phox2a expression directly or indirectly, we analysed ChIP-seq data (Borromeo et al., 2014) for Ascl1 and Ptf1a binding to the *Phox2a* locus. A genomic region (*ePhox2a*) located >30 kb downstream of the *Phox2a* transcription start site was bound by Ascl1 and Ptf1a, although no binding was detected for the Ptf1a co-factor Rbpj or Prdm13, both of which act to repress dI5 and promote dI4 identity (Fig. S5R; (Chang et al., 2013; Hori et al., 2008)). To test the ability of Ascl1, Ptf1a and Prdm13 to regulate Phox2a through *ePhox2a*, we co-electroporated plasmids encoding these proteins together with a plasmid containing an *ePhox2a* activity reporter *(ePhox2a:GFP*; Fig. S5S) into chick spinal neuron progenitors and monitored GFP expression. *ePhox2a:GFP* alone directed GFP expression in a small number of neurons located within the dI5 domain (Fig. 5R, S5T), supporting its function as a dI5-specific enhancer. Ectopic Ascl1, but not ectopic Ptf1a, dramatically increased the number of GFP+ cells (Fig. 5S, T). Furthermore, consistent with the absence of Prdm13 binding to *ePhox2a*, the increase in GFP numbers by Ascl1 expression was not suppressed by Prdm13 (Fig. 5U). Furthermore, Phox2a expression was entirely abolished in Lmx1b^-/-^ e11.5 mouse neural tubes (Fig. S5U). Together, these data suggest that Ascl1 and Lmx1b are required for Phox2a expression and Ascl1 acts directly through a distal 3’ enhancer. In contrast, Ptf1a represses Phox2a transcription, likely through indirect mechanisms (Fig. 5V).

Given that Phox2a labels a set of AS neurons, we sought to identify other genes expressed preferentially within AS neurons using available single cell RNA-Seq data from e9.5-e13.5 mouse spinal cords (Delile et al., 2019). Since Phox2a neurons are a subset of Lmx1b-expressing dI5 neurons, we performed UMAP dimensionality reduction analyses on two cohorts of Lmx1b+ neurons: 1) those found at all time points (e9.5– e13.5) to maximize statistical power for finding differentially expressed AS genes (2614 neurons, Fig. 5W) and 2) an earlier subset of Lmx1b neurons from e9.5–e11.5 spinal cords (186 neurons, Fig. 5Y) to attempt to separate early dI5 neurons (pre-lamina I neurons) into subsets. From both data sets, we were able to isolate a cluster of Lmx1b neurons enriched for Phox2a+ neurons (Fig. 5X, 5AA, S5V, S5W), as well as an early Lmx1b+ cluster enriched for Tac1, a marker for a previously identified neuronal population containing interneurons and AS neurons (Fig. 5Z, S5W). Top enriched transcripts for each cluster are listed in Table S1. Selected candidate transcripts enriched in clusters containing Phox2a+ neurons versus all other neurons were validated using immunohistochemistry and *in situ* mRNA detection, in e11.5 and e16.5 spinal cords. At e11.5 Phox2a neurons were enriched for the expression of *Nms, Tm4sf4, Scn9a*, and *Zim1* mRNAs (Fig. 5BB–HH), which remained expressed in e16.5 Phox2a^LamI^ neurons (Fig. 5II–OO), providing further support that the early Phox2a cells populate the superficial dorsal horn. Other dI5-enriched transcripts and proteins, *Syt4, Pdzrn3*, Shox2 and Pou6F2, were also highly co-expressed with Phox2a, but were less specific to Phox2a neurons (Fig. S5X–EE). Thus, in addition to identifying molecular markers of Phox2a neuron subpopulations, our analyses point to non-overlapping expression of *Phox2a* and *Tac1* in early Lmx1b neurons (likely lamina I-destined) as a potential molecular division of superficial dorsal horn AS neurons. Together, these experiments reveal the cellular and molecular mechanisms of AS neuron specification and unravel an array of AS-enriched mRNAs.

### Phox2a is required for AS neuron development

Given the requirement of Phox2a for normal locus coeruleus development (Morin et al., 1997), we hypothesised that its loss may also impact the development of spinal Phox2a neurons. As *Phox2a* null mice do not survive beyond birth, we used the *Hoxb8*^*Cre*^ driver to ablate *Phox2a* selectively in the caudal spinal cord (Fig. S6A (Witschi et al., 2010)), producing Phox2a^cKO^ (*Hoxb8*^*Cre*^; *Phox2a*^*f/f*^) and control (*Phox2a*^*f/f*^ or *Hoxb8*^*Cre*^; *Phox2a*^+*/*+^) adult mice. To determine whether Phox2a plays a role in AS connectivity, we genetically labelled spinofugal axons by crossing the axonal tdTomato Cre reporter *R26*^*LSL-tdT*^ into Phox2a^cKO^ and control lines, generating, respectively, (*Hoxb8*^*Cre*^; *Phox2a*^*f/f*^; *R26*^*LSL-tdT*^) and (*Hoxb8*^*Cre*^; *Phox2a*^+*/*+^; *R26*^*LSL-tdT*^) mice. While most spinofugal target nuclei appeared to be normally innervated in *Phox2a*^*cKO*^ mice (Fig. S6B), a dramatic loss of tdT axons in the pBil was observed (Fig. 6A, B). To examine this defect in more detail, we injected a retrograde tracer into the pB of control and Phox2a^cKO^ mice (Fig. S6C), and quantified the number of tracer+ lamina I and lamina V/LSN neurons in the cervical and lumbar spinal cord which are, respectively, rostral to and within the *Hoxb8*^*Cre*^ expression domain. In the upper cervical spinal cord, we found a similar number of Tracer-labelled lamina I and lamina V/LSN neurons in both groups (Fig. S6D-F). In contrast, in the caudal spinal cord, while Tracer-labelled lamina I neuron number was unchanged in Phox2a^cKO^ mice (Fig. 6C, D), the number of Tracer+ ipsilateral and contralateral lamina V/LSN neurons was dramatically decreased (Fig. 6C, E). To investigate cellular changes leading to these connectivity phenotypes, we analysed *Phox2a* mRNA in e16.5 control and Phox2a^cKO^ embryos. *Phox2a* mRNA could be detected in Phox2a^cKO^ embryos, likely due to persistence of the truncated *Phox2a* transcript, and revealed similar numbers of Phox2a neurons in control and Phox2a^cKO^ e16.5 mice (12.6±3.7 cells/section, n=4, and 16.6±1.5 cells/section, n=4, respectively, p=0.089 unpaired t-test) although Phox2a^Deep^ neurons were displaced medially (Fig. 6H, I), arguing that these cells do not die but are dysfunctional. *Phox2a* mRNA expression in Phox2a^cKO^ mice appeared elevated compared to controls, suggesting that Phox2a may negatively regulate *Phox2a* expression.

**Figure 6:**
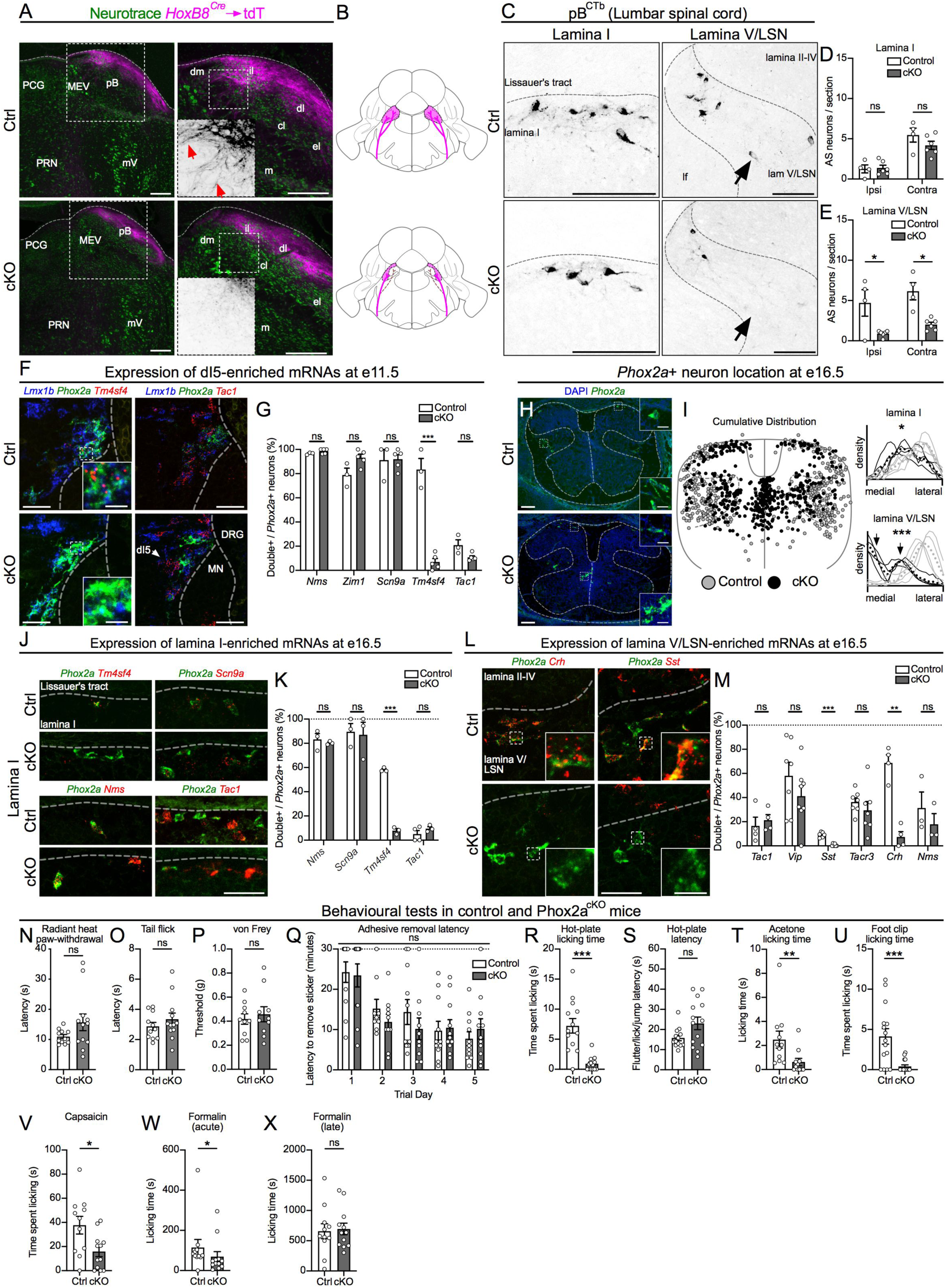
Phox2a is required for AS neuron development and function. (A) The parabrachial nucleus of control (*Hoxb8*^*Cre*^; *Phox2a*^+*/*+^; *R26*^*LSL-tdT/*+,^ Ctrl, top row) and Phox2a^cKO^ (*Hoxb8*^*Cre*^; *Phox2a*^*f/f*^; *R26*^*LSL-tdT/*+,^ cKO, bottom row) adult mice, depicting spino-parabrachial axons labelled via *Hoxb8*^*Cre*^-driven axonal tdT expression and counterstained with Neurotrace (magenta). Left panels show low power view, magnified in the right panels, with insets depicting tdT axons (black signal, arrows) (B) Diagram depicting the loss of spinal afferents (magenta) to the pB in Phox2a^cKO^ mice. (C–E) Spinal neurons labelled by CTb injections in the pB, in lamina I (left column of C, quantified in D) and lamina V/LSN (right column of C, quantified in E), in control (top row) and Phox2a^cKO^ (bottom row) adult mice. (F–G) Expression of *Phox2a, Lmx1b* and dI5-enriched *Tm4sf4, Tac1, Nms, and Scn9a* mRNAs in control (top row) and Phox2a^cKO^ (bottom row) e11.5 spinal cords. (G) Percent of *Phox2a*+ cells co-expressing each mRNA in control and Phox2a^cKO^ mouse embryos. See Fig. S6 for additional images. (H, I) Distribution of *Phox2a*+ neurons (green) in e16.5 control and Phox2a^cKO^ mouse embryos, co-stained with DAPI (blue). (H) Insets show individual *Phox2a*+ cells magnified. (I) Individual *Phox2a* cell locations in 3-5 sections (10 μm) of 4 control (grey circles) and 4 Phox2a^cKO^ (black circles) e16.5 spinal cords. Coordinates are normalised to the width and height and cells plotted on an idealised spinal cord, suggesting an abnormally centralised location of *Phox2a*+ cells in Phox2a^cKO^ embryos. The right panels are density plots of the normalized mediolateral distribution of lamina I and lamina V/LSN *Phox2a* neurons, with mediolateral position normalised to spinal cord width. Individual lines represent single animals, dotted lines represent mean distribution of 4 animals, grey lines represent control embryos and black lines represent Phox2a^cKO^ *Phox2a*+ cells. Black arrows point to the bimodal distribution of Phox2a^cKO^ *Phox2a*+ cells in deep laminae. (J, K) Expression of selected dI5-enriched *Nms, Scn9A, Tm4sf4* and *Tac1* mRNAs in lamina I *Phox2a*+ neurons of e16.5 control (top row) and Phox2a^cKO^ (bottom row) embryos. (K) Quantification of the percent of *Phox2a*+ neurons expressing selected mRNAs. (L, M) Expression of *Tac1, Vip, Sst, Tacr3, Crh, Nms* mRNAs encoding neuropeptides and neuropeptide receptors in lamina V/LSN *Phox2a*+ neurons of e16.5 control (top row) and Phox2a^cKO^ (bottom row) embryos. See Fig. S6 for additional images. (M) Quantification of the percent of lamina V/LSN *Phox2a*+ neurons expressing selected mRNAs. (N–P) Measures of spinal reflexes in control and Phox2a^cKO^ mice: (N) Radiant heat paw-withdrawal assay; n=11 control, n=12 Phox2a^cKO^. (O) Hot-water Tail Flick assay; n=11 control, n=12 Phox2a^cKO^. (P) von Frey test; n=10 control, n=10 Phox2a^cKO^. (Q) Adhesive removal latency in control and Phox2a^cKO^ mice over 5 days. Failure to remove the adhesive within 30 minutes is recorded as a 30-minute latency. n=11 control, n=10 Phox2a^cKO^. (R–X) Measures of supraspinal nociception. (R– U) Responses to non-injected noxious stimuli: Time spent licking (R) and latency to any response (S) during the hot-plate test; n=13 control, n=14 Phox2a^cKO^. (T) Time spent licking after hind paw application of acetone, n=11 control, n=11 Phox2a^cKO^. (U) Time spent licking after toothless alligator clip application to hind paw; n=15 control, n=16 Phox2a^cKO^. (V–X) Time spent licking after hind paw injection of noxious substances: (V) capsaicin; n=11 control, n=12 Phox2a^cKO^; (W, X) formalin (W, acute phase, X, late phase); n=11 control, n=12 Phox2a^cKO^. Data are represented as mean ± SEM. Numbers: A) n=4 control, n=4 Phox2a^cKO^ adult mice; (C–E) n=4 control, n=6 Phox2a^cKO^ adult mice; (F, G) n=3 control, n=5 Phox2a^cKO^ e11.5 mice; (H, I) n=4 control, n=4 Phox2a^cKO^ e16.5 mice; (J, K) n=3-4 control, n=3 Phox2a^cKO^ e16.5 mice; (L, M) n=3-8 control, n=3-8 Phox2a^cKO^ e16.5 mice; (N–X) Numbers described above. Statistics: (D, E) Two-way ANOVA with Tukey’s multiple comparisons test, (I) Unpaired t-test, (G, K, M) multiple t-tests using Holm-Sidak method, (Q) mixed-effects analysis with Sidak’s multiple comparisons test and Mann Whitney test (N, O, P, R, S, T, U, V, W, X). ns: non-significant, *: p<0.05, **: p<0.01, ***: p<0.001. Scale bars: (A) 250 μm, (C) 50 μm, (H) 100 μm, insets 20 μm, (F, J, L) 50 μm, insets 10 μm. Images in Fig. 5CC,DD have been re-used in Fig. 6F and images in Fig. 5 II, JJ, KK, LL have been re-used in Fig. 6J. Data from Fig. 5GG, NN representing the above images are re-used in Fig. 6G and K respectively. Validation of enriched mRNAs from RNA-Seq analysis in control embryos (Fig. 5) and comparison of enriched mRNAs between control and Phox2a^cKO^ embryos (Fig. 6) were performed as a single experiment. Abbreviations: cl (central lateral parabrachial nucleus), dl (dorsal lateral parabrachial nucleus), dm (dorsal medial parabrachial nucleus), el (external lateral parabrachial nucleus), il (internal lateral parabrachial nucleus), DRG (dorsal root ganglion), m (medial parabrachial nucleus), MEV (midbrain trigeminal nucleus), MN (motor neurons), mV (trigeminal motor nucleus), pB (parabrachial nucleus), PCG (pontine central gray), PRN (pontine reticular nucleus).

To understand the molecular underpinnings of these phenotypes, we compared the expression of Phox2a AS neuron-enriched mRNAs (Fig. 5) in e11.5 and e16.5 Phox2a^cKO^ and control mice, in neurons expressing *Phox2a* mRNA. Of these, only the expression of *Tm4sf4*, a gene encoding a protein implicated in cellular differentiation, was affected by *Phox2a*^*cKO*^ mutation in e11.5 dI5 neurons and e16.5 lamina I neurons (Fig. 6F, G, J, K, S6G). Given the peptidergic heterogeneity of lamina V/LSN neurons (Leah et al., 1988), we also monitored the expression of neuromodulatory peptides and receptors in presumptive Phox2^Deep^ neurons in e16.5 Phox2a^cKO^ and control spinal cords. Indeed, expression of genes encoding lamina V/LSN-enriched peptides *Sst* (Somatostatin) and *Crh* (Corticotrophin-releasing hormone) was reduced in Phox2a^cKO^ mice, while the expression of other Phox2a neuron-enriched transcripts such as *Tac1* (expressed in some Phox2a^Deep^ neurons), *Vip* (encoding Vasoactive Intestinal Peptide), *Tacr3* (encoding Tachykinin Receptor-3) and *Nms* (encoding Neuromedin S) co-expressed in some Phox2a^Deep^ neurons) remained unaffected (Fig. 6L, M, S6K, L). Phox2^Deep^ neurons also expressed *Tacr1* (NK1R), which is a known AS-enriched transcript, but not the interneuronal peptide *Cck* (Fig. S6M). We also monitored the expression of selected neuromodulatory genes in lamina I neurons and found elevated expression of *Vip* in Phox2a^cKO^ mice (Fig. S6J). Also, *Phox2a* mutant *Phox2a* neurons maintain their excitatory identity through the expression of *Slc17a6* or *Slc32a1* mRNAs encoding, respectively, neurotransmitter transporters vGluT2 and vGAT (Fig. S6H). Consistent with this, expression of spinal peptides associated with inhibitory neurons such as *Gal* (encoding Galanin), *pDyn* (encoding Dynorphin) and *pNoc* (encoding Nociceptin) were expressed sparsely among Phox2^Deep^ neurons (Fig. S6M). Together, these results demonstrate that Phox2a is essential for normal axonal connectivity and migration of AS neurons, as well as their transcriptional identity.

### Spinal *Phox2a* loss impairs supraspinal nocifensive behaviours

Given the central role of the AS in supraspinal nociceptive signal relay, we reasoned that defects in spino-parabrachial connectivity and Phox2^Deep^ neuromodulatory peptide expression in Phox2a^cKO^ mice might result in impaired nocifensive behaviours that are evoked by supraspinal circuits, with minimal effect on spinally mediated behaviours. Indeed, spinal-level thermal (radiant heat paw-withdrawal, Fig. 6N, hot water tail-flick, Fig. 6O) and mechanical assays (von Frey test, Fig. 6P), did not reveal any differences between control and Phox2a^cKO^ mice. However, using a battery of behavioural assays requiring supraspinal transmission of noxious information, significant differences between control and Phox2a^cKO^ mice emerged. Thermal preference to innocuous and noxious temperatures (Fig. S6T–V) and behaviours evoked by innocuous touch in the adhesive removal test (Fig. 6Q, S6N) were not affected by the *Phox2a*^*cKO*^ mutation. In contrast, Phox2a^cKO^ mice showed deficits in hind paw licking evoked by noxious stimuli – a nocifensive behaviour requiring ascending spinal projections. When mice were placed on a 53 **°**C hot-plate, which evokes licking of the hind paws Phox2a^cKO^ mice spent significantly less time licking compared to controls (Fig. 6R, 6SO). One of the common dependent measures in the hot-plate test is hind paw flutter/shake incidence, a spinal reflex measure, as well as latency to any behaviour (hind paw flutter, licking, or jumping) which was not different between the experimental groups (Fig. 6S, S6P). Though the frequency of jumping (escape) behaviours in the hot-plate test was not different (Fig. S6Q), 4/13 control mice attempted escape versus 1/14 Phox2a^cKO^ mice. Neither control nor Phox2a^cKO^ mice displayed nocifensive behaviours in the cold-plate test (Fig. S6R). Phox2a^cKO^ mice also spent less time licking their hind paw cooled with acetone (Fig. 6T, S6S) as well as following noxious mechanical stimulation (Fig. 6U). Furthermore, Phox2a^cKO^ mice also exhibited less licking of hind paws injected with TRPV1 and TRPA1 agonists capsaicin and formalin, respectively, although the late/tonic phase of post-formalin injection licking was not affected (Fig. 6V–X). Together these results show that a loss of Phox2a during development dramatically disrupts AS neuron innervation of the pB and their molecular differentiation and concomitantly affects supraspinal aspects of a variety of nocifensive behaviours associated with AS function.

### Phox2a neuron molecular identity is conserved in the developing human spinal cord

Given that many classical insights into AS function arose from clinical observations, and that little is known about the molecular identity of human spinal neurons, we wondered whether Phox2a expression in the developing human spinal cord might allow insight into human nociception. We thus examined the expression of Phox2a protein and that of dorsal horn neuronal markers Pax2, Lmx1b, Lbx1, Tlx3, Pou4F2, and the nociceptive afferent marker TrkA, in human spinal cords at developmental ages similar to mouse mid-gestation: two at gestational week (G.W.) 7.3, and one each at G.W. 7.4, 8.0 and 8.4, three of which are depicted here ((Altman and Bayer, 2001) Fig. 7A; S7). At G.W. 7.3 Phox2a neurons (identified using a commercial Phox2a antibody, Fig. S7C) were found in the superficial dorsal horn adjacent to TrkA+ fibers (Fig. 7B’, S7A, S7B), in deeper laminae (Fig. 7B’’) and near the roof plate (Fig. 7B’’’), resembling the location of mouse Phox2a^LamI^ and Phox2a^Deep^ neurons. Human spinal Phox2a neurons co-expressed Lmx1b, Lbx1, but not Pax2, or Tlx3 (Fig. 7B, S7A, S7B). As in mouse spinal cords, human Phox2a expression appeared weaker in older spinal cords (G.W. 8.4, Fig. 7A, S7B) and *Phox2a* mRNA was not detected in human cords at later gestational ages (between G.W. 15–20, S. R. and A. K., unpublished observations). Together, these data suggest that the spinal Phox2a neuron developmental program is evolutionarily conserved and that Phox2a expression is a molecular feature of developing human AS neurons.

**Fig. 7:**
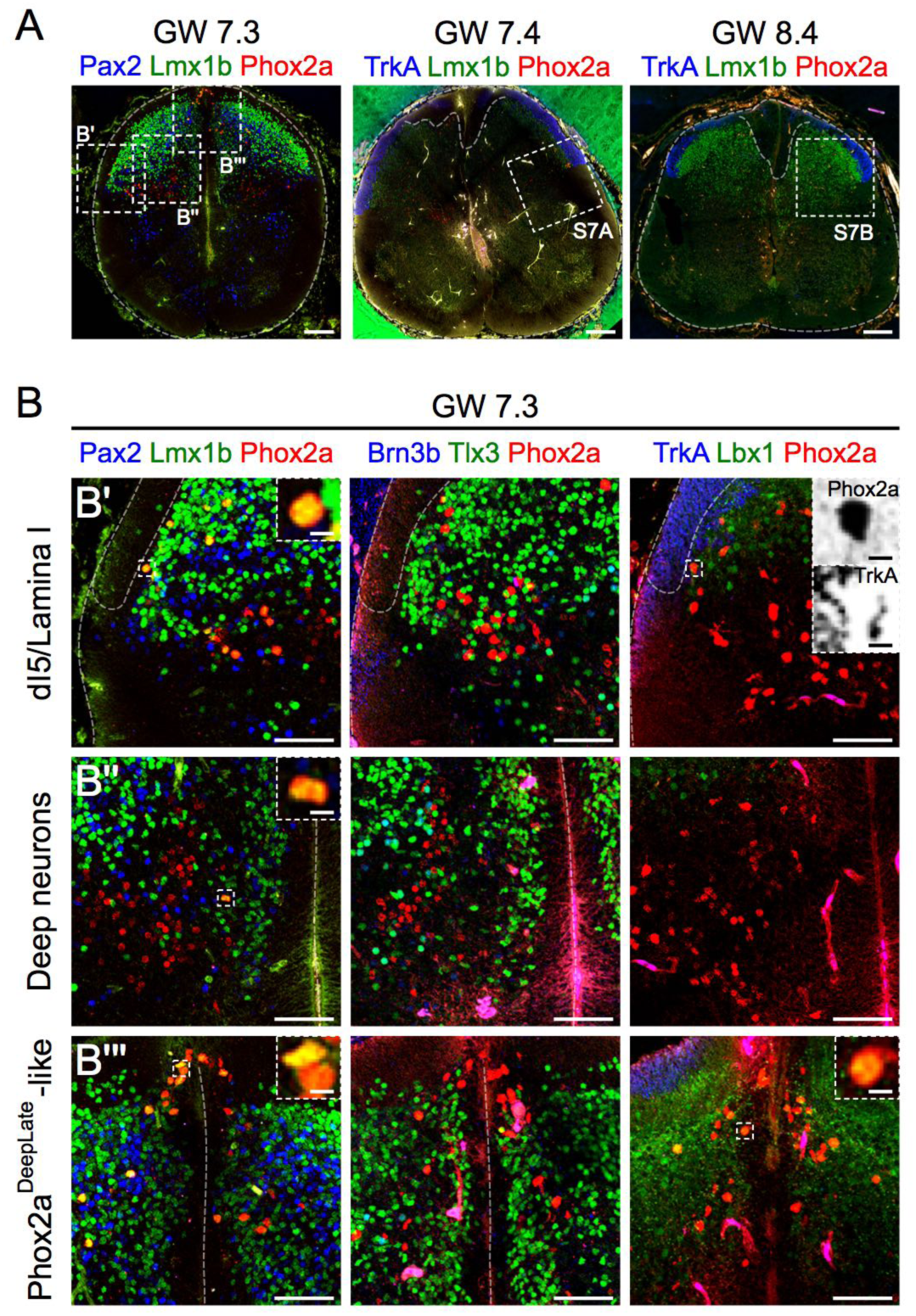
Phox2a neuron molecular identity is conserved in the developing human spinal cord. (A) Sections of G.W. 7.3-8.4 human spinal cords showing Phox2a, Lmx1b, TrkA and Pax2 expression. Location of higher magnification panels is shown in boxed regions. (B) Phox2a, Lmx1b, Pax2, Pou4F2, Tlx3, Lbx1 and TrkA expression in the G.W. 7.3 spinal cord, demonstrating co-labeling of Phox2a neurons with dorsal horn markers Lmx1b and Lbx1, but not with Pax2, Pou4F2 or Tlx3, in a Phox2a^LamI^/Phox2a^DeepEarly^-like cluster (B’), the deep dorsal horn (B’’) and a Phox2a^DeepLate^-like cluster near the roof plate (B’’’). Top row right panel shows apposition of Phox2a cells with TrkA+ sensory afferents, similar to that in mouse (Fig. 4). Insets show higher magnification of boxed regions (TrkA and Phox2a channels split and inverted). All experiments used the Abcam Phox2a antibody. See Fig. S7 for antibody specificity controls. Numbers: (A) Three human embryonic spinal cords (G.W. 7.3, 7.4 and 8.4 are represented here. (B) One G.W. 7.3 human embryonic spinal cord (from A) is represented here. Scale bars: (A) 200 μm, (B) 100 μm, Insets in (B’, B’’ and B’’’) 10 μm.

## Discussion

The anterolateral system (AS) is critical for the relay of nociceptive signals from the periphery to the brain yet, despite many years since its discovery, the molecular identity of AS neurons and their precise function remain obscure. Here we present evidence that essentially all spinal neurons that express Phox2a during their development innervate supraspinal targets and constitute a major tributary of the AS. Phox2a loss results in defects in spino-parabrachial connectivity and supra-spinal nocifensive behaviours. Furthermore, Phox2a neurons with molecular profiles identical to those in mice exist in the developing human spinal cord, arguing for an evolutionary conservation of this canonical pain relay pathway. Together, our observations reveal a rich developmental heterogeneity of AS neurons and provide insights into a molecular logic that underlies their functions.

### Diversity of AS neuron development revealed by Phox2a expression

Nearly all spinal Phox2a neurons can be retrograde labelled from the VPL thalamus and the pB and, as certain experiments resulted in virtually all spinal Phox2a neurons taking up retrograde tracer, we assume that their unlabelled fraction is a function of labelling efficiency or innervation of non-VP or pB brain targets. We thus propose that Phox2a is a genetic marker of AS neurons, allowing insights into the cellular and molecular mechanisms that produce their diversity. We classified AS neuron heterogeneity into at least three distinct and sequentially generated populations of spinal Phox2a neurons arising from the dI5 spinal progenitor domain: Phox2a^LamI^, Phox2a^DeepEarly^ and Phox2a^DeepLate^. Contrary to the notion that spinal neurons are born in a ventral to dorsal order (Nornes and Carry, 1978), superficial dorsal horn Phox2a neurons are born concurrently with motor neurons, as suggested recently for spinofugal neurons (Nishida and Ito, 2017). Ascl1 (expressed in dI5 progenitors) and Ptf1a (expressing in dI4 progenitors) were previously shown to, respectively, promote and inhibit dI5 neuron fates (Glasgow et al., 2005; Helms et al., 2005). Our data demonstrate that this may occur via direct action at a distal *Phox2a* enhancer defined in this study. The stereotyped birth order of Phox2a AS neurons raises the possibility that it is orchestrated by transcription factors involved in temporal competence of Ascl1-expressing progenitors, as in the cerebral cortex and retina (Kohwi and Doe, 2013).

Following birth and early specification, Phox2a^LamI^, Phox2a^DeepEarly^ and Phox2a^DeepLate^ AS neurons migrate along distinct trajectories. Phox2a^LamI^ neurons move tangentially to the surface of the developing dorsal horn, in contrast to radial trajectories of Phox2a^Deep^ neurons. The contacts between afferent axons and Phox2a^LamI^ neurons, and their implied importance for normal Phox2a^LamI^ migration suggest a developmental interplay between afferent sensory axons and their spinal neuron targets. One consequence of this interaction may be the settling of lamina I neurons in somatotopic order corresponding to their dermatome-specific sensory afferents (Willis et al., 1974). Consistent with their sparse sensory afferent innervation, Phox2a^Deep^ neuron position is unaffected by the loss of primary afferents. At the molecular level, the neuronal migration cue Reelin is likely mediating the radial migration of Phox2a^Deep^ neurons since its intracellular signalling effector Dab1 is required for normal positioning of lamina/LSN neurons (Yvone et al., 2017). Netrin signalling likely coordinates the interplay between sensory afferents and Phox2a^LamI^ neurons since its expression in the nascent dorsal horn prevents the premature ingrowth of primary afferents expressing the netrin1 repulsive receptor Unc5c (Watanabe et al., 2006), and the netrin1 attractive receptor DCC is required for the normal entry of Phox2a^LamI^ neurons into the dorsal horn (Ding et al., 2005).

Molecular profiling of early (e11.5) AS neurons under the assumption that they are an early-born dI5 Lmx1b-expressing cohort, reveals at least two distinct AS precursor populations: *Phox2a*+ cells and a complementary population of *Lmx1b*+ *Tac1*+ cells that likely give rise to the recently identified Tac1-expressing AS neurons (Huang et al., 2019). While no Tac1 dI5 neuron-enriched genes emerged from our analysis, we have identified Phox2a AS neuron-enriched transcripts and proteins with developmental and neuronal physiology functions. *Tm4sf4* and Shox2 are involved in cell fate specification (Anderson et al., 2011), while the axon guidance receptors DCC and Robo3 are critical for the commissural projection of spinal neuron (Fazeli et al., 1997; Sabatier et al., 2004), and in particular, that of spino-thalamic neurons (da Silva et al., 2018). *Pdzrn3*, a E3-ubiquitin ligase involved in Wnt receptor signalling, may contribute to the elaboration of long axons characteristic of AS neurons, given that many long axons tracts require the Wnt receptor Frizzled3 (Hua et al., 2014). Despite Phox2a^LamI^ and Phox2a^Deep^ neuron identities diverging through the expression of genes associated with neuronal function, nearly all Phox2a AS neurons share a glutamatergic identity. The function of Phox2a^LamI^ neurons could be modulated by a host of co-expressed factors, such as the peptides Nms, Crh, Sst, dynorphin, Vip or the acetylcholine receptor–binding neurotoxin Lypd1, as well as the alpha subunit of the Nav1.7 channel encoded by the *Scn9a*, known for its role in pain insensitivity syndromes in humans (Cox et al., 2006).

### Phox2a is required for the terminal differentiation of AS neurons

*Phox2a*-expressing neurons are present in normal numbers in neonatal Phox2a^cKO^ spinal cords suggesting that Phox2a is not required for their early specification or survival. However, *Phox2a* mutation results in the loss of *Tm4sf4* and gain of VIP expression in Phox2a^LamI^ neurons, indicating that Phox2a is required for their molecular differentiation, and thus possibly their function, despite apparently normal target connectivity. In contrast, nearly 75% of lamina V/LSN AS neurons fail to innervate the pB in Phox2a^cKO^ mice. This indicates that Phox2a is expressed in and required for the normal development of a vast majority of lamina V/LSN neurons, in agreement with our observation that *Phox2a*^*Cre*^ under-reports Phox2a expression in many of these neurons. One possibility is that the aberrant Phox2a^Deep^ neuron position in Phox2a^cKO^ mice, similar to that observed for LSN neurons in Reelin-deficient mice (Wang et al., 2012; Yvone et al., 2017), could impact their target connectivity or stability. Together, with the observation that *Phox2a* mutation also results in the loss of neuropeptide expression in Phox2a^Deep^ neurons, these defects argue that Phox2a specifies the terminal differentiation of AS neurons, and its absence likely impairs their function.

### Phox2a AS neuron function in supraspinal nociception

Adult Phox2a AS neuron morphologies and laminar organisation are typical of nociceptive AS neurons such that Phox2a^LamI^ neurons are likely a subset of the nociceptive-specific and somatotopically-organised lamina I AS projection neurons, while Phox2a^Deep^ neurons are likely a subset of the wide dynamic-range lamina V AS projection neurons, with the neuropeptide-expressing LSN neurons implicated in deep tissue nociception (Keay and Bandler, 2002). Spinal Phox2a neurons also innervate many of the principal AS brain targets involved in nociception, including the VPL thalamus, pB and the periaqueductal gray. Phox2a^cKO^ mice exhibit disrupted spinofugal connectivity and deficiencies in nocifensive behaviours associated with supraspinal circuit functions, in line with the requirement of the AS in relaying nociceptive information to the brain. Despite thermosensation relay being a feature of AS neurons (Hyndman and Wolkin, 1943), the apparently normal temperature preference of Phox2a^cKO^ mice may result from *Hoxb8*^*Cre*^ expression omitting the upper cervical spinal cord (Witschi et al., 2010) which receives thermal information from the forelimb and neck. In contrast, the normal hind limb adhesive tape-evoked behaviours in Phox2a^cKO^ mice are consistent with the notion that fine touch sensation is not a function of the AS (Hyndman and Wolkin, 1943). Despite having a large population of aberrantly-developed caudal AS neurons, Phox2a^cKO^ mice have normal spinal nocifensive reflexes indicating that both local reflex circuitry and the descending pathways that modulate these behaviours (Ren and Dubner, 2009) do not depend on normal AS function.

Phox2a^LamI^ neurons have been proposed to transmit sensory-discriminative information, and their function is likely impaired in Phox2a^cKO^ mice, but the study of this AS function is constrained by the motivational-affective drive of behavioural measures of sensory-discriminative nociceptive function. Indeed, Phox2a^cKO^ mice have a reduction in the frequency and duration of nociception-related behaviours evoked by the relay of spinal nociceptive signals to the brain, suggesting that the transmission of motivational-affective information is likely carried out by Phox2a AS neurons. In Phox2a^cKO^ mice, lamina V/LSN neuron innervation of the pB is severely reduced, consistent with the notion that lamina V/LSN AS neurons convey noxious motivational information through the spino-pBil-medial thalamus pathway that impinges on the pre-frontal cortex (Bourgeais et al., 2001). At the molecular level, *Phox2a* mutation also causes decreased expression of mRNAs encoding neuropeptides Sst (Leah et al., 1988) and Crh, normally enriched in Phox2a^Deep^ neurons of lamina V/LSN. Given the role of Crh in stress responses, Crh-expressing Phox2a^Deep^ neurons may convey motivational information linked to noxious stimuli.

Recent experiments point to another genetically defined component of the AS: an intersectional genetic ablation of Tac1 neurons, some of which are spino-parabrachial AS neurons, result in behavioural deficits similar to those in Phox2a^cKO^ mice (Huang et al., 2019). This could be explained by ∼20% of Phox2a^Deep^ neurons expressing Tac1 and is in line with lamina V/LSN neurons transmitting the emotional-motivational aspect of nociception, although it is unclear whether all Tac1 AS neurons are a subset of the Phox2a^Deep^ neuron population. Another possibility is that Tac1 non-AS interneurons ablated by the intersectional approach regulate Phox2a^Deep^ neuron function. Some distinctions may be made between Tac1 and Phox2a neurons, as deficits in capsaicin-evoked licking are seen in Phox2a^cKO^ mice but not in Tac1-ablated mice; thus, Phox2a neurons relaying noxious heat or chemical stimuli may be Phox2a^LamI^ neurons that are distinct from Tac1 lamina I AS neurons.

### The molecular logic of the anterolateral system

Supraspinal Phox2a lineage-derived neurons exist in a variety of autonomic circuits raising the question of whether these may be functionally intertwined with Phox2a AS neurons. Two lines of thought shed some light on this: firstly, Phox2a and its closely-related transcription factor Phox2b, specify the development of neurons afferent to medullary visceral reflex circuits that control many autonomic functions implicated in homeostasis (Brunet and Pattyn, 2002). Our genetic tracing experiments reveal that Phox2a AS neurons participate in this connectivity logic by innervating brain stem pre-autonomic regions such as the NTS and CVLM, as well as higher autonomic regulatory regions such as the pB. Secondly, because pain motivates behaviours that correct homeostatic changes, it has been proposed as a “homeostatic emotion” (Craig, 2003a). In light of this, the AS can be viewed as a pathway signalling deviations from homeostasis, such as changes in skin temperature, or the presence of noxious or pruritogenic stimuli, to brain regions that trigger compensatory autonomic responses (e.g.: CVLM) or drive compensatory behavioural responses such as licking or scratching (e.g.: pB). Given this, Phox2a AS neurons may specialize in transmitting somatic sensations with a motivational character such as cutaneous and deep pain, thermosensation, itch, visceral pain, nausea, and sexual arousal, all of which are abolished by anterolateral cordotomy in humans (Hyndman and Jarvis, 1940; Hyndman and Wolkin, 1943).

Our results suggest that the molecular identity of mouse Phox2a AS neurons is conserved in the developing human spinal cord, pointing to a conserved molecular logic of somatosensory circuit development, supported, in part, by the expression of *PHOX2A* in the human locus coeruleus (Fan et al., 2018). A genetic proof of this idea remains out of reach because of the lack of obvious nociceptive or autonomic deficits in humans with *PHOX2A* mutations, which may be due to hypomorphic alleles (Nakano et al., 2001). Nevertheless, *PHOX2A* is a compelling molecular marker of human AS neurons and given the effectiveness of cordotomy as a crude treatment of intractable chronic pain, a molecularly-defined inactivation of a Phox2a AS neuron subpopulations could be its more refined iteration.

## Acknowledgements

We thank Meirong Liang, Julie Cardin, Qinzhang Zhu and Colleen Barrick for technical assistance, Laura Kus (GENSAT) for advice on transgenic mouse design, Caroline Grou and Virginie Calderon for bioinformatics analyses, J.-F. Brunet for the Phox2a and Phox2b antiserum and advice, Carmen Birchmeier for Lmx1b, Tlx3 and Lbx1 antisera, Jay Bikoff for the Pou6F2 antiserum, Laskaro Zagoraiou for Shox2 antiserum, Hanns Ulrich Zeilhofer for the *Hoxb8*^*Cre*^ mice, and, Jeff Mogil, Yves de Koninck and Stefano Stifani for discussions, and Jean-François Brunet, Adam Hantman, Denis Jabaudon, Jeff Mogil, Michel Cayouette, Sonia Paixao, Samuel Ferland and Feng Wang for comments on the manuscript.

R.B.R. was a recipient of a Ph.D. studentship from Fonds de recherche du Québec –Santé. S.R. received a studentship from McGill University’s Healthy Brains, Healthy Lives (HBHL) initiative supported, in part, by Canada First Research Excellence Fund. J.E.J was supported by the National Institutes of Health R37 HD091856. A.C. was supported by Agence Nationale pour la Recherche and INSERM (transversal program HuDeCa). L.T. was supported by the Intramural Research Program of NCI, NIH. A.K. and M.K. were funded by Operating and Project Grants from the Canadian Institutes of Health Research (to A.K.: PJT-162225, MOP-77556, PJT-153053, PJT-159839; to M.K.: MOP-127110 and PJT-162143).

## Author contributions

Conceptualisation, R. B. R. and A. K.; Methodology, R. B. R., B. M., R. B., C. S., J. E. J., A. D., M. K.; Validation, R. B. R.; Formal Analysis, R. B. R. and B. M; Investigation, R. B. R., F. B. B., B.M., S. R., R.B., C. S., W. S. T., and M. B.; Resources, R. B. R., A. D., M. K., L. T., Y. G., M. G., and A. K.; Data Curation, R. B. R., B. M., J. E. J.; Writing Original Draft, R. B. R.; Writing – Review & Editing, R. B. R., F. B. B., S. R., M. K., L. T., J. E. J., M. K., A. C., and A. K.; Visualisation, R. B. R., B. M., J. E. J.; Supervision, J. E. J., M. K., A. C., A. K.; Project Administration, A. K.; Funding Acquisition, J. E. J., M. K., A. C., A. K.

## STAR Methods

### LEAD CONTACT AND MATERIALS AVAILABILITY

Further information and requests for resources and reagents should be directed to and will be fulfilled by the Lead Contact, Artur Kania

### EXPERIMENTAL MODEL AND SUBJECT DETAILS

#### Mouse lines and *Phox2a*^*Cre*^ mouse line generation

Adult male and female mice, between 6–19 weeks of age, were used in this study. Sex ratios were kept as close to 1:1 as possible in all experiments, though not all experiments had the power to distinguish sex differences. Mice were kept on a 12 hour light : 12 hour dark cycle (light 6:00-18:00) with food and water provided *ad-libitum*. All procedures (except those involving *TrkA*^*-/-*^, *Ptf1a*^*CRE*^ and *Ascl1*^*GFP*^ mice) were approved by the IRCM Animal Care Committee, using regulations and guidelines provided by the Canadian Council for Animal Care (CCAC). *TrkA*^*-/-*^ mouse use was approved by the Committee of Animal Care and Use of the National Cancer Institute, while the use of *Ptf1a*^*CRE*^ and *Ascl1*^*GFP*^ mouse lines (maintained on a mixed background of ICR and C57Bl/6), was approved by the Institutional Animal Care and Use Committee at University of Texas Southwestern. Phox2a^Cre^ mice were generated at the IRCM where *Phox2a*-containing BAC RP23-333J21 (GENSAT, 2008) was modified by insertion of a Cre-PolyA sequence into the ATG site of *Phox2a* using GalK recombineering strategies (Warming et al., 2005). The Cre-containing BAC was injected into fertilized ova, and the resulting offspring were screened for genomic insertion of the BAC using Cre PCR. In total, we screened 230 pups, and were able to produce one founder from which all mouse lines containing *Phox2a*^*Cre*^ were derived. Genotyping was done by PCR for *Cre, FlpO, R26*^*LSL-tdT/*+^ (Ai14), *R26*^*FSF*^*-*^*LSL-tdT/*+^ (Ai65), *Phox2a*^*f/f*^ and *TrkA*^-/-^ as previously described (Glasgow et al., 2005; Kim et al., 2008). The *Ptf1a*^*CRE*^ mouse line replaces the coding sequence for *Ptf1a* with that for *Cre* recombinase (Kawaguchi et al., 2002) and the *Ascl1*^*GFP*^ (*Ascl1*^*tm1Reed*^/J) mouse strain replaces the coding sequence of *Ascl1* with that for *GFP* (Leung et al., 2007).

#### Generation of mice and mouse embryos

Mice containing the following transgenes or alleles were generated: *Phox2a*^*Cre*^; *R26*^*LSL-tdT/*+^, *Phox2a*^*Cre*^; *Cdx2*^*FlpO*^; *R26*^*FSF-LSL-tdT/*+^, and *Hoxb8*^*Cre*^, *Phox2a*^*f/f*^, *R26*^*LSL-tdT/*+^, *Ascl1*^*GFP/GFP*^, *Ptf1a*^*Cre/Cre*^, *TrkA*^*-/-*^, *Hoxb8*^*Cre*^; *Phox2a*^*f/f*^; *R26*^*LSL-tdT/*+^, by breeding parents bearing one or more of the necessary alleles/transgenes. Vaginal plugs were checked daily at 6:00am, and the day of plug detection was noted as embryonic day 0.5 (e0.5). Mothers were anesthetised with a 0.3 mL intra-peritoneal injection of Ketamine/Xylazine solution. Embryos were dissected in ice-cold 1x phosphate-buffered saline (1x PBS), transferred to 4% paraformaldehyde in 1x PBS (4 °C) and left to fix for two hours on a moving shaker (except *Ptf1a*^*Cre/Cre*^ and *Ascl1*^*GFP/GFP*^ embryos which were fixed for one hour). After fixation, embryos were washed briefly in 1x PBS, then cryoprotected in 30% sucrose for 1-2 days or until sunk. Embryos were harvested and fixed on the following embryonic days: *TrkA*^*-/-*^ on e14.5, *Ptf1a*^*Cre/Cre*^ and *Ascl1*^*GFP/GFP*^ both on e11.5 and e14.5, *Hoxb8*^*Cre*^; *Phox2a*^*f/f*^; *R26*^*LSL-tdT/*+^ on E11.5 and E16.5, and *Phox2a*^*Cre*^; *R26*^*LSL-tdT/*+^ on e9.5, e10.5, e11.5, e12.5, e13.0, e13.5, e14.5, e15.5, e16.5 and e18.5.

#### Acquisition of human embryonic spinal cords

Human embryos were obtained with the parent’s written informed consent (Gynaecology Hospital Jeanne de Flandres, Lille, France) with approval of the local ethic committee. Tissues were made available via the INSERM-funded Human Developmental Cell Atlas resource (HuDeCA) in accordance with the French bylaw (Good practice concerning the conservation, transformation and transportation of human tissue to be used therapeutically, published on December 29, 1998). Permission to use human tissues was obtained from the French agency for biomedical research (Agence de la Biomédecine, Saint-Denis La Plaine, France). Human embryo spinal cords were fixed by immersion for 12–24 hours in 4% paraformaldehyde in 0.12 M phosphate buffer, pH 7.4 (PFA) over night at 4 °C. Samples were cryoprotected in a solution of 10% sucrose in 0.12 M phosphate buffer (pH7.2), frozen in isopentane at 50 °C and then cut at 20 µm with a cryostat (NX70 Thermo Fisher). Spinal cords from five separate embryos were used in this study: two from G.W. 7.3, and one each from G.W. 7.4, 8.0 and 8.4.

## METHOD DETAILS

### Neuronal birthdating

Pregnant female mice were given an i.p. injection of BrdU on e9.5, e10.5, e11.5 or e12.5 and embryos were harvested and fixed at e11.5, e12.5, e13.5 or e16.5. The BrdU dose was 50 mg/kg for all time points except e9.5, where this dose produced ubiquitous BrdU+ immunoreactivity in the spinal cord and thus was reduced to 25 mg/kg.

### Stereotaxic surgery

Prior to surgery mice were given 1 mg/kg buprenorphine for analgesia, then anesthetised using a mixture of 5% isoflurane in oxygen and maintained using 2% isoflurane in oxygen. Eyes were coated in eye ointment to prevent drying during anesthesia. Prior to incision, the top of the head was shaved and decontaminated using an iodine solution. Mice were fitted into a stereotaxic frame with digital coordinate display and an incision was made longitudinally along the scalp to bare skull sutures. Injections were made via a hole drilled in the skull, which was made using medial-lateral and anterior-posterior coordinates for underlying brain regions as defined by the coronal Allen Brain reference atlas (Dong, 2008). Retrograde tracers (fluorogold or CTb-488) were injected using a 5 μl Hamilton syringe fitted with a pulled glass needle backfilled with mineral oil, which were injected in the VPL thalamus (coordinates AP -1.7, ML -2.0, DV -3.2), the MD thalamus (AP -1.25, ML -0.4, DV -3.2), or the parabrachial nucleus (AP -5.35, ML -1.4, DV - 3.05), identified using the coronal Allen Brain reference atlas (Allen Institute for Brain Science, 2004). Injection volumes of 500 μl (fluorogold, 2%) were injected into the VPL and 300 μl (CTb-488, 1%) into the MD thalamus or parabrachial nucleus. The needle was left in place for 5 minutes before slowly withdrawing to prevent reflux. The incision was then stitched together using silk sutures and mice were allowed to recover under a heating lamp before being returned to their home cage.

### Mouse behavioural assays

R. B. R. performed all behavioural assays, and was blinded to genotypes. R. B. R. and M. B. analysed video-recorded mouse behaviour, though each experimenter analyzed equal numbers of mice from each sex and genotype per assay. Mice of both sexes were used in each behavioural assay. Mice from control and Phox2a^cKO^ groups were always littermates and the same sex, to prevent confounding effects of litter versus sex. Control and Phox2a^cKO^ groups thus always contained an equal proportion of mice from each sex, and the proportion of male to female mice within groups was kept as close to 50% as possible, constrained only by the number of Phox2a^cKO^ mice generated (at an expected rate of 12.5% in a given litter). Mice were habituated in a dedicated mouse behaviour room for at least 30 minutes prior to onset of tests. Mice received no other treatments other than the test itself. Mice were habituated in a small plexiglass chamber measuring 4 cm long, 2.2 cm wide and 2.5 cm high for von Frey, radiant heat paw-withdrawal, acetone and adhesive removal tests. For the von Frey and acetone tests, the chambers were placed atop a perforated stainless steel floor due to the need for physical hind paw manipulations. For the radiant heat paw-withdrawal and adhesive removal test, the chambers were placed atop a transparent glass sheet. For all other assays mice were habituated in their home cages. When necessary, all behavioural tests were filmed using an iPhone SE except for the temperature preference assay, where the video camera included in the apparatus was used.

The **von Frey test** involved using a set of nylon filaments (0.008, 0.02, 0.04, 0.07, 0.16, 0.4, 0.6, 1.0, 1.4 g) to stimulate the hind paw plantar surface of each mouse in order to determine the median force which produces a withdrawal reflex. Mice were tested with a series of filaments using the “up-down” method of Dixon, as described previously (Chaplan et al., 1994; Mogil et al., 1999), with an inter-trial interval of at least 5 minutes.

The radiant heat paw-withdrawal (**Hargreaves)** test involved stimulating the hind paw plantar surface from below with a focused beam of light (set to 10% maximum intensity of the machine) and verifying latency to withdraw either hind paw. Each hind paw was stimulated eight times (16 total stimulations), and data was represented as the average of 16 withdrawal latencies, with an inter-trial interval of at least 2 minutes, performed as previously described (Hargreaves et al., 1988; Mogil et al., 1999).

The **hot water tail-withdrawal** test was performed as described previously (Mogil et al., 1999). Mice were placed in a small cloth pouch into which they entered voluntarily, the distal portion of the tail was dipped into a hot water bath maintained at 49±1 °C and the latency to withdraw the tail was recorded. Mice were tested three times with an inter-trial interval of at least 2 minutes, and data was represented as the average of 3 withdrawal latencies.

The **adhesive removal test** was performed as described previously (Bouet et al., 2009). Mice were tested on five consecutive days for the ability/motivation to remove an adhesive placed on the plantar surface of the hind paw. The adhesive was half of a 1.5 ml Eppendorf tube cap label, cut into a semicircle, and placed on the plantar surface. The latency to remove the label was recorded to the nearest minute, and these data were reported exactly as recorded (with only one test per day and no averaging between trials). If mice did not remove the adhesive within 30 minutes of the start of the test, latency was recorded as “30 minutes” for the purpose of data analysis, and mice were then returned to their home cage.

The **two-plate temperature preference assay** was performed as described previously (Minett et al., 2012). Two temperature-controlled metal plates were abutted together within a plexiglass enclosure. Mice were given the choice to travel between a probe temperature plate and a control temperature (always 30 °C) plate for 10 minutes and the time spent per plate, distance traveled per plate and transitions between plates were recorded via a video camera above the enclosure (included with apparatus) and analyzed automatically via the accompanying software. Mice were tested twice for each probe temperature, and data for time/distance/transitions were represented as the average of both trials. In order to prevent mice from associating one plate as the control plate, the control plate was switched for each trial. Moreover, between testing for different probe temperatures, the initial position of the control plate was switched with the probe plate to prevent mice from associating the order of trials with the location of the control plate. As well, to encourage mice to sample both plates, mice were placed randomly on either the control plate or the probe plate for the first trial, and this order was then switched for the second trial.

The **hot-plate test** was performed as described previously (Mogil et al., 1999), and the **cold-plate test** was performed using similar methods. Mice were placed within the hot-cold plate apparatus (IITC PE34) on a stainless-steel metal plate heated to 53±0.1 °C or cooled to 0±0.1 °C and were video-recorded from the side (with a mirror opposite the test chamber to view each side of the mouse) for 60 seconds at which point they were returned to their home cage. The latency to either lick the hind paw, flutter of the hind paw or to attempt to escape via jumping was recorded. Additional behaviours were recorded: total time spent licking either hind paw, total hind paw licking episodes, total jumps and total hind paw flutters. Mice were tested once, and data were represented directly based on behaviours recorded in one 60-second trial. Entirely different cohorts of mice were used for the hot and cold-plate tests respectively, to prevent behavioural adaptation to the test.

The **acetone test** was performed as described previously (Colburn et al., 2007). Briefly, the mouse’s hind paw was stimulated with a drop of acetone extruded from the blunt end of a 1ml syringe. Mice were recorded for 60 seconds following the application, and total time spent licking was recorded as well as the magnitude of behaviour on a 0–2 scale as reported previously (Colburn et al., 2007). Mice were stimulated 5 times, with an inter-trial interval of at least 5 minutes. Total licking time was reported as a sum of 5 trials, and the behavioural score (0–2) was reported as an average of 5 trials.

The **foot clip test** was performed as described previously (Huang et al., 2019). Briefly, a toothless mechanical clip was used to pinch skin on the plantar surface of the hind paw, and mice were placed in a plexiglass cylinder (dimensions) on the glass sheet used previously and video recorded from below for 60 seconds (this recording setup is identical to the following formalin and capsaicin tests). The total amount of time licking the clipped hind paw was recorded, and data is presented as the total time licking during the one trial.

The **capsaicin and formalin tests** (Mogil et al., 1999; Sakurada et al., 1992) were performed similarly – mice were injected with approximately 20 μl of capsaicin solution (1.5 μg/20 μl in 1x PBS) or formalin solution (2% in 1x PBS) in the plantar surface of the right hind paw using a standard 28G insulin syringe (BD) and video recorded from below for either 15 or 60 minutes respectively. Mice were tested only once on each test, with different cohorts of mice used for each respective test. Data were represented as time spent licking the injected hind paw. For formalin-injected animals, these data were analyzed separately acutely after injection (0–10 minutes) or chronically after injection (11–60 minutes).

### Tissue fixation, freezing and sectioning

Adult mice were first anesthetised with a 0.3 ml i.p injection of Ketamine/Xylazine solution (10 mg/ml Ketamine, 1 mg/ml Xylazine, in 0.9% saline). Transcardial perfusion was done with a peristaltic pump (Gilson miniPuls2). Mice were perfused with 10 ml of ice cold 1x PBS followed by 20 ml of ice cold 4% PFA in 1x PBS. Brains and spinal cords were dissected and post-fixed in 4% PFA in 1x PBS at 4 °C for two hours, washed briefly in 1x PBS, and acclimated to 30% sucrose for 1–2 days or until sunk. After cryoprotection, tissue was frozen in OCT Compound and cryosectioned at -22 °C. Tissue was cut into 25 μm sections for all experiments other than RNA Scope, in which case 10 μm sections were used, and those involving *Ptf1a* ^*CRE*^ and *Ascl1* ^*GFP*^ lines where 30 μm sections were used.

### Immunohistochemistry

For mouse tissue, sections were heated at 37 °C for 15 minutes prior to immunohistochemistry. Following this, sections were washed three times in 1x PBS for 10 minutes, blocked using a solution of 5% heat-inactivated horse serum (HIHS) and 0.1% Triton X-100 in 1x PBS (0.1% tPBS) for 30 minutes, and incubated with a primary antibody solution (in 1% HIHS, 0.1% tPBS) overnight at 4 °C. The following day, sections were again washed three times in 1x PBS for 10 minutes, and incubated with a secondary antibody solution (in 1% HIHS, 0.1% tPBS) at room temperature for 1 hour. Following this, sections were washed three more times in 1x PBS for 10 minutes and coverslipped using a Mowiol solution (10% Mowiol - Sigma, 25% glycerol). Slides were allowed to dry in the dark at room temperature and subsequently imaged using fluorescent microscopy. For immunohistochemistry involving the anti-BrdU antibody, two rounds of immunohistochemistry were done: the first round involved staining for RFP or Phox2a and the second round for BrdU with some modifications. Prior to the anti-BrdU primary antibody incubation, slides were treated in a 2 N hydrochloric acid solution at 37 °C for 30 minutes. Subsequently, slides were neutralized by washing in a Tris-buffered saline solution (pH 8.5, 50 mM Tris, 150 mM NaCl) for 10 minutes at room temperature, after which primary antibody incubation was done. BrdU immunohistochemistry proceeded in two steps, as acid denaturation of DNA reveals anti-BrdU epitopes but destroys RFP/Phox2a epitopes; however, acid denaturation does not destroy secondary antibody-conjugated fluorophores from the first round of immunohistochemistry. Immunohistochemistry on human tissue was performed on cryostat sections after blocking in 0.2% gelatin in PBS containing 0.25% Triton-X100 (Sigma). Sections were then incubated overnight with respective primary antibodies, all used at 1:500 dilutions, followed by 2 hours incubation in appropriate secondary antibodies.

### In situ-hybridisation (ISH)

ISH was done using RNA Scope® Multiplex Fluorescent v2 kits, according to manufacturer’s instructions. All experiments used Mm-Phox2a-C2 coupled to Opal™ 520, Mm-Lmx1b-C3 coupled to Opal™ 690, and all other candidate probes being compared to Phox2a (all Mm C1 probes) were coupled to Opal™ 570.

### *In-ovo* chicken electroporation and tissue processing

Fertilized White Leghorn eggs were obtained from the Texas A&M Poultry Department (College Station, TX, USA) and incubated for 48 hours at 39°C. The supercoiled reporter plasmid *ePhox2a-GFP* was diluted to 1.5 mg/mL in H_2_O/1X loading dye and injected into the lumen of the closed neural tube at stages Hamburger-Hamilton (HH) stages 13-15 (∼E2) along with either a *pMiWIII-Myc* epitope tagged plasmid serving as an electroporation control or the same plasmid containing the coding region of *Ascl1, Ptf1a*, or *Prdm13* (Hamburger and Hamilton, 1951). The injected embryos were then electroporated with 5 pulses of 25 mV each for 50 msec with intervals of 100 msec. Embryos were harvested 48 hours later at HH stages 22–23 (∼E4), fixed with 4% paraformaldehyde for 45 minutes, and processed for cryosectioning and immunofluorescence.

### Generation of reporter constructs and expression vectors

Previously published ChIP-seq data for *Ascl1, Ptf1a, Rbpj*, and *Prdm13* (Borromeo et al., 2014; Meredith et al., 2013; Mona et al., 2017) (GSE55840; GSE90938) were used to identify a putative enhancer for *Phox2a* in the chicken dorsal neural tube. A 851 bp region (chr7:101834344–101835194 from mm10) encompassing two peaks bound by Ascl1 and Ptf1a was cloned into the MCSIII GFP reporter cassette (*ePhox2a-GFP*). This reporter cassette contains the β*-globin* minimal promoter, a nuclear localised fluorescence reporter, and the 3’ cassette from the human growth hormone. The *ePhox2a* sequence was PCR amplified from ICR mouse DNA. *Prdm13*, ^*myc*^*Ptf1a*, and ^*myc*^*Ascl1* were expressed in the pMiWIII expression vector (Chang et al., 2013; Gowan et al., 2001; Matsunaga et al., 2001). All constructs were sequence-verified and expression of the transcription factor confirmed by immunohistochemistry with antibodies to the myc tag or with factor-specific antibodies.

### Epifluorescence and Confocal Microscopy

Micrographs of tissue sections were taken either with epifluorescence microscopes (Leica DM6, DM6000) or confocal microscopes (Leica SP8 or Zeiss LSM710). Whole embryo images were taken with a fluorescence dissecting stereomicroscope (Leica MZ16FA). All RNA Scope images, for quantification and for analysis, were taken using confocal microscopes on a 40x objective in order to resolve single puncta. Human sections were imaged with a laser scanning confocal microscope (FV1000, Olympus) and processed using ImageJ (NIH) and Adobe Photoshop.

## QUANTIFICATION AND STATISTICAL ANALYSIS

### Bioinformatics

scRNA-seq data used in this study were previously published (Delile et al., 2019), using spinal cord cells from e9.5, e10.5, e11.5, e12.5 and e13.5 mouse embryos and processed via the 10x Genomics Chromium Single Cell 3′ v2 protocol. Raw data were extracted from ArrayExpress E-MTAB-7320). CellRanger v 2.1.1 (Zheng et al., 2017) was used to align reads to the 10X mm10 mouse reference genome v2.1.0, filter barcodes and quantify genes. Biological replicates from the same time points were then merged using CellRanger’s *aggregate* function. All downstream analyses were performed on these 41 009 cells, using the Seurat v.3.1.1. R package.

Using Delile *et al*.’s (2019) annotation, only cells classified as neurons were kept for further analysis (18 048 cells). From these, data for each timepoint was normalized and highly variable genes identified using SCTansform’s normalization and variance stabilizing methods (Hafemeister and Satija, 2019). Different timepoints were then integrated using CCA (Stuart et al., 2019). From the integrated dataset, 2 subsets were created: 1) all cells expressing *Lmx1b* (*Lmx1b*+) were isolated (2614 cells) and 2) all early (e9.5, e10.5 and e11.5) *Lmx1b*+ cells (186 cells). For each of these subsets, dimensionality reduction (PCA and UMAP) was applied and clusters identified using Seurat’s SNN modularity optimisation-based clustering Louvain algorithm. Differential expression analysis was performed on the non-integrated assay to identify markers for each cluster (Wilcoxon Rank Sum test).

A differential expression analysis was then performed on clusters of interest in each of these 3 subsets (cluster 6 from *Lmx1b*+ cells and clusters 2 and 3 in early *Lmx1b*+ cells). Each of these clusters was compared to all other neuron cells in order to identify specific markers within these clusters. This analysis was limited to genes which had on average, at least 0.1 log-Fold difference between the two groups compared and present in at least 10% of cells of either group. Markers were then considered significantly differentially expressed if adjusted p-values < 0.05.

### Cell counts

All cell counts were done with Image J v.2.0.0 software, using the cell counter plugin. Data was recorded and sorted in Microsoft Excel.

### Animal Behaviour

Video-recordings of mice were quantified by R.B.R and M.B using Aegisub (free subtitling software), which includes video annotation functions allowing precise start and stop times for specified behaviours to be recorded. Data was exported and then sorted in Microsoft Excel.

### Data representation

Graphs display data points from individual animals (hollow circles), mean data from all animals (bars), and ± standard error of the mean (error bars).

### Numbers

All numbers are noted in figure legends. In all experiments using adult mice, n represents unique individuals. In all experiments using embryonic mice, n represents unique embryos. No data represented as single ns have been pooled from multiple individual animals.

### Statistics

All statistical analyses were performed using GraphPad Prism v8.3.0 software, except those involving single cell RNA-Seq data processing, which are described in Bioinformatics (above). Statistical tests used are described in figure legends. Significance is represented as ns: non significant, *: p<0.05, **: p<0.01 or ***: p<0.001.

## DATA AND CODE AVAILABILITY

The published article includes all datasets/code generated during this study. Single cell RNA-Seq data analyzed here was generated by Delile et al. (2019) and was obtained per their instructions from Array Express (https://www.ebi.ac.uk/arrayexpress/) with accession number “E-MTAB-7320”.

## KEY RESOURCES TABLE

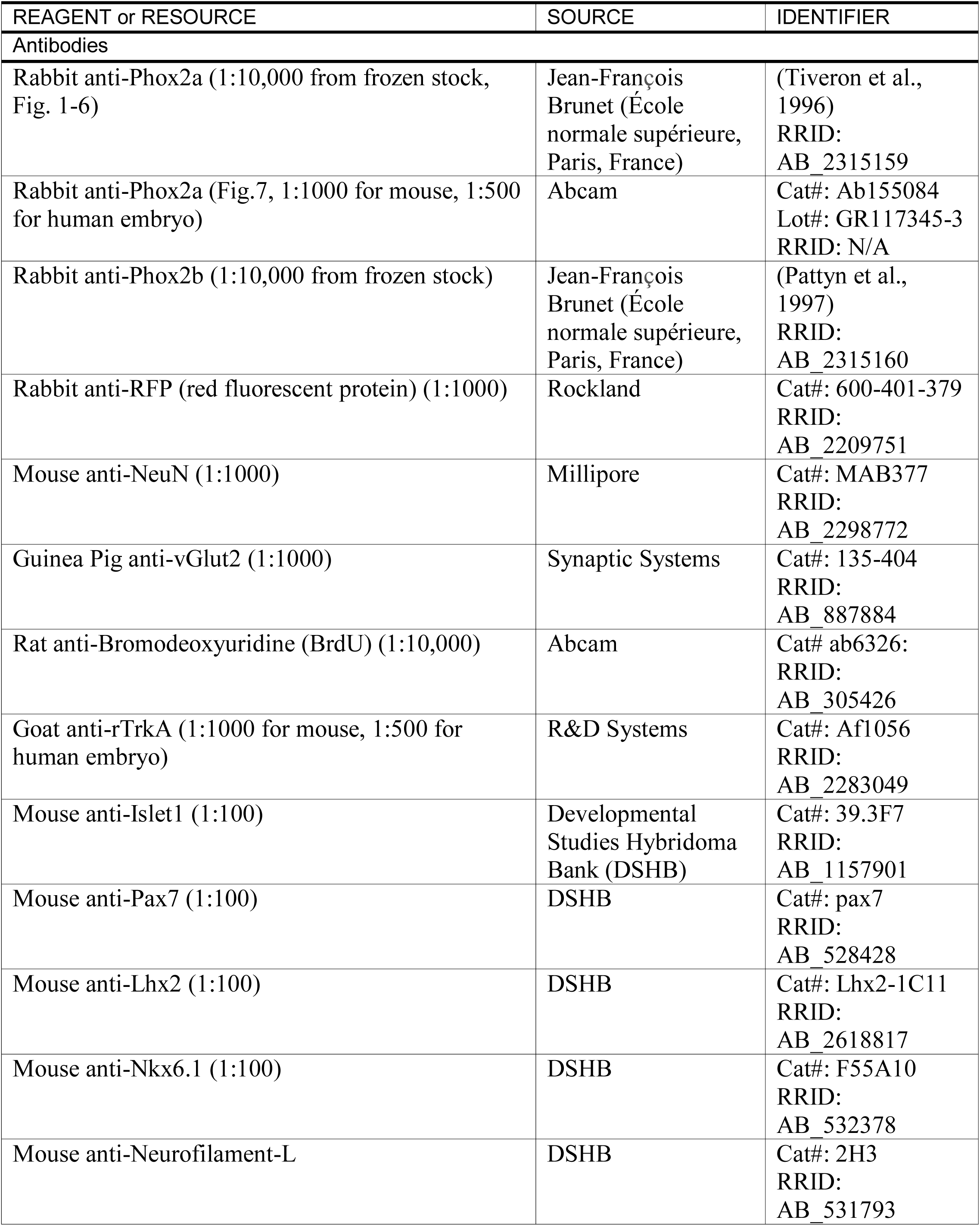
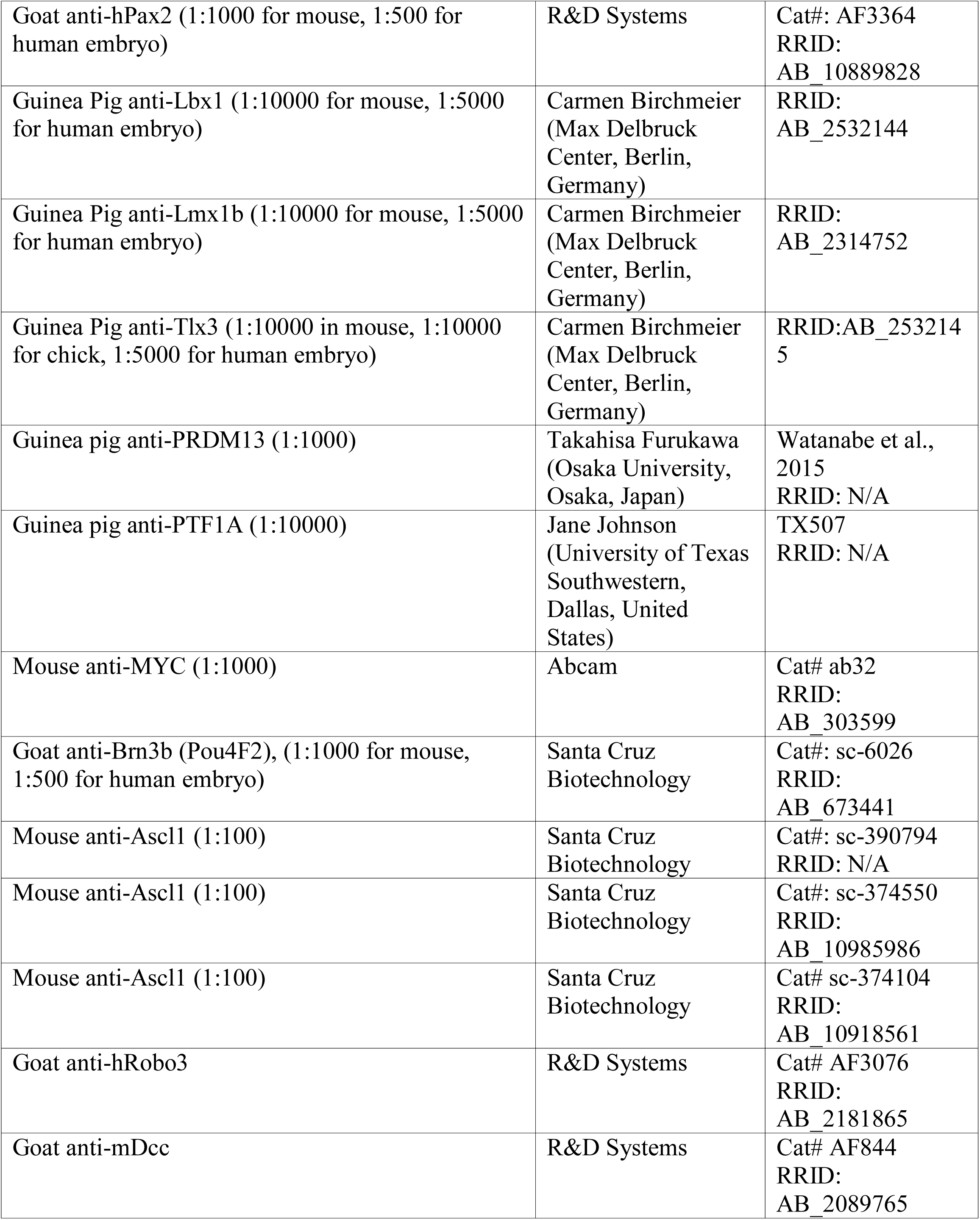
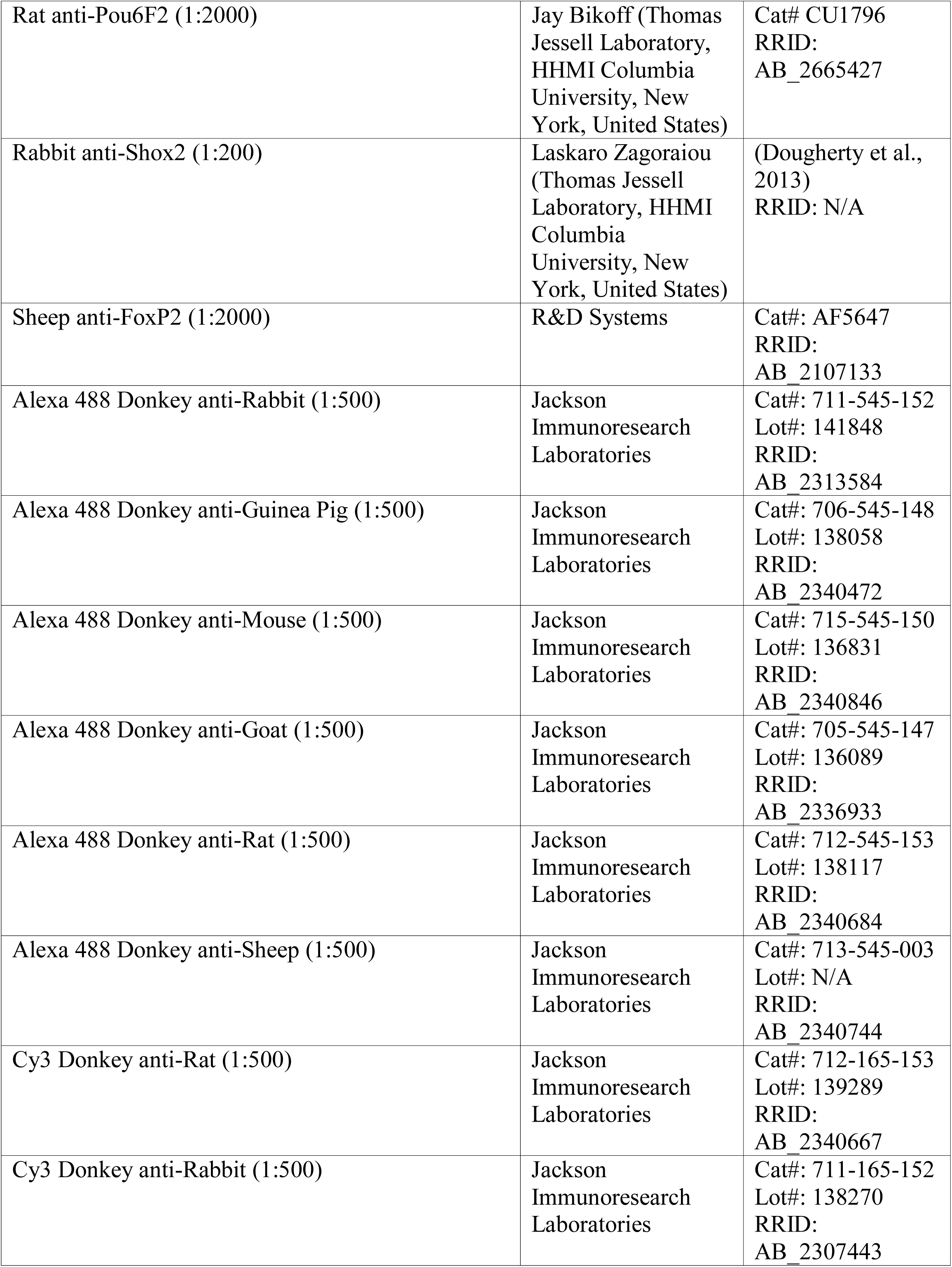
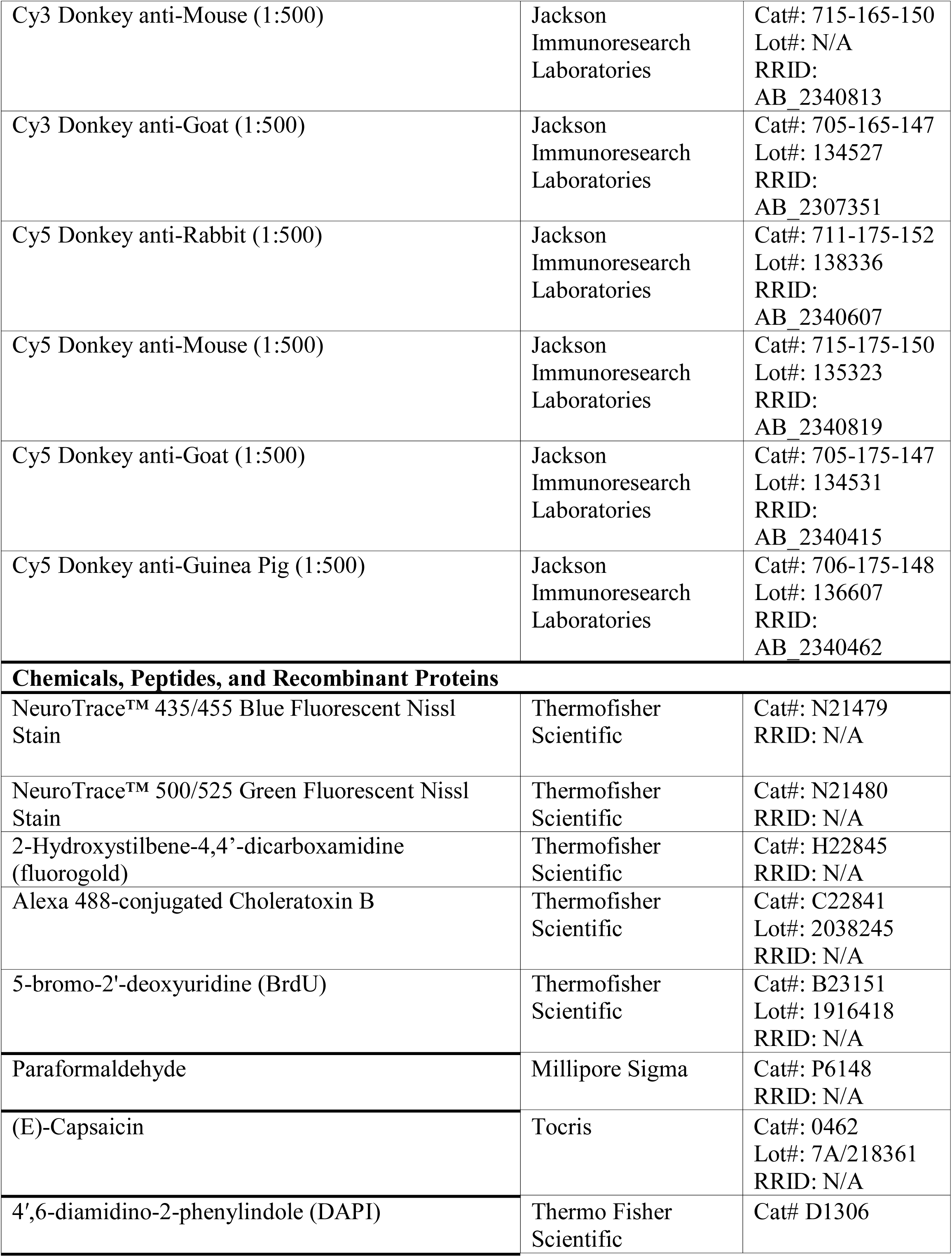
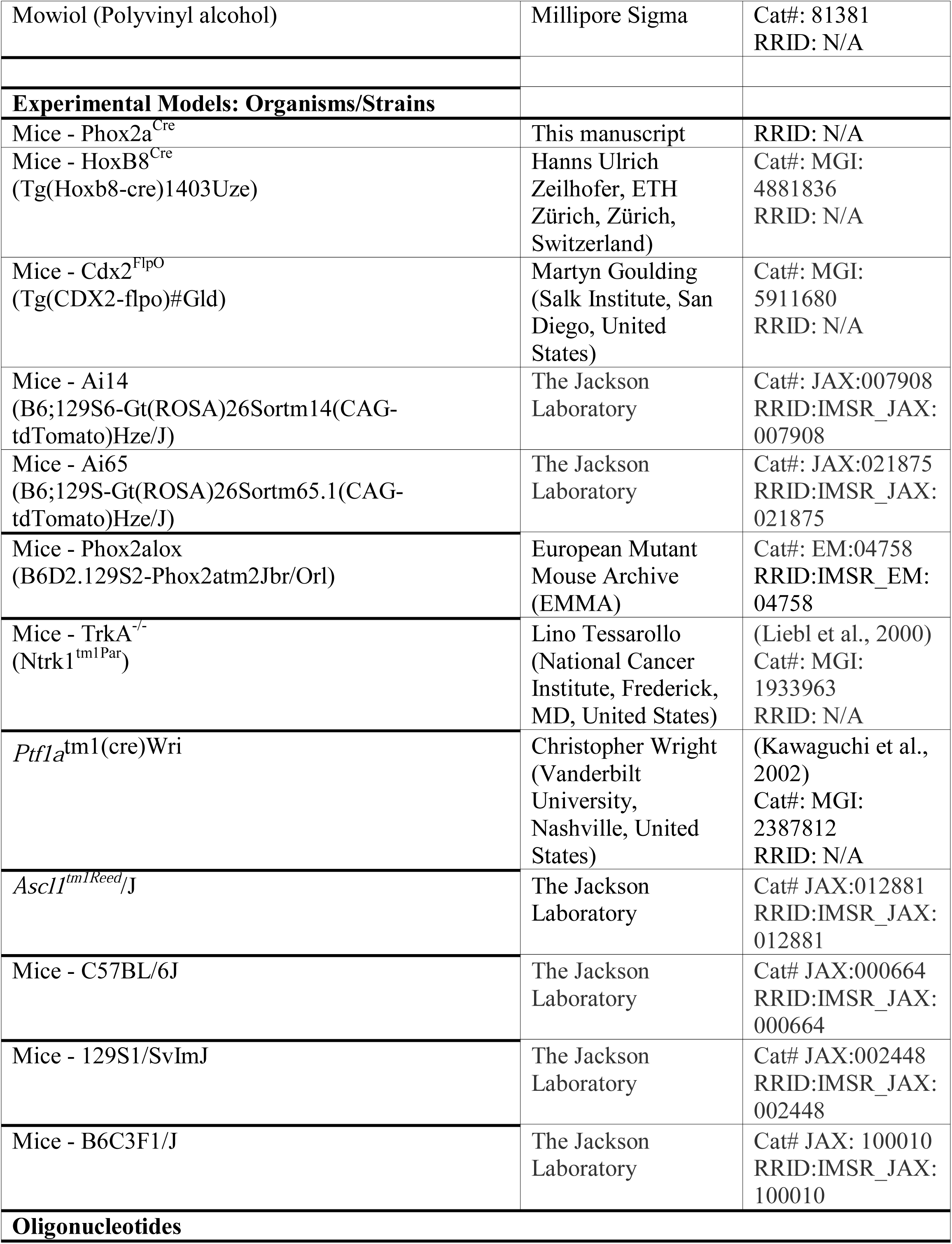
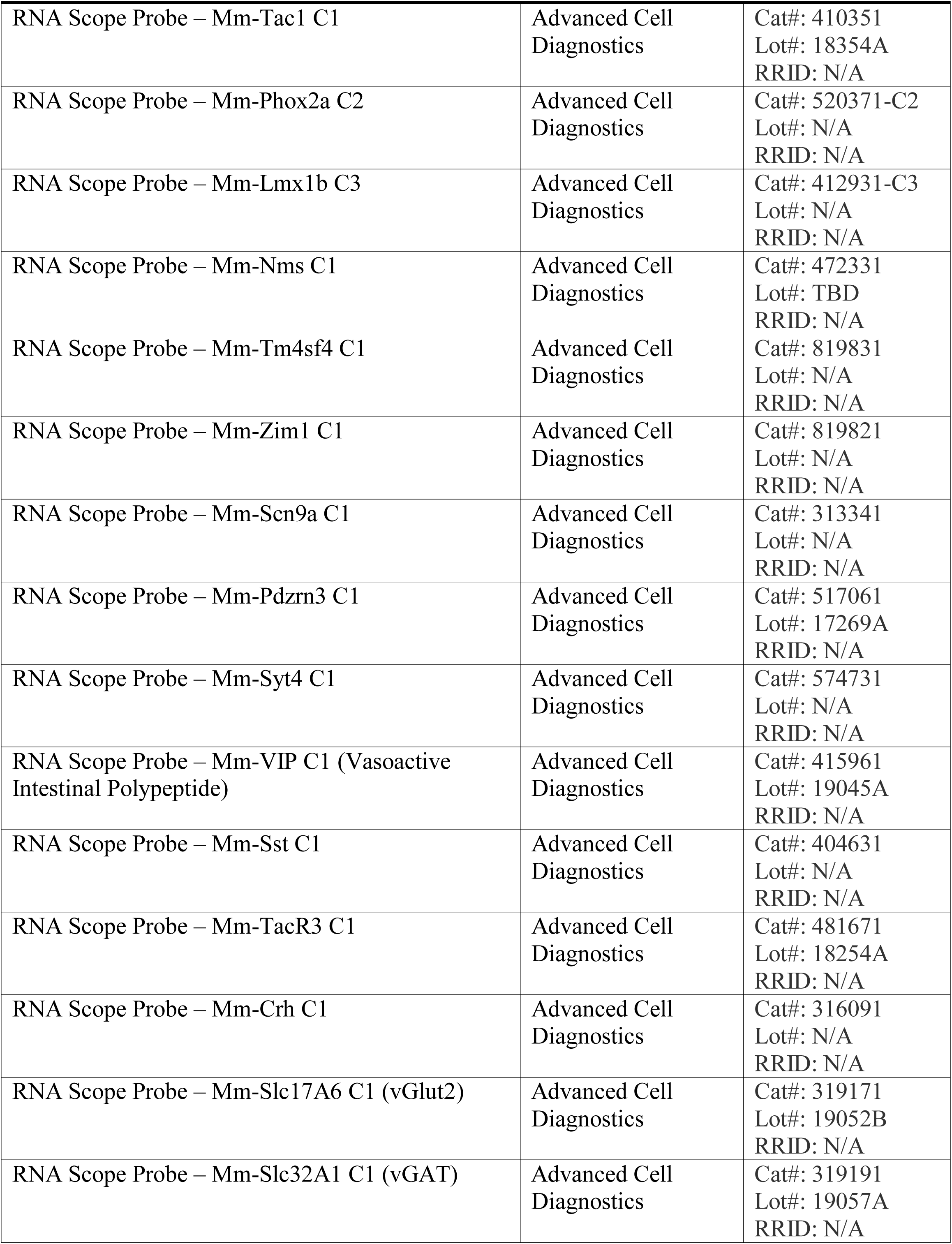
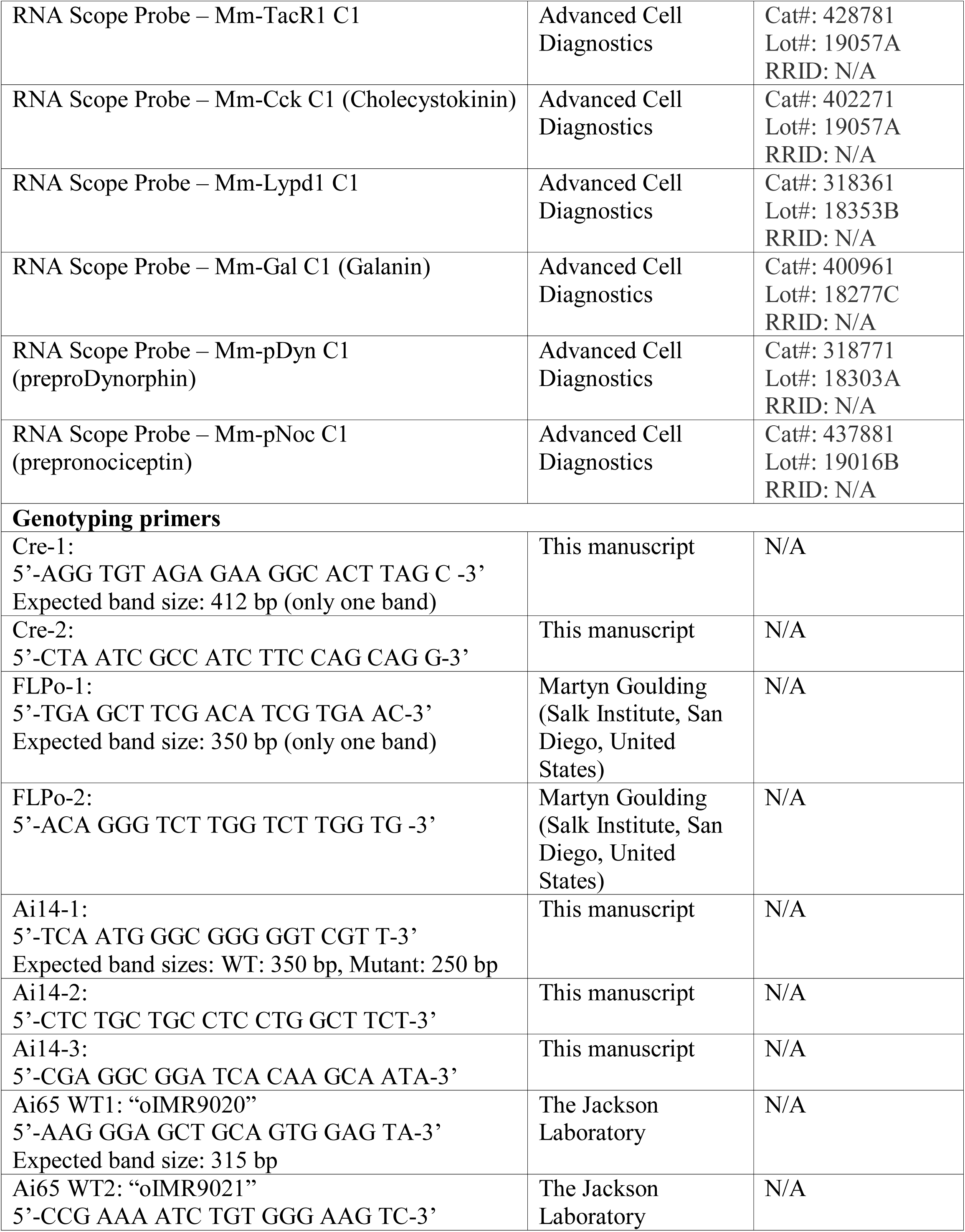
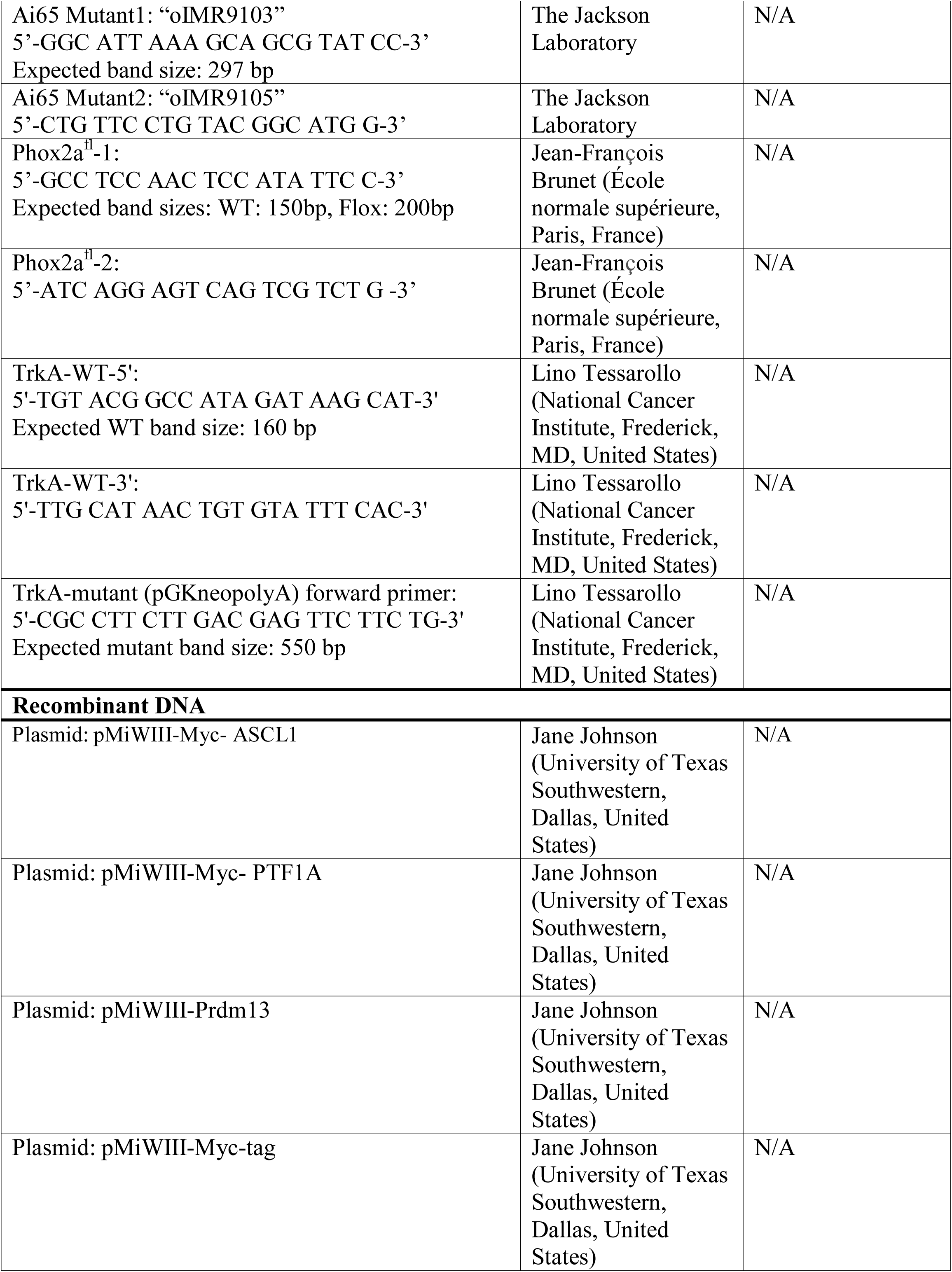
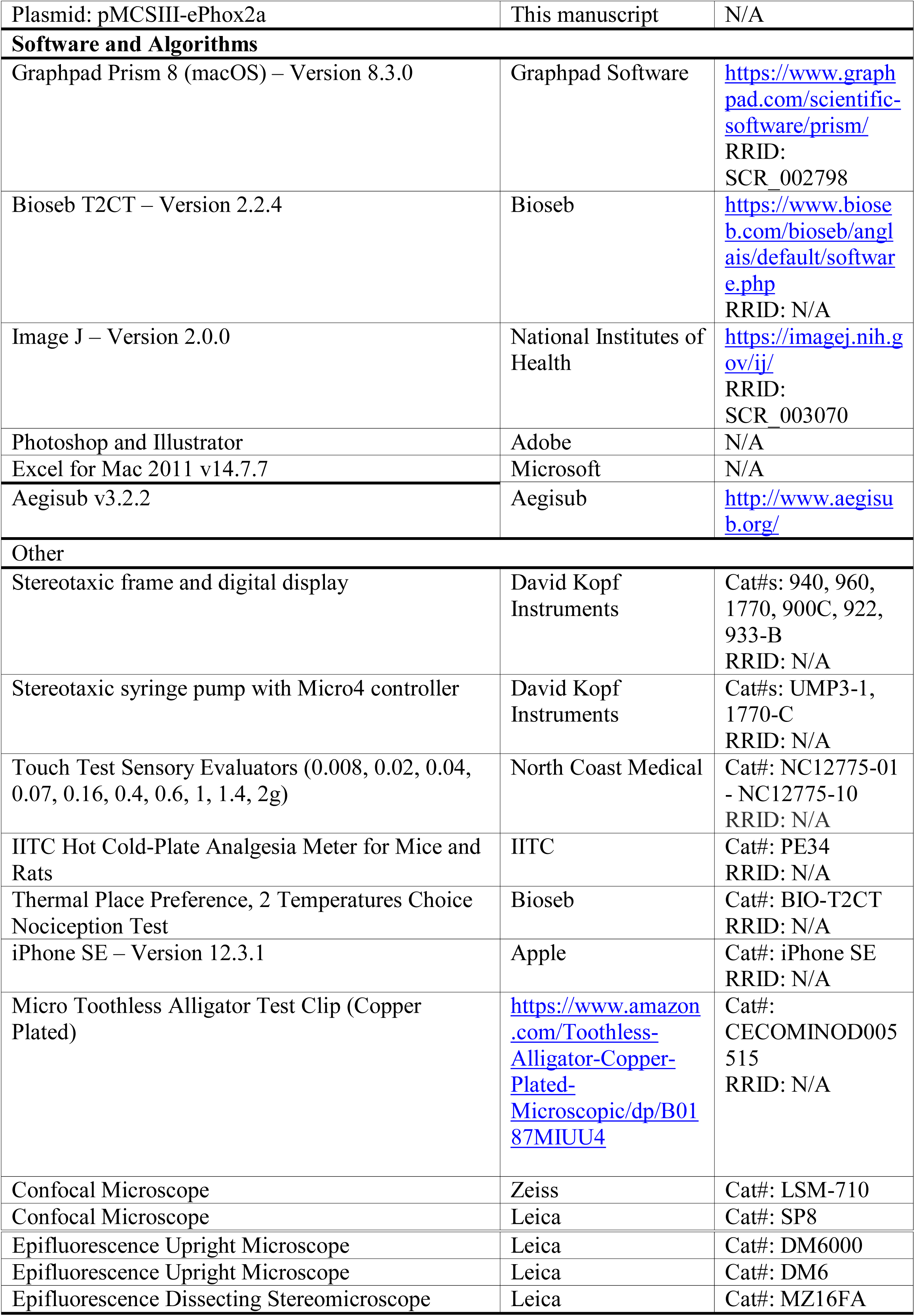

## References

Allen Institute for Brain Science (2004). © 2004 Allen Institute for Brain Science. Allen Mouse Brain Atlas. Available from: https://mouse.brain-map.org/static/atlas.

Allen Institute for Brain Science (2008). © 2008 Allen Institute for Brain Science. Allen Spinal Cord Atlas. Available from: [http://mousespinal.brain-map.org/].

Altman, J., and Bayer, S.A. (2001). Development of the Human Spinal Cord: An Interpretation Based on Experimental Studies in Animals (Oxford University Press).

Anderson, K.R., Singer, R.A., Balderes, D.A., Hernandez-Lagunas, L., Johnson, C.W., Artinger, K.B., and Sussel, L. (2011). The L6 domain tetraspanin Tm4sf4 regulates endocrine pancreas differentiation and directed cell migration. Development 138, 3213–3224.

Andrew, D., and Craig, A.D. (2001). Spinothalamic lamina I neurons selectively sensitive to histamine: a central neural pathway for itch. Nat Neurosci 4, 72–77.

Apkarian, A.V., Stevens, R.T., and Hodge, C.J. (1985). Funicular location of ascending axons of lamina I cells in the cat spinal cord. Brain Res 334, 160–164.

Arber, S. (2012). Motor circuits in action: specification, connectivity, and function. Neuron 74, 975–989.

Bernard, J.F., Dallel, R., Raboisson, P., Villanueva, L., and Le Bars, D. (1995). Organization of the efferent projections from the spinal cervical enlargement to the parabrachial area and periaqueductal gray: a PHA-L study in the rat. J Comp Neurol 353, 480–505.

Berthier, M., Starkstein, S., and Leiguarda, R. (1988). Asymbolia for pain: a sensory-limbic disconnection syndrome. Ann Neurol 24, 41–49.

Borromeo, M.D., Meredith, D.M., Castro, D.S., Chang, J.C., Tung, K.C., Guillemot, F., and Johnson, J.E. (2014). A transcription factor network specifying inhibitory versus excitatory neurons in the dorsal spinal cord. Development 141, 2803–2812.

Bouet, V., Boulouard, M., Toutain, J., Divoux, D., Bernaudin, M., Schumann-Bard, P., and Freret, T. (2009). The adhesive removal test: a sensitive method to assess sensorimotor deficits in mice. Nat Protoc 4, 1560–1564.

Bourgeais, L., Monconduit, L., Villanueva, L., and Bernard, J.F. (2001). Parabrachial internal lateral neurons convey nociceptive messages from the deep laminas of the dorsal horn to the intralaminar thalamus. J Neurosci 21, 2159–2165.

Britz, O., Zhang, J., Grossmann, K.S., Dyck, J., Kim, J.C., Dymecki, S., Gosgnach, S., and Goulding, M. (2015). A genetically defined asymmetry underlies the inhibitory control of flexor-extensor locomotor movements. Elife 4.

Brunet, J.F., and Pattyn, A. (2002). Phox2 genes - from patterning to connectivity. Curr Opin Genet Dev 12, 435–440.

Cameron, D., Polgar, E., Gutierrez-Mecinas, M., Gomez-Lima, M., Watanabe, M., and Todd, A.J. (2015). The organisation of spinoparabrachial neurons in the mouse. Pain 156, 2061–2071.

Chang, J.C., Meredith, D.M., Mayer, P.R., Borromeo, M.D., Lai, H.C., Ou, Y.H., and Johnson, J.E. (2013). Prdm13 mediates the balance of inhibitory and excitatory neurons in somatosensory circuits. Dev Cell 25, 182–195.

Chaplan, S.R., Bach, F.W., Pogrel, J.W., Chung, J.M., and Yaksh, T.L. (1994). Quantitative assessment of tactile allodynia in the rat paw. J Neurosci Methods 53, 55–63.

Colburn, R.W., Lubin, M.L., Stone, D.J., Jr., Wang, Y., Lawrence, D., D’Andrea, M.R., Brandt, M.R., Liu, Y., Flores, C.M., and Qin, N. (2007). Attenuated cold sensitivity in TRPM8 null mice. Neuron 54, 379–386.

Cox, J.J., Reimann, F., Nicholas, A.K., Thornton, G., Roberts, E., Springell, K., Karbani, G., Jafri, H., Mannan, J., Raashid, Y., et al. (2006). An SCN9A channelopathy causes congenital inability to experience pain. Nature 444, 894–898.

Craig, A.D. (1996). An ascending general homeostatic afferent pathway originating in lamina I. Prog Brain Res 107, 225–242.

Craig, A.D. (2003a). A new view of pain as a homeostatic emotion. Trends Neurosci 26, 303–307.

Craig, A.D. (2003b). Pain mechanisms: labeled lines versus convergence in central processing. Annu Rev Neurosci 26, 1–30.

Craig, A.D. (2004). Lamina I, but not lamina V, spinothalamic neurons exhibit responses that correspond with burning pain. J Neurophysiol 92, 2604–2609.

Craig, A.D., and Serrano, L.P. (1994). Effects of systemic morphine on lamina I spinothalamic tract neurons in the cat. Brain Res 636, 233–244.

da Silva, R.V., Johannssen, H.C., Wyss, M.T., Roome, R.B., Bourojeni, F.B., Stifani, N., Marsh, A.P.L., Ryan, M.M., Lockhart, P.J., Leventer, R.J., et al. (2018). DCC Is Required for the Development of Nociceptive Topognosis in Mice and Humans. Cell Rep 22, 1105–1114.

Davidson, S., Truong, H., and Giesler, G.J., Jr. (2010). Quantitative analysis of spinothalamic tract neurons in adult and developing mouse. The Journal of comparative neurology 518, 3193–3204.

Delile, J., Rayon, T., Melchionda, M., Edwards, A., Briscoe, J., and Sagner, A. (2019). Single cell transcriptomics reveals spatial and temporal dynamics of gene expression in the developing mouse spinal cord. Development 146.

Ding, Y.Q., Kim, J.Y., Xu, Y.S., Rao, Y., and Chen, Z.F. (2005). Ventral migration of early-born neurons requires Dcc and is essential for the projections of primary afferents in the spinal cord. Development 132, 2047–2056.

Ding, Y.Q., Yin, J., Kania, A., Zhao, Z.Q., Johnson, R.L., and Chen, Z.F. (2004). Lmx1b controls the differentiation and migration of the superficial dorsal horn neurons of the spinal cord. Development 131, 3693–3703.

Dong, H.W. (2008). The Allen reference atlas: A digital color brain atlas of the C57Bl/6J male mouse (John Wiley & Sons Inc).

Duan, B., Cheng, L., Bourane, S., Britz, O., Padilla, C., Garcia-Campmany, L., Krashes, M., Knowlton, W., Velasquez, T., Ren, X., et al. (2014). Identification of spinal circuits transmitting and gating mechanical pain. Cell 159, 1417–1432.

Fan, Y., Chen, P., Raza, M.U., Szebeni, A., Szebeni, K., Ordway, G.A., Stockmeier, C.A., and Zhu, M.Y. (2018). Altered Expression of Phox2 Transcription Factors in the Locus Coeruleus in Major Depressive Disorder Mimicked by Chronic Stress and Corticosterone Treatment In Vivo and In Vitro. Neuroscience 393, 123–137.

Fazeli, A., Dickinson, S.L., Hermiston, M.L., Tighe, R.V., Steen, R.G., Small, C.G., Stoeckli, E.T., Keino-Masu, K., Masu, M., Rayburn, H., et al. (1997). Phenotype of mice lacking functional Deleted in colorectal cancer (Dcc) gene. Nature 386, 796–804.

Feil, K., and Herbert, H. (1995). Topographic organization of spinal and trigeminal somatosensory pathways to the rat parabrachial and Kolliker-Fuse nuclei. J Comp Neurol 353, 506–528.

Fernandes, E.C., Santos, I.C., Kokai, E., Luz, L.L., Szucs, P., and Safronov, B.V. (2018). Low- and high-threshold primary afferent inputs to spinal lamina III antenna-type neurons. Pain 159, 2214–2222.

Freeman, W., and Watts, J.W. (1948). Pain mechanisms and the frontal lobes; a study of prefrontal lobotomy for intractable pain. Ann Intern Med 28, 747–754.

Gauriau, C., and Bernard, J.F. (2004). A comparative reappraisal of projections from the superficial laminae of the dorsal horn in the rat: the forebrain. J Comp Neurol 468, 24–56.

GENSAT (2008). The Gene Expression Nervous System Atlas (GENSAT) Project, NINDS Contracts N01NS02331 & HHSN271200723701C to The Rockefeller University (New York, NY). Available from: http://www.gensat.org/GeneProgressTracker.jsp?gensatGeneID=1715.

Glasgow, S.M., Henke, R.M., Macdonald, R.J., Wright, C.V., and Johnson, J.E. (2005). Ptf1a determines GABAergic over glutamatergic neuronal cell fate in the spinal cord dorsal horn. Development 132, 5461–5469.

Goulding, M. (2009). Circuits controlling vertebrate locomotion: moving in a new direction. Nat Rev Neurosci 10, 507–518.

Gowan, K., Helms, A.W., Hunsaker, T.L., Collisson, T., Ebert, P.J., Odom, R., and Johnson, J.E. (2001). Crossinhibitory activities of Ngn1 and Math1 allow specification of distinct dorsal interneurons. Neuron 31, 219–232.

Guilbaud, G., Peschanski, M., Gautron, M., and Binder, D. (1980). Neurones responding to noxious stimulation in VB complex and caudal adjacent regions in the thalamus of the rat. Pain 8, 303–318.

Hafemeister, C., and Satija, R. (2019). Normalization and variance stabilization of single-cell RNA-seq data using regularized negative binomial regression. Genome Biol 20, 296.

Hamburger, V., and Hamilton, H.L. (1951). A series of normal stages in the development of the chick embryo. J Morphol 88, 49–92.

Han, S., Soleiman, M.T., Soden, M.E., Zweifel, L.S., and Palmiter, R.D. (2015). Elucidating an Affective Pain Circuit that Creates a Threat Memory. Cell 162, 363–374.

Hargreaves, K., Dubner, R., Brown, F., Flores, C., and Joris, J. (1988). A new and sensitive method for measuring thermal nociception in cutaneous hyperalgesia. Pain 32, 77–88.

Helms, A.W., Battiste, J., Henke, R.M., Nakada, Y., Simplicio, N., Guillemot, F., and Johnson, J.E. (2005). Sequential roles for Mash1 and Ngn2 in the generation of dorsal spinal cord interneurons. Development 132, 2709–2719.

Hori, K., Cholewa-Waclaw, J., Nakada, Y., Glasgow, S.M., Masui, T., Henke, R.M., Wildner, H., Martarelli, B., Beres, T.M., Epstein, J.A., et al. (2008). A nonclassical bHLH Rbpj transcription factor complex is required for specification of GABAergic neurons independent of Notch signaling. Genes Dev 22, 166–178.

Hua, Z.L., Jeon, S., Caterina, M.J., and Nathans, J. (2014). Frizzled3 is required for the development of multiple axon tracts in the mouse central nervous system. Proc Natl Acad Sci U S A 111, E3005–3014.

Huang, T., Lin, S.H., Malewicz, N.M., Zhang, Y., Zhang, Y., Goulding, M., LaMotte, R.H., and Ma, Q. (2019). Identifying the pathways required for coping behaviours associated with sustained pain. Nature 565, 86–90.

Hyndman, O.R., and Jarvis, F.J. (1940). Gastric crisis of tabes dorsalis: Treatment by anterior chordotomy in eight cases. Archives of Surgery 40, 997–1013.

Hyndman, O.R., and Wolkin, J. (1943). Anterior chordotomy: Further observations on physiologic results and optimum manner of performance. Archives of Neurology & Psychiatry 50, 129–148.

Kawaguchi, Y., Cooper, B., Gannon, M., Ray, M., MacDonald, R.J., and Wright, C.V. (2002). The role of the transcriptional regulator Ptf1a in converting intestinal to pancreatic progenitors. Nat Genet 32, 128–134.

Keay, K.A., and Bandler, R. (2002). Distinct central representations of inescapable and escapable pain: observations and speculation. Exp Physiol 87, 275–279.

Kim, E.J., Battiste, J., Nakagawa, Y., and Johnson, J.E. (2008). Ascl1 (Mash1) lineage cells contribute to discrete cell populations in CNS architecture. Mol Cell Neurosci 38, 595–606.

Kitamura, T., Yamada, J., Sato, H., and Yamashita, K. (1993). Cells of origin of the spinoparabrachial fibers in the rat: a study with fast blue and WGA-HRP. J Comp Neurol 328, 449–461.

Kohwi, M., and Doe, C.Q. (2013). Temporal fate specification and neural progenitor competence during development. Nat Rev Neurosci 14, 823–838.

Lai, H.C., Seal, R.P., and Johnson, J.E. (2016). Making sense out of spinal cord somatosensory development. Development 143, 3434–3448.

Leah, J., Menetrey, D., and de Pommery, J. (1988). neuropeptides in long ascending spinal tract cells in the rat: evidence for parallel processing of ascending information. Neuroscience 24, 195–207.

Leung, C.T., Coulombe, P.A., and Reed, R.R. (2007). Contribution of olfactory neural stem cells to tissue maintenance and regeneration. Nat Neurosci 10, 720–726.

Mantyh, P.W., Rogers, S.D., Honore, P., Allen, B.J., Ghilardi, J.R., Li, J., Daughters, R.S., Lappi, D.A., Wiley, R.G., and Simone, D.A. (1997). Inhibition of hyperalgesia by ablation of lamina I spinal neurons expressing the substance P receptor. Science 278, 275–279.

Marshall, G.E., Shehab, S.A., Spike, R.C., and Todd, A.J. (1996). Neurokinin-1 receptors on lumbar spinothalamic neurons in the rat. Neuroscience 72, 255–263.

Masullo, L., Mariotti, L., Alexandre, N., Freire-Pritchett, P., Boulanger, J., and Tripodi, M. (2019). Genetically Defined Functional Modules for Spatial Orienting in the Mouse Superior Colliculus. Curr Biol 29, 2892-2904.e2898.

Matsunaga, E., Araki, I., and Nakamura, H. (2001). Role of Pax3/7 in the tectum regionalization. Development 128, 4069–4077.

McMahon, S.B., and Wall, P.D. (1983). A system of rat spinal cord lamina 1 cells projecting through the contralateral dorsolateral funiculus. J Comp Neurol 214, 217–223.

Melzack, R., and Casey, K.L. (1968). Sensory, Motivational, and Central Control Determinants of Pain: A new conceptual model. In The Skin Senses, D.R. Kenshalo, ed. (Charles C. Thomas), pp. 423–439.

Meredith, D.M., Borromeo, M.D., Deering, T.G., Casey, B.H., Savage, T.K., Mayer, P.R., Hoang, C., Tung, K.C., Kumar, M., Shen, C., et al. (2013). Program specificity for Ptf1a in pancreas versus neural tube development correlates with distinct collaborating cofactors and chromatin accessibility. Mol Cell Biol 33, 3166–3179.

Minett, M.S., Nassar, M.A., Clark, A.K., Passmore, G., Dickenson, A.H., Wang, F., Malcangio, M., and Wood, J.N. (2012). Distinct Nav1.7-dependent pain sensations require different sets of sensory and sympathetic neurons. Nat Commun 3, 791.

Mogil, J.S., Wilson, S.G., Bon, K., Lee, S.E., Chung, K., Raber, P., Pieper, J.O., Hain, H.S., Belknap, J.K., Hubert, L., et al. (1999). Heritability of nociception I: responses of 11 inbred mouse strains on 12 measures of nociception. Pain 80, 67–82.

Mona, B., Uruena, A., Kollipara, R.K., Ma, Z., Borromeo, M.D., Chang, J.C., and Johnson, J.E. (2017). Repression by PRDM13 is critical for generating precision in neuronal identity. Elife 6.

Morin, X., Cremer, H., Hirsch, M.R., Kapur, R.P., Goridis, C., and Brunet, J.F. (1997). Defects in sensory and autonomic ganglia and absence of locus coeruleus in mice deficient for the homeobox gene Phox2a. Neuron 18, 411–423.

Nakano, M., Yamada, K., Fain, J., Sener, E.C., Selleck, C.J., Awad, A.H., Zwaan, J., Mullaney, P.B., Bosley, T.M., and Engle, E.C. (2001). Homozygous mutations in ARIX(PHOX2A) result in congenital fibrosis of the extraocular muscles type 2. Nat Genet 29, 315–320.

Nishida, K., and Ito, S. (2017). Developmental origin of long-range neurons in the superficial dorsal spinal cord. Eur J Neurosci 46, 2608–2619.

Nornes, H.O., and Carry, M. (1978). Neurogenesis in spinal cord of mouse: an autoradiographic analysis. Brain Res 159, 1–6.

Pattyn, A., Morin, X., Cremer, H., Goridis, C., and Brunet, J.F. (1997). Expression and interactions of the two closely related homeobox genes Phox2a and Phox2b during neurogenesis. Development 124, 4065–4075.

Petitjean, H., Bourojeni, F.B., Tsao, D., Davidova, A., Sotocinal, S.G., Mogil, J.S., Kania, A., and Sharif-Naeini, R. (2019). Recruitment of Spinoparabrachial Neurons by Dorsal Horn Calretinin Neurons. Cell Rep 28, 1429-1438.e1424.

Price, D.D., and Dubner, R. (1977). Neurons that subserve the sensory-discriminative aspects of pain. Pain 3, 307–338.

Ren, K., and Dubner, R. (2009). Descending Control Mechanisms. In Science of Pain, A.I. Basbaum, and C. Bushnell, eds. (Elsevier), pp. 723–749.

Rubins, J.L., and Friedman, E.D. (1948). Asymbolia for pain. Archives of Neurology & Psychiatry 60, 554–573.

Sabatier, C., Plump, A.S., Le, M., Brose, K., Tamada, A., Murakami, F., Lee, E.Y., and Tessier-Lavigne, M. (2004). The divergent Robo family protein rig-1/Robo3 is a negative regulator of slit responsiveness required for midline crossing by commissural axons. Cell 117, 157–169.

Sakurada, T., Katsumata, K., Tan-No, K., Sakurada, S., and Kisara, K. (1992). The capsaicin test in mice for evaluating tachykinin antagonists in the spinal cord. Neuropharmacology 31, 1279–1285.

Schoenen, J. (1982). The dendritic organization of the human spinal cord: the dorsal horn. Neuroscience 7, 2057–2087.

Smeyne, R.J., Klein, R., Schnapp, A., Long, L.K., Bryant, S., Lewin, A., Lira, S.A., and Barbacid, M. (1994). Severe sensory and sympathetic neuropathies in mice carrying a disrupted Trk/NGF receptor gene. Nature 368, 246–249.

Spiller, W.G., and Martin, E. (1912). The treatment of persistent pain of organic origin in the lower part of the body by division of the anterolateral column of the spinal cord. Journal of the American Medical Association LVIII, 1489–1490.

Stuart, T., Butler, A., Hoffman, P., Hafemeister, C., Papalexi, E., Mauck, W.M., 3rd, Hao, Y., Stoeckius, M., Smibert, P., and Satija, R. (2019). Comprehensive Integration of Single-Cell Data. Cell 177, 1888-1902.e1821.

Szabo, N.E., da Silva, R.V., Sotocinal, S.G., Zeilhofer, H.U., Mogil, J.S., and Kania, A. (2015). Hoxb8 intersection defines a role for Lmx1b in excitatory dorsal horn neuron development, spinofugal connectivity, and nociception. J Neurosci 35, 5233–5246.

Wang, X., Babayan, A.H., Basbaum, A.I., and Phelps, P.E. (2012). Loss of the Reelin-signaling pathway differentially disrupts heat, mechanical and chemical nociceptive processing. Neuroscience 226, 441–450.

Warming, S., Costantino, N., Court, D.L., Jenkins, N.A., and Copeland, N.G. (2005). Simple and highly efficient BAC recombineering using galK selection. Nucleic Acids Res 33, e36.

Watanabe, K., Tamamaki, N., Furuta, T., Ackerman, S.L., Ikenaka, K., and Ono, K. (2006). Dorsally derived netrin 1 provides an inhibitory cue and elaborates the ‘waiting period’ for primary sensory axons in the developing spinal cord. Development 133, 1379–1387.

Willis, W.D., Kenshalo, D.R., Jr., and Leonard, R.B. (1979). The cells of origin of the primate spinothalamic tract. J Comp Neurol 188, 543–573.

Willis, W.D., Trevino, D.L., Coulter, J.D., and Maunz, R.A. (1974). Responses of primate spinothalamic tract neurons to natural stimulation of hindlimb. J Neurophysiol 37, 358–372.

Witschi, R., Johansson, T., Morscher, G., Scheurer, L., Deschamps, J., and Zeilhofer, H.U. (2010). Hoxb8-Cre mice: A tool for brain-sparing conditional gene deletion. Genesis 48, 596–602.

Yvone, G.M., Zhao-Fleming, H.H., Udeochu, J.C., Chavez-Martinez, C.L., Wang, A., Hirose-Ikeda, M., and Phelps, P.E. (2017). Disabled-1 dorsal horn spinal cord neurons co-express Lmx1b and function in nociceptive circuits. Eur J Neurosci 45, 733–747.

Zheng, G.X., Terry, J.M., Belgrader, P., Ryvkin, P., Bent, Z.W., Wilson, R., Ziraldo, S.B., Wheeler, T.D., McDermott, G.P., Zhu, J., et al. (2017). Massively parallel digital transcriptional profiling of single cells. Nat Commun 8, 14049.

