## Supplemental Information for "Phox2a defines a developmental origin of the anterolateral system in mice and humans"

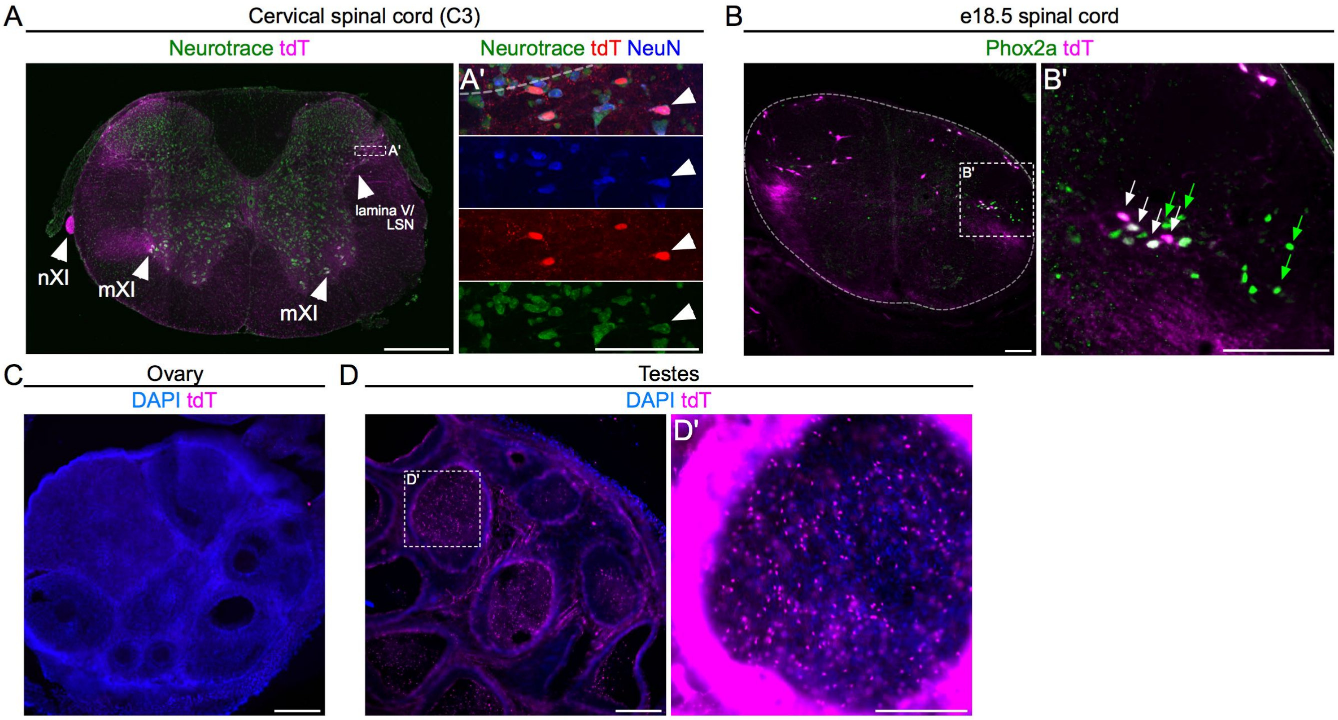


Roome et al., Figure S1

**Figure S1: Spinal Phox2a$^{Cre}$ neurons reside in lamina I, V and LSN. Related to Figure 1.**

(A) TdT+ neurons in the cervical spinal cord of adult *Phox2a*$^{Cre}$; *R26*$^{LSL\text{-}tdT/+}$ mice. (A') Magnified box from (A) showing lamina V/LSN Neurotrace, tdT and NeuN co-labelling. (B) Expression of tdT in Phox2a+ neurons in an e18.5 *Phox2a*$^{Cre}$; *R26*$^{LSL\text{-}tdT/+}$ mouse embryo. (C) Adult *Phox2a*$^{Cre}$; *R26*$^{LSL\text{-}tdT/+}$ mouse ovaries do not express tdT. (D) Adult *Phox2a*$^{Cre}$; *R26*$^{LSL\text{-}tdT/+}$ mouse testes express tdT. A minority of cells in the lumen of seminiferous tubules express tdT; occasional germline recombination was detected in progeny of *Phox2a*$^{Cre}$; *R26*$^{LSL\text{-}tdT/+}$ males.

Numbers: (A) n=3 adult *Phox2a*$^{Cre}$; *R26*$^{LSL\text{-}tdT/+}$ mice, (B) n=3 e18.5 *Phox2a*$^{Cre}$; *R26*$^{LSL\text{-}tdT/+}$ embryos.

Scale bars: (A) 500 µm, (A', B, D') 100 µm, (C, D) 200 µm.

Abbreviations: mXI (accessory motor nucleus), nXI (accessory motor nerve).

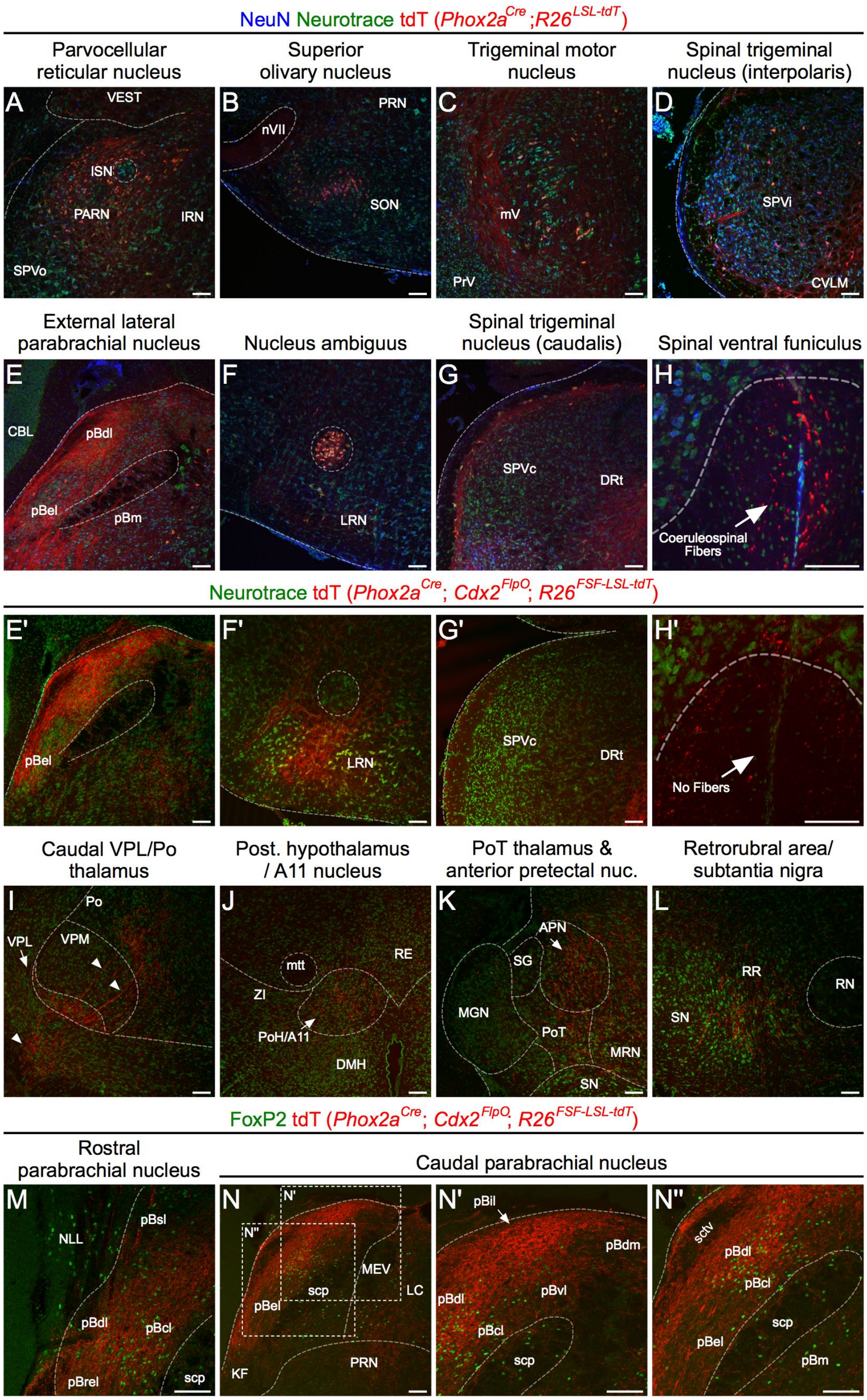


Roome et al., Figure S2

**Figure S2: Spinal Phox2a$^{Cre}$ neurons innervate AS targets. Related to Figure 2.**

(A–G) NeuN, Neurotrace and tdT staining in brain regions expressing *Phox2a*$^{Cre}$; *R26*$^{LSL-tdT/+}$ not depicted in Fig. 2. (E'–G') Lack of cellular tdT expression in *Phox2a*$^{Cre}$*; Cdx2*$^{FLPo}$*; R26*$^{FSF-LSL-tdT/+}$ mice compared to the same brain regions of *Phox2a*$^{Cre}$; *R26*$^{LSL-tdT/+}$ mice (E–G). (H, H') Large axons in the spinal ventral funiculus of *Phox2a*$^{Cre}$; *R26*$^{LSL-tdT/+}$ mice (H), absent in *Phox2a*$^{Cre}$*; Cdx2*$^{FLPo}$*; R26*$^{FSF-LSL-tdT/+}$ mice (H'), are likely coeruleospinal. (I–L) Additional brain regions receiving Phox2a$^{Cre}$+ spinofugal axons not depicted in Fig. 2. (M–N'') Spinal Phox2a$^{Cre}$+ axons in parabrachial subnuclei identified by FoxP2 expression (present in pBdl, pBcl but absent from pBel), in *Phox2a*$^{Cre}$*; Cdx2*$^{FLPo}$*; R26*$^{FSF-LSL-tdT/+}$ mice. N' and N'': higher magnification of boxes in N.

Numbers: (A–H) n=3 *Phox2a*$^{Cre}$; *R26*$^{LSL-tdT/+}$ adult mice, (E'–H', I–N, N', N'') n=3 *Phox2a*$^{Cre}$*; Cdx2*$^{FLPo}$*; R26*$^{FSF-LSL-tdT/+}$ adult mice.

Scale bars: 100 µm.

Abbreviations: A11 (dopaminergic cell group A11), APN (anterior pretectal nucleus), CBL (cerebellum), CVLM (caudal ventrolateral medulla), DMH (dorsomedial hypothalamus), DRt (dorsal reticular nucleus), IRN (intermediate reticular nucleus), ISN (inferior salivatory nucleus), KF (Kölliker-Fuse nucleus), LC (locus coeruleus), MEV (midbrain trigeminal nucleus), MGN (medial geniculate nucleus), MRN (midbrain reticular nucleus), mtt (mamillothalamic tract), mV (trigeminal motor nucleus), NLL (nucleus of the lateral lemniscus), nVII (facial motor nerve), PARN (parvocellular reticular nucleus), pBcl (central-lateral parabrachial nucleus), pBdl (dorsal-lateral parabrachial nucleus), pBdm (dorsal-medial parabrachial nucleus), pBel (external-lateral parabrachial nucleus), pBil (internal-lateral parabrachial nucleus), pBrel (rostral external-lateral parabrachial nucleus), pBsl (superior-lateral parabrachial nucleus), pBvl (ventral-lateral parabrachial nucleus), pBm (medial parabrachial nucleus), Po (posterior thalamus), PoH (posterior hypothalamus), PoT (posterior triangular thalamus), PRN (pontine reticular nucleus), RE (Reuniens nucleus), RN (red nucleus), RR (retrorubral nucleus), scp (superior cerebellar peduncle), sctv (ventral spinocerebellar tract), SG (suprageniculate nucleus), SN (substantia nigra), SON (superior olivary nucleus), SPVc (spinal trigeminal nucleus, caudalis), SPVi (spinal trigeminal nucleus, interpolaris), SPVo (spinal trigeminal nucleus, oralis), VEST (vestibular nuclei), VPL (ventral posterolateral thalamus), VPM (ventral posteromedial thalamus), ZI (zona incerta).

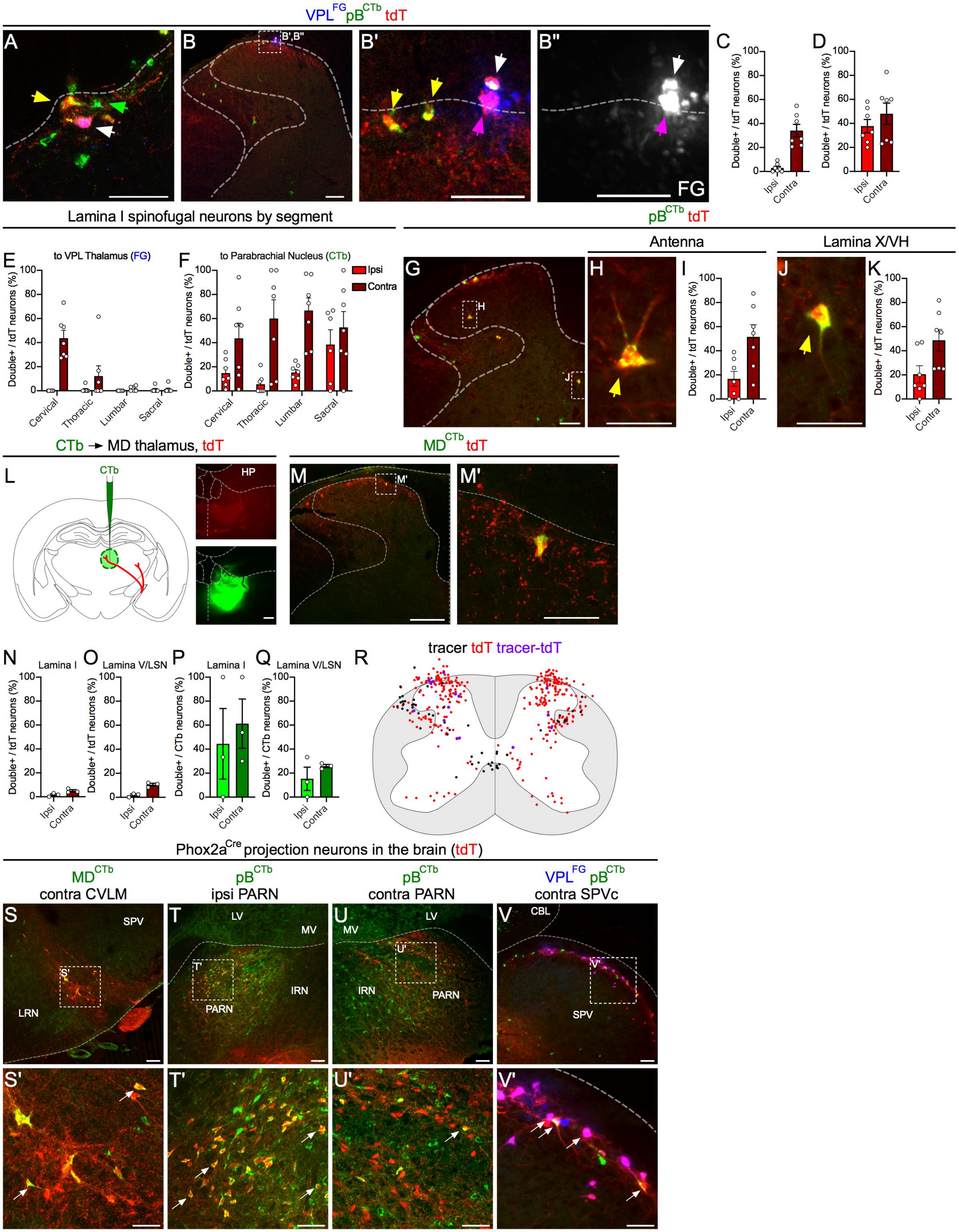


Roome et al., Figure S3

**Figure S3: Spinal Phox2a$^{Cre}$ neurons are predominantly AS neurons. Related to Figure 3.**

(A) Inset of Fig. 3G, demonstrating tdT and retrograde tracer co-localization in lamina I of *Phox2a$^{Cre}$*; *R26$^{LSL\text{-}tdT/+}$* mice with VPL-injected FG and pB-injected CTb. White arrow: tdT+ cell labelled with both FG and CTb; green arrow: CTb-only cell; yellow arrow: tdT+ cell labelled with CTb. (B–B'') Examples of FG-labelled neurons. As FG+ cells were difficult to detect with the confocal microscope, an epifluorescence microscope was used to photograph them prior to cover-slipping (B'') after which, CTb and tdT were imaged using a confocal microscope. FG and CTb/tdT images were then overlaid in register and merged; (B', B'') are higher magnifications of boxed lamina I area in B. White arrow: tdT+ cell doubly labelled with FG and CTb; magenta arrow: tdT+ cell labelled with FG; yellow arrow: tdT+ cell labelled with CTb. (C, D) Percent of cervical spinal cord tdT+ neurons labelled with FG (C) or CTb (D) injections. (E) Percent of tdT+ neurons labelled with FG (E) or CTb (F) in different spinal regions. (G–K) Depiction of rare Phox2a$^{Cre}$ neuron types such as antenna neurons (H) and lamina X/VH neurons (J). Percent of tdT+ neurons labeled with either or both tracers in Antenna (I) or lamina X/VH (K) neurons. (L–R) Analysis of *Phox2a$^{Cre}$*; *R26$^{LSL\text{-}tdT/+}$* mice injected in the MD thalamus with CTb. (L) Representative image of a CTb injection site. HP: Hippocampus (M, M') Representative cervical spinal cord section with a lamina I tdT+ neuron labelled with CTb (M'). (N–R) Percent of tdT neurons labelled with CTb in lamina I (N) or lamina V/LSN (O) at different spinal levels. (P–Q) Percent of CTb-labelled neurons expressing tdT in lamina I (P) or lamina V/LSN (Q) of all spinal segments. (R) Diagram of the location of tdT+ only (red), CTb only (black) or tdT+ and CTb-labelled (purple) neurons, in 5 non-sequential 25 μm sections of the cervical spinal cord compiled from all 3 animals analysed. (S–V) Examples of tdT+ neurons in various brain regions (arrows), labelled by retrograde tracer injections in the pB, VPL and MD thalamus. (S, S') tdT+ caudal ventrolateral medulla (CVLM) neurons project to the MD Thalamus, (T, T') tdT+ parvocellular reticular nucleus (PARN) neurons project to the ipsilateral parabrachial nucleus (U, U') but rarely to the contralateral parabrachial nucleus, and lamina I/paratrigeminal neurons of the spinal trigeminal nucleus (SPVc) project to the VPL thalamus and, much more sparsely, to the parabrachial nucleus (V, V').

Data are represented as mean ± SEM.

Numbers: (S4A–K, T–V) n=7 *Phox2a*$^{Cre}$; *R26*$^{LSL\text{-}tdT/+}$ adult mice (4 male, 3 female), (S4L–S) n=3 *Phox2a*$^{Cre}$; *R26*$^{LSL\text{-}tdT/+}$ adult mice.

Scale bars: 100 μm, except (A, B', B'', H, J, N, S'–V') 50 μm.

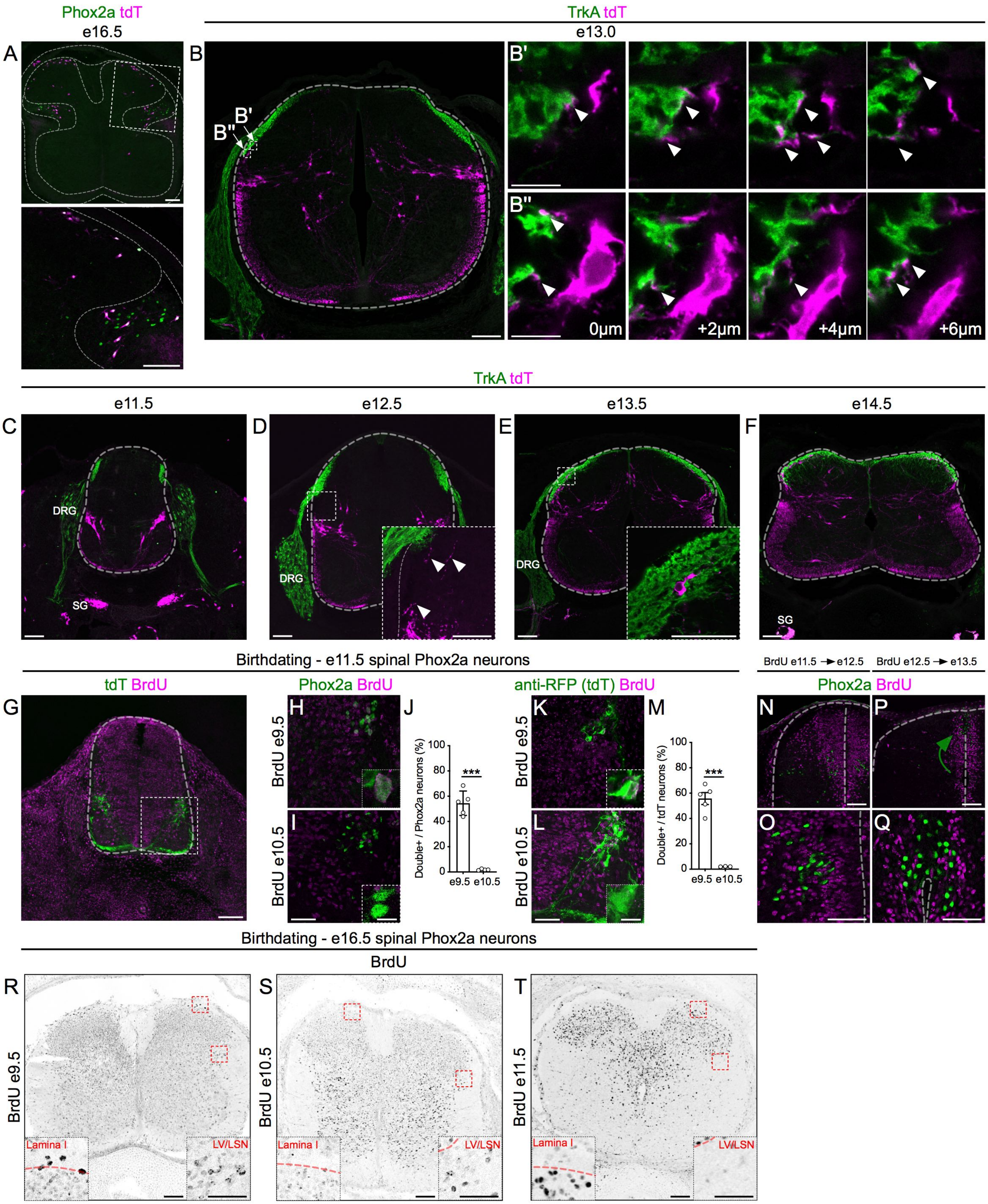


Roome et al., Figure S4

**Figure S4: Heterogeneity of spinal Phox2a neuron migration, sensory afferent interaction and birth time. Related to Figure 4.**

(A) Position of Phox2a+ (green) , tdT+ (magenta) and Phox2a+ tdT+ (white) neurons in e16.5 embryonic spinal cords of $Phox2a^{Cre}$; $R26^{LSL\text{-}tdT/+}$ mice. (B–F) Development of contacts between lamina I Phox2a neurons (magenta) and TrkA+ primary afferents (green) in e13.0 (B), e11.5 (C), e12.5 (D), e13.5 (E) and e14.5 (F) $Phox2a^{Cre}$; $R26^{LSL\text{-}tdT/+}$ mouse spinal cords. (B',B'') Consecutive confocal microscopy z-stack images of lamina I neuron (magenta) contacts with TrkA primary afferents (green) in e13.0 $Phox2a^{Cre}$; $R26^{LSL\text{-}tdT/+}$ mouse embryos, magnified boxed areas from (B). (D) inset: Phox2a neuron fibres near the dorsal root entry zone, prior to Phox2a cell soma entry into the dorsal horn. (E) inset: a lamina I Phox2a cell surrounded by TrkA afferents. (G–M) Birthdating of spinal Phox2a neurons in e11.5 $Phox2a^{Cre}$; $R26^{LSL\text{-}tdT/+}$ mouse embryos, labelled by BrdU exposure at e9.5 (H, K) or e10.5 (I, L). (G) The spinal cord of an e11.5 embryo exposed to BrdU at e10.5, showing BrdU (magenta) and RFP (tdT, green) labelling. (J, M) Percent of all tdT cells labelled with BrdU (J) and the percent of all Phox2a+ cells labelled with BrdU (M), in e11.5 $Phox2a^{Cre}$; $R26^{LSL\text{-}tdT/+}$ embryos exposed to BrdU at e9.5 or e10.5 (n=4-5). (N, O) e12.5 spinal cord of $Phox2a^{Cre}$; $R26^{LSL\text{-}tdT/+}$ embryos showing $Phox2a^{DeepLate}$ neurons (Phox2a+, green) intermingled with newly born (BrdU+, magenta) neurons exposed to BrdU at e11.5. (P, Q) e13.5 spinal cord of $Phox2a^{Cre}$; $R26^{LSL\text{-}tdT/+}$ mouse embryos with $Phox2a^{DeepLate}$ neurons (green) neurons clustered near the roof of the central canal, separate from other newborn (BrdU+, magenta) neurons, labelled with BrdU at e12.5. (R–T) e16.5 spinal cord of $Phox2a^{Cre}$; $R26^{LSL\text{-}tdT/+}$ mouse embryos pulsed with BrdU at e9.5 (R), e10.5 (S) or e11.5 (T), showing BrdU+ cells (black) in lamina I and lamina V/LSN (insets). Lamina I cells are labelled by a BrdU pulse at e9.5 but e10.5 and e11.5 BrdU pulses result in sparser labelling. Lamina V/LSN cells show labelling by e9.5 and e10.5 BrdU pulses but minimal labelling by a e11.5 BrdU pulse.

Data are represented as mean ± SEM.

Numbers: *Phox2a*$^{Cre}$; *R26*$^{LSL\text{-}tdT/+}$ embryos: (A) n=3 e16.5, (B) n=3 e13.0, (C) n=3 e11.5, (D) n=3 e12.5, (E) n=3 e13.5, (F) n=3 e14.5, (G–M) n=3-5 e11.5 per condition, (N, O) n=3 e12.5, (P, Q) n=3 e13.5, (R) n=3 e16.5, (S) n=3 e16.5, (T) n=3 e16.5.

Statistics: (J, M) Unpaired t-test, ***: p<0.0001.

Scale bars: (A–G, N, P, R–T) 100 µm. (H, I, K, L, O, Q) and insets of (D, E, R, S, T) 50 µm, (B’, B’’) and insets of (H, I, K, L) 10 µm.

Abbreviations: SG: sympathetic ganglion; DRG: Dorsal root ganglion.

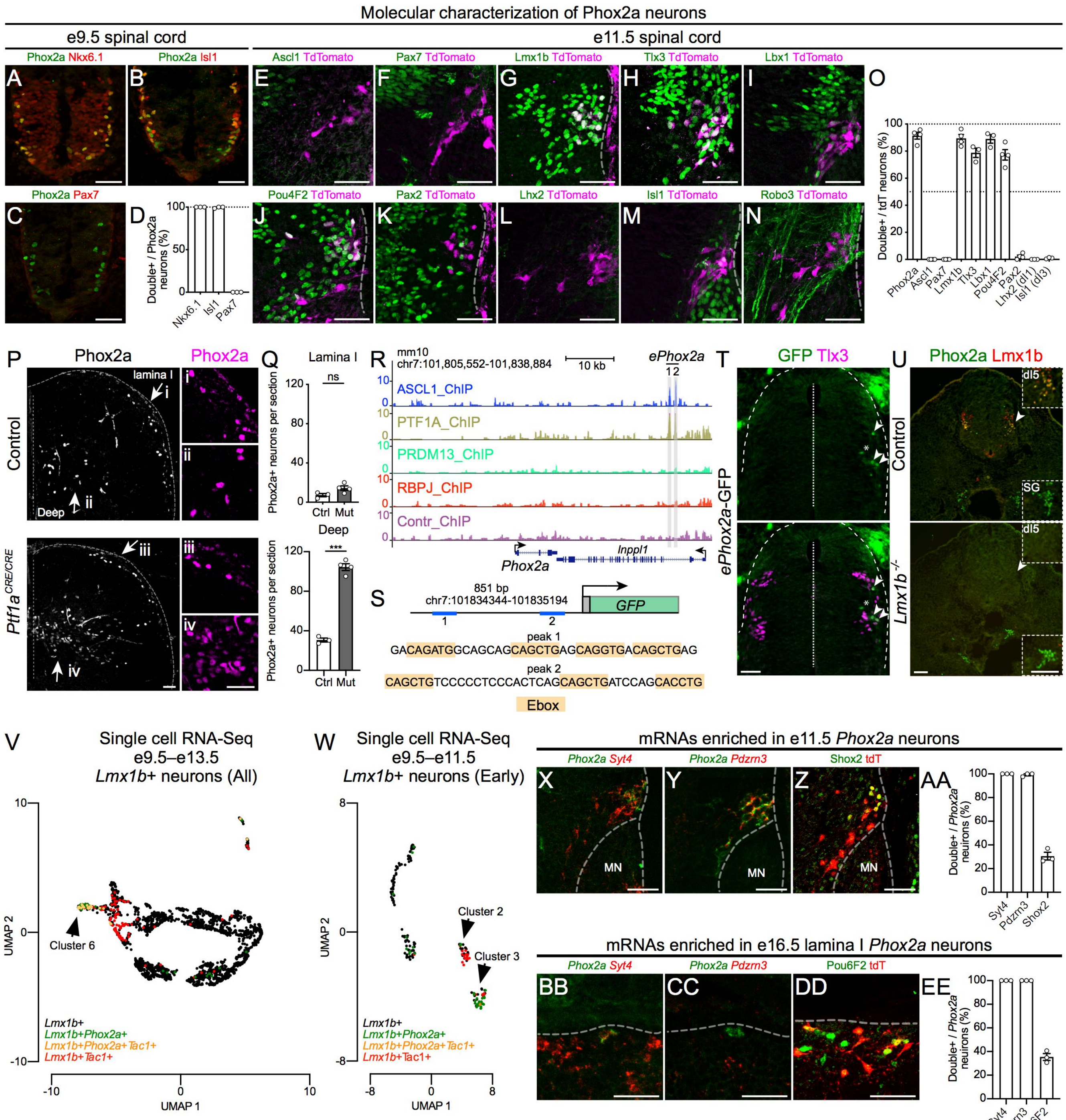


Roome et al., Figure S5

**Figure S5: The molecular identity and specification of spinal Phox2a neurons. Related to Figure 5.**

(A–D) Molecular characterisation of spinal Phox2a neurons in the e9.5 *Phox2a*$^{Cre}$; *R26*$^{LSL\text{-}tdT/+}$ spinal cord. Phox2a neurons co-express ventral motor neuron markers Nkx6.1 (A), Isl1 (B), but not the progenitor marker Pax7 (C), and are most likely accessory motor neurons. (D) Percent of Phox2a+ cells co-expressing Nkx6.1, Isl1 and Pax7. (E–O) Molecular characterization of spinal tdT+ neurons in the e11.5 *Phox2a*$^{Cre}$; *R26*$^{LSL\text{-}tdT/+}$ spinal cord. (E, F) tdT+ neurons do not express Ascl1 (E) and Pax7 (F); (G–N) TdT+ neuron expression of dorsal interneuron markers Lmx1b (G), Tlx3 (H), Lbx1 (I), Pou4F2 (J), Pax2 (K), Lhx2 (L), Isl1 (M), and commissural neuron marker Robo3 (N). (O) Percent of tdT+ cells expressing the markers visualised in (E–N). (P) Detection of Phox2a cells in spinal cords of control and *Ptf1a*$^{Cre/Cre}$ e14.5 mice. Panels with Roman numerals are magnifications of regions marked by arrows. (Q) Number of Phox2a+ cells per average section in lamina I and deep laminae of e14.5 control and *Ptf1a*$^{Cre/Cre}$ mice. (R, S) Genome sequences ChIP-Seq hits detected using antibodies against Ascl1, Ptf1a, Prdm13, and Rbpj at the *Phox2a* locus (R). The two sites bound by both Ascl1 and Ptf1a are highlighted in gray. (S) Diagram of the *ePhox2a* GFP reporter construct with the cis-regulatory sequence containing Ascl1 and Ptf1a sites. The sequence at the center of each ChIP-seq peak is shown with the E-box bHLH transcription factors binding motif is highlighted in orange. The active Ptf1a containing transcription complex requires a TC-Rbpj binding sequence that is not present in this enhancer. (T) Transverse sections of embryonic day 4 (E4) chicken neural tubes electroporated with the *ePhox2a-GFP* reporter at embryonic day 2. *ePhox2a* directs dI5-specific GFP expression in Tlx3+ cells. (U) e11.5 *Lmx1b*$^{-/-}$ spinal cords lack Phox2a expression (dI5 inset), but sympathetic ganglia maintain it (SG inset). (V–W) UMAP plots of e9.5-e13.5 *Lmx1b*+ neurons (V) and e9.5-e11.5 *Lmx1b*+ neurons (W) showing *Lmx1b*+/*Phox2a*+, *Lmx1b*+/*Tac1*+, and *Lmx1b*+/*Phox2a*+/*Tac1*+ cell clusters. (X–AA) Validation of selected dI5-enriched mRNAs at e11.5 by RNA-Scope in situ detection of *Synaptotagmin 4* (X), *Pdzrn3* (Y) mRNAs and Shox2 protein immunofluorescence (Z). (AA) Percent of e11.5 Phox2a+ or *Phox2a* mRNA+ neurons co-expressing the selected mRNAs or proteins. (BB–EE) Validation of selected dI5-enriched mRNAs at e16.5 in lamina I, by RNA-Scope in situ detection of *Synaptotagmin 4* (BB) and *Pdzrn3* mRNAs (CC) and Pou6F2 protein immunofluorescence (DD). (EE) Percent of e16.5 Phox2a+ or *Phox2a* mRNA+ neurons co-expressing the selected mRNAs or proteins.

Data are represented as mean ± SEM.

Numbers: *Phox2a*$^{Cre}$; *R26*$^{LSL\text{-}tdT/+}$ embryos (A–D) n=3 e9.5, (E–O) n=3-4 e11.5, (X–AA) n=3 e11.5, (BB–EE) n=3 e16.5. (P–Q) n=3 control, n=4 *Ptf1a*$^{Cre/Cre}$ e14.5 embryos, (T) n=6 E4 chicken embryos, (U) n=3 control, n=3 Lmx1b$^{-/-}$ e11.5 embryos. Statistics: (P, Q) Groups compared with Student's t-test, ***: p<0.001. (V, W) Data derived from Delile et al., (2019); data processing and statistics described in STAR Methods.

Scale bars: All 50 µm, except (U) 100 µm.

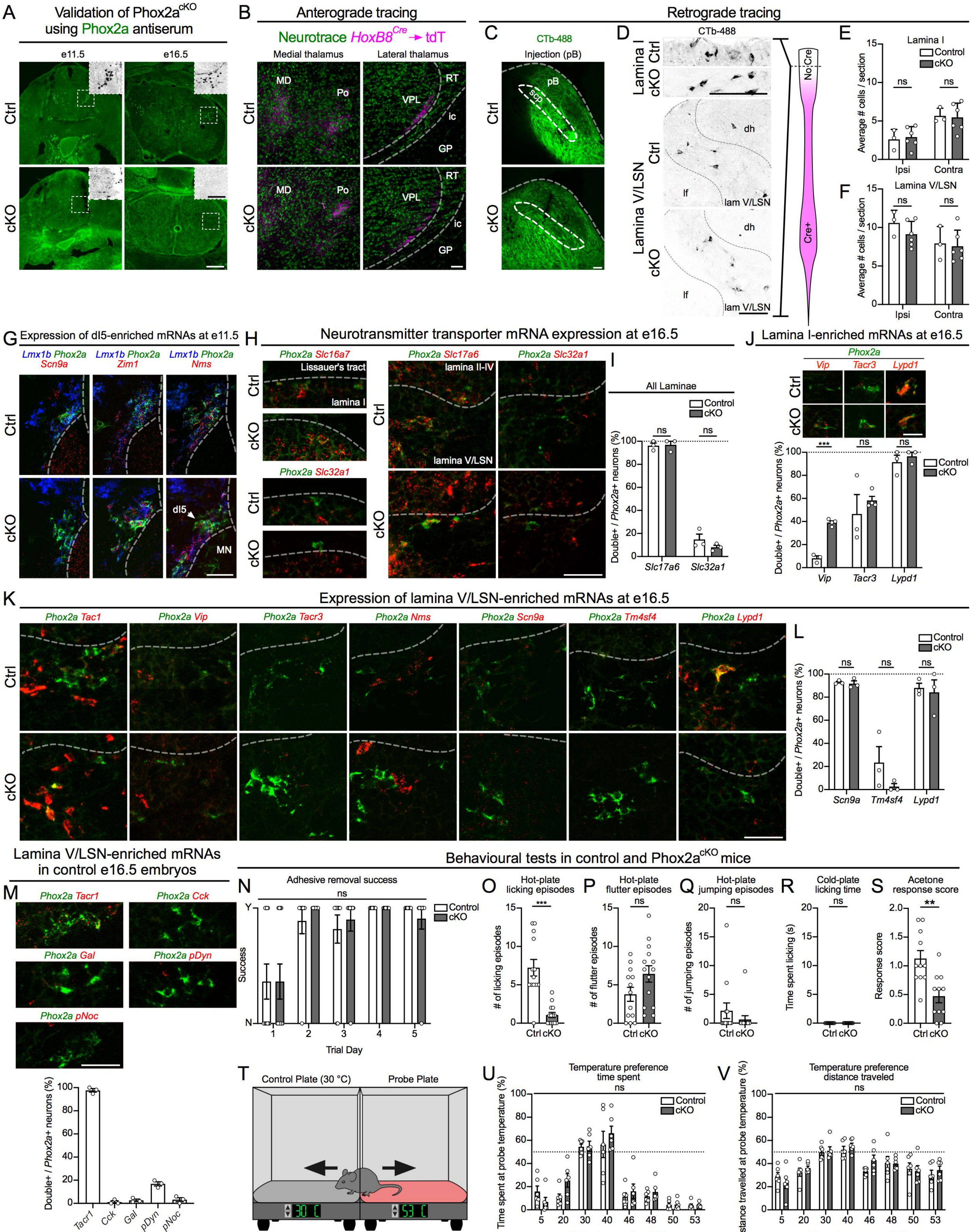


Roome et al., Figure S6

**Figure S6: Phox2a is required for AS neuron development and function. Related to Figure 6.**

(A) Phox2a expression (green) in e11.5 (left) and e16.5 (right), control (top row, Ctrl) and Phox2a$^{cKO}$ (bottom row; cKO) spinal cords. Insets are magnified boxed areas, depicting the loss of Phox2a expression (black) in cKO. (B) Thalamus of *HoxB8*$^{Cre}$; *Phox2a*$^{+/+}$; *R26*$^{LSL\text{-}tdT/+}$ (Ctrl, top row) and *HoxB8*$^{Cre}$; *Phox2a*$^{f/f}$; *R26*$^{LSL\text{-}tdT/+}$ (Phox2a$^{cKO}$ or cKO, bottom row) mice, in which spino-thalamic axons are labelled via *HoxB8*$^{Cre}$-driven axonal tdT (magenta), and are found in the medial (left) or lateral (right) thalamus, stained with Neurotrace in green. (C) Representative CTb injection site in the pB of control and Phox2a$^{cKO}$ adult mice. (D–F) CTb-labeled neurons in lamina I (upper panels, quantified in E) and lamina V/LSN (lower panels, quantified in F), in the cervical spinal cord of control and Phox2a$^{cKO}$ mice, rostral to HoxB8$^{Cre}$ expression domain, diagrammed in magenta. (G) Expression of dI5-enriched mRNAs in e11.5 control (top row) and Phox2a$^{cKO}$ (bottom row) spinal cords not shown in Fig. 6 but quantified in Fig. 6G. (H) Expression of *Slc17a6* and *Slc32a1* mRNAs encoding, respectively, the neurotransmitter transporters vGlut2 and vGAT, in lamina I and lamina V/LSN of e16.5 control and Phox2a$^{cKO}$ embryos. (I) Percent of control and Phox2a$^{cKO}$ *Phox2a*+ neurons expressing *Slc17a6* and *Slc32a1* mRNAs (n=3 control, n=3 Phox2a$^{cKO}$). (J) Expression and quantification of candidate AS-enriched mRNAs in lamina I *Phox2a*+_neurons of e16.5 control and Phox2a$^{cKO}$ embryos (n=3 control, n=3 Phox2a$^{cKO}$). (K) Expression of mRNAs encoding neuropeptides and neuropeptide receptors in lamina V/LSN *Phox2a*+ neurons of e16.5 control (top row) and Phox2a$^{cKO}$ (bottom row) spinal cords not shown in Fig. 6 but quantified in Fig. 6M. (L) Expression and quantification of mRNAs encoding neuropeptides expressed in lamina V/LSN *Phox2a*+ neurons, not depicted in Fig. 6L. (M) Expression and quantification of mRNAs encoding neuropeptides and neuropeptide receptors in *Phox2a*+ neurons of control e16.5 embryos. (N) Adhesive removal success ratio between control and Phox2a$^{cKO}$ mice over 5 days. (O–S) Responses to non-injected noxious stimuli: episodes of (O) licking, (P) hind paw flutter, and (Q) jumping during the hot-plate test (n=13 control, n=14 Phox2a$^{cKO}$). (R) Time spent licking during the cold-plate test (n=9 control, n=10 Phox2a$^{cKO}$). (S) The acetone response score after hind paw application of acetone (n=6 control, n=6 Phox2a$^{cKO}$). (T–V) The two choice temperature preference assay (T), demonstrating time spent at the probe temperature versus 30 °C (U), and the distance traveled in the probe temperature compartment versus 30 °C (V) (n=6 control, n=6 Phox2a$^{cKO}$).

Data are represented as mean ± SEM.

Numbers: (A) n=3 control, n=3 Phox2a$^{cKO}$ e11.5 and e16.5 mice. (B) n=4 control, n=4 Phox2a$^{cKO}$ adult mice; (C–F) n=3 control, n=6 Phox2a$^{cKO}$ adult mice; (G) n=3 control, n=5 Phox2a$^{cKO}$ e11.5 mice; (H, I) n=3 control, n=3 Phox2a$^{cKO}$ e16.5 mice; (J) n=3 control, n=4 Phox2a$^{cKO}$ e16.5 mice; (K) n=3 control, n=3 Phox2a$^{cKO}$ e16.5 mice; (L) n=3 control e16.5 mice; (N–V) Numbers described above.

Statistics: (E, F) Two-way ANOVA with Tukey's multiple comparisons test, (I, J, K) multiple t-tests using Holm-Sidak method, (M) mixed-effects analysis with Sidak's multiple comparisons test, (O, P, Q, R, S) Mann Whitney test, and (U, V) two-way ANOVA with Sidak's multiple comparisons test. ns: non-significant, **: $p<0.01$, ***: $p<0.001$.

Scale bars: (A) 200 μm, insets in (A) and (B–D) 100 μm, (G, H, K, M) 50 μm and (J) 25 μm.

Images in Fig. 5BB, EE, FF have been re-used in Fig. S6G. Validation of enriched mRNAs from RNA-Seq analysis in control embryos (Fig. 5) and comparison of enriched mRNAs between control and Phox2a$^{cKO}$ embryos (Fig. S6) were performed as a single experiment.

Abbreviations: dh (dorsal horn), DRG (dorsal root ganglion), ic (internal capsule), GP (globus pallidus), lf (lateral funiculus), MD (mediodorsal thalamus), MN (motor neurons), pB (parabrachial nucleus), Po (posterior thalamus), scp (superior cerebellar peduncle), RT (reticular thalamus), VPL (ventroposterolateral thalamus).

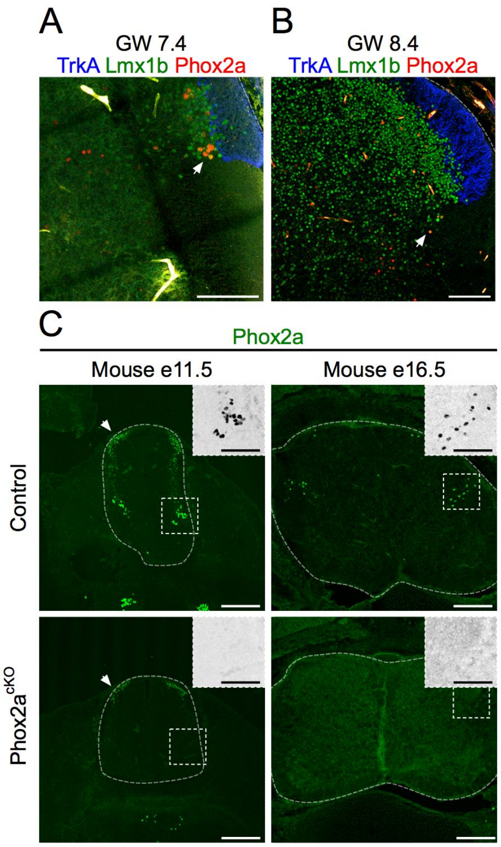


Roome et al., Figure S7

**Fig. S7: Phox2a neuron molecular identity is conserved in the developing human spinal cord. Related to Figure 7.**

(A, B) Insets from G.W 7.4 (A) and G.W 8.4 (B) human spinal cords in Fig. 7, demonstrating Phox2a and Lmx1b co-expression, and apposition of TrkA fibres with Phox2a neurons. (C) Validation of the Abcam Phox2a antibody in mouse Phox2a$^{cKO}$ tissue at both e11.5 and e16.5. Arrows show immunostaining in dorsal dI neurons, which was not observed with the Phox2a antibody used in previous experiments (from JF Brunet) or by RNA in situ detection of *Phox2a* mRNA, and it is not eliminated in the Phox2a$^{cKO}$.

Numbers: (A) Single G.W 7.4 human spinal cord, (B) single G.W 8.4 human spinal cord, (C) n=3 e11.5 control, n=3 e11.5 Phox2a$^{cKO}$, n=3 e16.5 control, n=3 e16.5 Phox2a$^{cKO}$ mouse embryos.

Scale bars: (A, B) and insets in (C) 100 μm, (C) 200 μm.

**Tables:**

**Table S1: mRNAs enriched in embryonic AS neurons. Related to Figure 5.**

Top 25 enriched mRNAs per three clusters identified in Fig. 5W and 5Y, when compared to all other spinal neurons in the dataset at the same embryonic ages. For each cluster, the transcript name, log$_{10}$fold change (log$_{10}$FC) and -log$_{10}$p values are represented. P-values rounded to 0 (<2.22x10^-308) were recorded as having a -log$_{10}$P value of ">307.65". Transcripts with high -log10P values that were not expressed in non-Lmx1b+ neurons were considered to be useful markers for Phox2a neurons.

| e9.5-e13.5 *Lmx1b* Cluster 6 | | | e9.5-e11.5 *Lmx1b* Cluster 2 | | | e9.5-e11.5 *Lmx1b* Cluster 3 | | |
|---|---|---|---|---|---|---|---|---|
| **Name** | **log$_{10}$FC** | **-log$_{10}$p** | **Name** | **log$_{10}$FC** | **-log$_{10}$p** | **Name** | **log$_{10}$FC** | **-log$_{10}$p** |
| *Phox2a* | 0.97 | >307.65 | *Lmx1b* | 0.75 | 218.80 | *Lmx1b* | 0.74 | 270.53 |
| *Nms* | 0.44 | >307.65 | *Tac1* | 1.16 | 94.60 | *Nms* | 0.65 | 177.44 |
| *Syt13* | 0.66 | 271.21 | *C130021* | 0.53 | 48.54 | *Phox2a* | 1.11 | 158.15 |
| *Zim1* | 0.20 | 176.74 | *Pou4f2* | 0.45 | 37.76 | *C130021I* | 0.67 | 99.06 |
| *6330403* | 0.28 | 146.78 | *Gabra2* | 0.50 | 29.19 | *Car10* | 0.64 | 88.30 |
| *Arhgdig* | 0.66 | 138.31 | *Tlx3* | 0.60 | 24.04 | *Tm4sf4* | 0.14 | 86.40 |
| *Snca* | 1.01 | 123.62 | *Syt13* | 0.42 | 18.00 | *Syt13* | 0.66 | 63.69 |
| *Syt4* | 2.18 | 120.03 | *Chl1* | 0.86 | 17.43 | *Mar1* | 0.49 | 50.42 |
| *Lmx1b* | 0.56 | 119.42 | *Tcerg1l* | 0.51 | 16.68 | *Rit2* | 0.31 | 48.88 |
| *Scn9a* | 0.50 | 112.81 | *Mab21l2* | 1.07 | 16.43 | *Pde8b* | 0.14 | 46.44 |
| *Sema3a* | 0.44 | 107.43 | *Pou4f1* | 1.26 | 15.95 | *Fam19a2* | 0.47 | 46.09 |
| *Kctd4* | 0.33 | 103.94 | *Tmeff2* | 0.57 | 14.85 | *Tcerg1l* | 0.74 | 41.81 |
| *Sncg* | 1.77 | 103.89 | *Kif26b* | 0.36 | 13.68 | *Mdga1* | 0.41 | 40.92 |
| *Tac1* | 0.48 | 101.61 | *Cadm2* | 0.40 | 13.52 | *Pdzrn3* | 0.91 | 39.54 |
| *Rit2* | 0.45 | 101.60 | *Slc17a6* | 0.54 | 13.39 | *Nrxn1* | 1.26 | 38.57 |
| *Car10* | 0.50 | 99.29 | *Tshz2* | 1.03 | 12.96 | *Tmeff2* | 0.67 | 37.60 |
| *Pcp4* | 1.18 | 94.51 | *Mab21l1* | 0.42 | 12.76 | *Negr1* | 0.58 | 36.64 |
| *Tcerg1l* | 0.62 | 92.38 | *Ntrk3* | 0.50 | 12.00 | *Scn3b* | 0.65 | 31.99 |
| *Pde8b* | 0.12 | 91.23 | *Ppp3ca* | 0.78 | 11.09 | *Syt4* | 1.91 | 30.60 |
| *Tspan17* | 0.65 | 88.46 | *Nrxn1* | 0.77 | 10.56 | *Stk39* | 0.86 | 27.30 |
| *Scn3b* | 0.63 | 83.81 | *Nms* | 0.16 | 10.19 | *Shox2* | 0.85 | 26.70 |
| *Adap1* | 0.27 | 81.32 | *Mtus2* | 0.70 | 10.11 | *Stk32b* | 0.55 | 25.69 |
| *Fam19a* | 0.48 | 80.39 | *Scn3b* | 0.46 | 9.95 | *Chl1* | 0.88 | 25.24 |
| *Nptxr* | 0.28 | 76.58 | *Stk39* | 0.69 | 9.84 | *Sema3a* | 0.45 | 25.20 |
| *Gap43* | 1.91 | 76.21 | *Hs3st2* | 0.10 | 9.56 | *Necab2* | 0.50 | 24.31 |
